# Asymmetrical lineage introgression and recombination in populations of *Aspergillus flavus*: implications for biological control

**DOI:** 10.1101/2022.03.12.484001

**Authors:** Megan S. Molo, James B. White, Vicki Cornish, Richard M Gell, Oliver Baars, Rakhi Singh, Mary Anna Carbone, Thomas Isakeit, Kiersten A. Wise, Charles P. Woloshuk, Burton H. Bluhm, Bruce W. Horn, Ron W. Heiniger, Ignazio Carbone

## Abstract

*Aspergillus flavus* is an agriculturally important fungus that causes ear rot of maize and produces aflatoxins (AFs), of which B_1_ is the most potent carcinogen known. In the US, the management of AFs includes the deployment of biological control agents that comprise two nonaflatoxigenic *A. flavus* strains, either Afla-Guard® (member of lineage IB) or AF36 (lineage IC). We used genotyping-by-sequencing to examine the influence of both biocontrol agents on native populations of *A. flavus* in cornfields in Texas, North Carolina, Arkansas, and Indiana. This study examined up to 27,529 single-nucleotide polymorphisms (SNPs) in a total of 815 *A. flavus* isolates, and 353 genome-wide haplotypes sampled before biocontrol application, three months after biocontrol application, and up to three years after initial application. Here, we report that the two distinct *A. flavus* evolutionary lineages IB and IC differ significantly in their frequency distributions across states. We provide evidence of increased unidirectional gene flow from lineage IB into IC, inferred to be due to the applied Afla-Guard biocontrol strain. Genetic exchange and recombination of biocontrol strains with native strains was detected in as little as three months after biocontrol application and up to one and three years later. There was limited inter-lineage migration in the untreated fields. These findings suggest that biocontrol products that include strains from lineage IB offer the greatest potential for sustained reductions in aflatoxin levels over several years. This knowledge has important implications for developing new biocontrol strategies.

## INTRODUCTION

Many fungi produce mycotoxins (secondary metabolites) with negative health impacts on humans and animals (Pitt 2000; Sweeney & Dobson 1998). An agriculturally important mycotoxin producer is *Aspergillus flavus*, which causes ear rot of maize and produces aflatoxins (AFs). AFs are polyketide secondary metabolites produced by *A. flavus* and several other species in *Aspergillus* section *Flavi. A. flavus* is a soilborne filamentous fungus that commonly contaminates many economically important crops, such as corn, peanuts, cotton, tree nuts, and spices notably curry and chili (Makhlouf *et al*. 2019), with AFs (Bennett & Klich 2003; Eaton & Groopman 1994; Horn 2007; Squire 1981; Wu 2004). In addition to AFs, *A. flavus* also produces cyclopiazonic acid (CPA), an indole- tetramic acid that targets the liver, kidneys, and gastrointestinal tract of animals (Burdock & Flamm 2000; Moore *et al*. 2009; Vonberg & Gastmeier 2006). Because of the adverse effects on human and animal health, the US Food and Drug Administration (FDA) strictly regulate the levels of AFs in grain. Grains must yield levels of AF below 20 parts per billion (ppb) for human consumption and 100 ppb for animal feed (FDA 1979). Crops exceeding these limits are rejected resulting in millions of dollars in losses (Wu & Guclu 2012).

Current management of AFs includes the application of nonaflatoxigenic *A. flavus* biological control agents, either Afla-Guard® (active ingredient=NRRL 21882) or AF36 Prevail (formerly AF36, active ingredient=NRRL 18543), that competitively exclude native aflatoxigenic *A. flavus* strains for space and resources in corn ears (Dorner 2005; Moore *et al*. 2009). Biocontrol strategies have been shown to effectively reduce AF levels by approximately 70-90% (Dorner 2004; Dorner 2008; Pitt & Hocking 2006; Yin *et al*. 2008). Afla-Guard® was registered with the EPA in 2004 for reduction of AF contamination in peanuts and corn in the United States (EPA 2004). *A. flavus* NRRL 21882 is nonaflatoxigenic and missing the entire AF and CPA gene clusters (Chang *et al*. 2005; Moore *et al*. 2009). AF36 was registered with the U.S. Environmental Protection Agency (EPA) in 2003 for use on cotton and corn in Arizona and Texas (EPA 2003). A nonsense mutation in the *pksA* (*aflC*) gene early in the AF biosynthetic pathway renders *A. flavus* NRRL 18543 nonaflatoxigenic (Ehrlich & Cotty 2004; Moore *et al*. 2009). Moreover, NRRL 18543 produces CPA and its application has been shown to significantly increase CPA contamination in grain (Abbas *et al*. 2011b). While CPA is currently unregulated by the FDA, it has been linked to adverse health effects in humans and animals (Burdock & Flamm 2000).

Aflatoxin production is a polygenic trait and influenced by multiple environmental factors that also play a role in crop health, such as elevated temperature and water stress (Medina *et al*. 2014), insect damage (Mc Millian *et al*. 1985), soil nitrogen (Anderson *et al*. 1975; Jones & Duncan 1981; Payne 1989), pH (Ehrlich *et al*. 2005) and carbon availability (Lasram *et al*. 2016). The persistence of mycotoxins in field populations also has a genetic basis, and both aflatoxin and CPA clusters show high heritability from parents to progeny strains (Olarte *et al*. 2012; Olarte *et al*. 2015).

Aflatoxin biosynthesis is controlled by enzymes encoded by approximately 30 genes located in the 75-kb subtelomeric region of chromosome 3 (Carbone *et al*. 2007).

Sequence polymorphisms and deletions in the aflatoxin gene cluster as well as genome- wide variation separate *A. flavus* strains into two distinct evolutionary lineages, IB and IC (Moore *et al*. 2009; Olarte *et al*. 2012). Lineage IB is composed of strains with partial and full cluster deletions and strains with full clusters with many fixed polymorphisms (Moore *et al*. 2009), resulting primarily in strains that are nonaflatoxigenic or strains producing very low amounts of AF (Geiser *et al*. 2000; Moore *et al*. 2017). Lineage IC includes predominantly full cluster strains that are aflatoxigenic and harbor extensive quantitative variation in aflatoxin production, from very high AF producers to nonproducers (Moore *et al*. 2009).

Populations of *A. flavus* have a clonal and recombining population structure (Drott *et al*. 2020; Geiser *et al*. 1998; Moore *et al*. 2013; Moore *et al*. 2017; Moore *et al*. 2009; Olarte *et al*. 2012), and frequently maintain a mix of aflatoxigenic and nonaflatoxigenic strains (Chang *et al*. 2005; Cotty & Bhatnagar 1994; Drott *et al*. 2020). Olarte *et. al* (2012) provided the first direct evidence of sexual recombination and identified chromosome crossover events in parents that influence mycotoxin phenotypes of *A. flavus* progeny strains. *A. flavus* is heterothallic and sexual reproduction occurs between isolates of opposite mating types, *MAT1-1* and *MAT1-2*, that belong to different vegetative compatibility groups (VCGs) (Horn *et al*. 2009; Ramirez-Prado *et al*. 2008).

Previous research (Horn *et al*. 2016) reported that the degree of fertility in *A. flavus* was strongly influenced by the parental origin of the sclerotia, whereby one combination of sclerotia-conidia was highly fertile and the reciprocal combination exhibited low fertility. Since there is also a demonstrated level of fertility among inter-lineage (Horn *et al*. 2009; Olarte *et al*. 2012) and interspecific (Olarte *et al*. 2015) crosses, we cannot rule out genetic exchange and recombination among closely related lineages and sibling species impacting current estimates of genetic diversity in *A. flavus*. Genetic exchange is also possible between compatible strains via heterokaryon formation within a VCG, a process known as parasexuality (Papa 1973). In filamentous fungi, the vegetative compatibility system is a self/non-self recognition system controlled by a series of heterokaryon *(het)* incompatibility loci (Leslie 1993). Heterokaryon incompatibility is the inability of two strains to undergo fusion of vegetative fungal cells. Twelve putative *het* loci have been identified in *A. flavus* and in most cases alleles must be identical at all *het* loci for stable hyphal fusions to occur (Monacell 2014). Fungal individuals can be grouped into VCGs based on their multilocus genotypes, provided that marker loci used for genotyping are in linkage disequilibrium with *het* loci on the same chromosomal segment (Monacell 2014).

Several studies report a high turnover of VCGs in the same regions from year to year (Atehnkeng *et al*. 2014; Bayman & Cotty 1991) and in progeny strains when compared to parental VCGs after a single generation of sex (Olarte *et al*. 2012). This is an important observation because each VCG is a clone and all isolates within a VCG have a similar toxin profile (Ehrlich & Cotty 2004). Therefore, the observed diversity in aflatoxin concentrations in field populations is largely driven by sexual reproduction increasing VCG diversity (Horn & Dorner 1999; Olarte *et al*. 2012). Laboratory and field experiments indicate that a single round of sex can significantly increase genetic diversity (Horn *et al*. 2016; Olarte *et al*. 2012); however, the impact that sexual reproduction has in structuring natural populations of these fungi over time has not been studied. Although there is some yearly carry over of biocontrol agents, current management practices recommend reapplication of biocontrol agents each growing season because presumably their population sizes decline, but their influence on the native population structure is unknown. The lack of predictive models for aflatoxin contamination and high cost of application deters growers in moderate to low-risk areas from applying the biocontrol products. Improving on biocontrol to make it more sustainable is attractive for economical and practical reasons; however, this requires better knowledge of the underlying genetic and environmental processes that influence biocontrol. This suggests that a better understanding of *A. flavus* population genetics over time and space is key to developing more sustainable and predictable approaches to manage mycotoxin contamination of crops. Here we adopt a population genomics and phylogenetic approach to explore the effect of released biocontrol strains on native populations in cornfields, for up to three years after initial application.

## Materials and Methods

### Sampling and DNA Isolation

In 2013 and 2014 *A. flavus* was sampled from cornfields in Texas (Texas A&M University Farm, Burleson County, TX; 30.5472° -96.4297°), North Carolina (Upper Coastal Plain Research Station, Fountain Farm, Rocky Mount, Edgecombe County, NC; 35.8937° -77.6811°), Arkansas (Newport Research Station, Newport, Jackson County, AR; 35.5714° -91.2602°), and Indiana (Southeast-Purdue Agricultural Center, Jennings County, IN; 39.0334°, -85.5258°). Prior to our study, these maize fields had no history of biological control application. Fields were separated into twelve plots with eight 30-ft rows in each plot with three treatments: untreated control, Afla-Guard, and AF36, replicated four times. Best management practices for irrigation and fertilization were implemented by location. Corn hybrid selection, fertility and weed management programs also followed local recommendations. The selected corn hybrids were BT- traited and herbicide-tolerant, if appropriate, and fields were under a no-till, no crop rotation system. Commercial biocontrol inoculum was used, using either sterilized wheat kernels (AF36) or dehulled barley (Afla-Guard) as the carrier/nutrient source for dormant *A. flavus* propagules. Biocontrols were applied over the top of the corn crop at tasseling (VT corn stage) at a rate of 20 lb/acre. All treatments and samples were applied and taken from the middle of rows 3 and 4 of each plot to allow for a 100-ft buffer between plots.

Soil samples were collected by pooling 20, 10-cc sub-samples from each of the two rows of the plot, and *A. flavus* was isolated from each plot sample at three different time points: before biocontrol treatment (pre-application), three months after biocontrol application (post 3-months), and one year after application (post 1-year). *A. flavus* was isolated from soil samples by dilution plating onto modified dicloran rose Bengal agar, as described previously (Horn & Greene 1995). At crop maturity, eight ears were collected from each plot, shelled separately, and 100 kernel sub-samples from each ear were surface disinfested in 10% bleach, followed by two rinses in sterile water. The 100 kernel samples were evenly spaced onto moistened sterile paper towels in 20 cm × 20 cm sterile aluminum pans. The pans were placed in Ziploc bags and incubated at ambient room temperature for one week. Colonies of *A. flavus* from kernels were subcultured onto Czapek-Dox agar.

To examine more long-term changes in population genetic structure after initial biocontrol application we focused on four commercial dryland cornfields in Burleson County, Texas. Fields A (Ships clay, 30.5487° -96.4275°) and C (Weswood silt loam, 30.5398° -96.4165°) had never received a biocontrol treatment, while fields B (Weswood silty clay loam, 30.5545° -96.4243°) and D (Weswood silt loam, 30.5392° -96.4159°) were treated with 20 lb/acre of Afla-Guard in May 2011. Since 2011, fields A, C, and D were continually planted with corn, while in field B was rotated with wheat in 2013 prior to planting with corn in 2014. Because aflatoxin assays in these fields consistently showed aflatoxin levels well below 100 ppb over several years, *A. flavus* was isolated in 2014 from soil samples and surface-disinfested kernels obtained from individual ears, as described previously, and included in population genomic analyses to better understand how the underlying lineage composition and population genetic structure contribute to overall lower aflatoxin levels.

DNA was isolated from *A. flavus* spores derived from pure cultures using DNeasy UltraClean® Microbial Kit (Qiagen) with the following modifications. When resuspending the fungal cells in PowerBead Solution, RNase A was added to degrade any RNA present. Spores were vortexed using the MOBIO Vortex Adapter and Disrupter Genie, incubated at 65°C, and vortexed again to ensure that spores were fully ruptured. Finally, DNA was eluted in PCR-quality water for genotyping by sequencing.

### Genotyping by Sequencing

We used double digest Restriction Site-Associated DNA Sequencing (ddRADseq) to identify genome-wide Single Nucleotide Polymorphisms (SNPs) (Peterson *et al*. 2012). These dense SNP markers allow us to unambiguously track the applied *A. flavus* NRRL 21882 and NRRL 18543 strains that are the active ingredients in the Afla-Guard and AF36 biocontrol products, respectively. To determine optimal restriction enzyme combinations for ddRADseq we performed *in silico* digestions of *A. flavus* NRRL 3357 (Nierman *et al*. 2015) and *A. oryzae* RIB40 (Machida *et al*. 2005) reference genomes to identify pairs of enzymes that yielded approximately 5,000 fragments that were 350-450 bp in size. By targeting this number of fragments, we were able to estimate the maximum number of strains that could be multiplexed on an Illumina NextSeq machine (400M paired-end reads) and achieve at least 20X read coverage per individual.

Total genomic DNA was isolated and quantified using Quant-iT™ PicoGreen® dsDNA Assay Kit (Invitrogen). Two restriction enzymes, MluCI(=Tsp509I) and MspI, were identified that would allow us to multiplex up to 1,152 strains and sequence 5,000 fragments/individual to a depth of at least 20X. DNA (200 ng) was digested with both enzymes; a universal adapter was ligated to the overhangs produced by MluCI and one of 48 barcoded adapters was ligated to overhangs produced by MspI. A second quality check with Quant-iT™ PicoGreen® dsDNA Assay Kit was performed to ensure DNA was digested. Size selection was done using a Pippin Prep (Sage Science) to keep fragments in the range of 450-550 bp. This is the total size that includes the targeted 350-450 bp fragment pool plus the ligated universal and barcode adapters. A universal primer was annealed to the barcoded adapter and one of 24 indexed primers was annealed to the universal adapter followed by PCR amplification using KAPA HiFi Hotstart Readymix (Kapa BioSystems). This design ensured that only DNA sequences with both ligated adapters will be sequenced. All strains in each sublibrary were quantified using Bioanalyzer (Agilent) and then pooled in equimolar ratios. Sublibraries were quantified and a final pooling was performed before paired-end sequencing. The unique combination of indexed primer and barcoded adapter allowed for demultiplexing first by Illumina index and then by barcode. The DNA for all isolates were pooled and sequenced using 150 bp paired end reads on the Illumina NextSeq® platform (NC State University Genomic Sciences Laboratory).

### Data Processing for Variant Discovery

All ddRADseq data were analyzed using workflows implemented in a locally deployed instance of Galaxy (Jalili *et al*. 2020) at NC State University and the Mobyle SNAP Workbench (Monacell & Carbone 2014). Briefly, the process_radtags script from the Stacks package (http://catchenlab.life.illinois.edu/stacks/) was used to demultiplex 48 barcodes for each Illumina NextSeq sublibrary. Trimmomatic (Bolger *et al*. 2014) was used to trim low quality bases from the end of reads and crop bases from the ends regardless of quality. Filtered read pairs were aligned to the *A. oryzae* RIB40 reference genome using the MEM algorithm in BWA (Li & Durbin 2009). Sequence alignment (SAM) files generated from BWA-MEM were converted to Binary (BAM) files and sorted by coordinate order using SortSam in Picard tools (https://broadinstitute.github.io/picard/). PCR duplicates were removed using Picard MarkDuplicates and read groups were added for each strain using Picard AddOrReplaceReadGroups. The strains were then genotyped using the HaplotypeCaller variant discovery pipeline in GATK v3.5-2 (McKenna *et al*. 2010). GATK variant calling is designed to maximize sensitivity, so there could be many false positives. Subsequent filtering of variants eliminated false positives and negatives and was performed according to GATK Best Practices recommendations (DePristo *et al*. 2011; Van der Auwera *et al*. 2013). Variant Call Format (VCF) files from GATK were subjected to various levels of filtering using VCFtools (DePristo *et al*. 2011) and variant calls were visualized in JBrowse (Buels *et al*. 2016).

### Genome-wide Haplotype Analysis

We inferred genome-wide haplotypes (GWHs) for 815 *A. flavus* isolates using MeShClust (James *et al*. 2018) implemented in the DeCIFR toolkit (https://tools.decifr.hpc.ncsu.edu/meshclust). MeShClust uses a mean shift algorithm to cluster DNA sequence data, which performs better than other clustering methods when the exact sequence similarity threshold is not known (James *et al*. 2018). The optimal sequence similarity threshold in MeShClust was determined as the number that correctly clustered a reference panel of *A. flavus* VCGs. This was also aided with an additional parameter, the delta number, that determines how many clusters are looked around in the final clustering stage. It is expected that isolates within a VCG are very similar to each other and should cluster more tightly than with isolates from different VCGs. Using the VCG panel to guide our clustering, we also explored a more conservative approach of collapsing strains into haplotypes by excluding all sites with unknown base calls using SNAP Map (Aylor *et al*. 2006). For the larger data set of 815 isolates, it is expected that this approach would eliminate many SNPs and group isolates from different VCGs together. The distributions of GWHs were analyzed in JMP Pro 14 (SAS Institute Inc., Cary, NC, 1989-2020) using the “Summary” and “Distribution” functions. To observe trends in their distribution, the GWHs were grouped by state (TX, NC, AR, IN), substrate (kernel, soil), treatment (untreated, Afla-Guard, AF36) and sampling period (pre-application, post 3-months, post 1-year, post 3-years). Comparisons of GWHs unique and/or in common among the conditions were analyzed using Venny 2.1 (https://bioinfogp.cnb.csic.es/tools/venny/).

### Phylogenetic Inference and Patristic Distances

We inferred maximum likelihood (ML) phylogenies of *A. flavus* at different sampling time points: pre-application kernel and soil isolates, kernel isolates post 3-months, and soil and kernel isolates post 1-year. Additionally, kernel isolates in Texas were examined three-years post-application with Afla-Guard (post 3-year). ML analysis was performed using the program Randomized Axelerated Maximum Likelihood or RAxML version 8 (Stamatakis 2006) via the Cyberinfrastructure for Phylogenetic Research (CIPRES) Representational State Transfer Application Program Interface (REST API) (Miller *et al*. 2015) implemented in the DeCIFR toolkit (https://tools.decifr.hpc.ncsu.edu/denovo). The best-scoring ML majority rule consensus tree was based on 1,000 rapid bootstrap searches in RAxML using a GTRGAMMA model of rate heterogeneity with empirical base frequencies; all phylogenies were rooted with NRRL 29506 (IC277) belonging to the IB lineage, which was determined to be ancestral in previous work (Moore *et al*. 2009). Trees were visualized using the upload tree option of the Tree-Based Alignment Selector (T-BAS, version 2.3) toolkit (https://tbas.hpc.ncsu.edu/) (Carbone *et al*. 2017; Carbone *et al*. 2019).

The possibility of released biocontrol agents recombining with native strains was first examined using pairwise evolutionary distance or patristic distance. A patristic distance is the sum of the length of branches that connect any two nodes in a phylogeny. A matrix of patristic distances, normalized to a maximum value of one, was generated for all pairs of terminal nodes that represent sampled individuals. When calculated separately for phylogenies inferred from different chromosomes and compared, such distances can identify incongruences in tree topologies that may indicate genetic exchange and recombination, horizontal gene transfer and genomic reassortment (Hernandez-Lopez *et al*. 2013; Peris *et al*. 2014). Patristic distances were calculated using DendroPy (Sukumaran & Holder 2010) implemented in the DeCIFR toolkit (https://tools.decifr.hpc.ncsu.edu/patrdist). This tool displays patristic distances as a heat map using outer rings (one per chromosome) that surround the total evidence inner tree that was based on the concatenated chromosomal character matrix. In the heat map a blue color across most chromosomes indicates close genetic similarity to the selected biocontrol reference strain; a red color across one or more chromosomes indicates high genetic divergence. Inter-lineage recombination would result in strains that have blue, red and intermediate colors indicating a mixed genetic background.

### Population Structure

Population structure was first examined using principal component analysis (PCA). PCA uses a variance–covariance matrix to reduce the dimensionality of the original variables into a smaller number of new variables called principal components (Abdi & Williams 2010). The plotted principal components (eigenvectors) provide different axes of variation that explain most of the variance in the original data. The first eigenvector or PC1 considers the most variation possible with subsequent eigenvectors, PC2, PC3, etc., having less variation. Principal components were normalized to sum to 1 to reveal which eigenvectors explained more than half of the genetic variation, and the number of significant axes of variation was determined using the Tracy–Widom statistic (Tracy & Widom 1994). The number of distinct *k* clusters was determined using the Gap Statistic (Tibshirani *et al*. 2001), which is an unbiased estimate of the number of distinct clusters based on the top three PCs with the largest eigenvalues. Significant PC’s and *k* clusters were displayed in three-dimensional graphs using the scatterplot3d package in R (Ligges & Machler 2003). Tests of association of *k* cluster with lineage (IB and IC), sampling location (TX, IN, NC, AR), treatment (Afla-Guard, AF36, untreated) and substrate (soil and kernel) were performed using Fisher’s exact test, implemented in R (R Core Team 2020). Lineage-specific population parameter estimates of mean mutation rate (*θ*) and pairwise nucleotide diversity (*π*) in each state were calculated by averaging across all chromosomes in untreated and treated samples using the program SITES version 1.1 (Hey & Wakeley 1997). The pairwise fixation index (*F*_ST_) implemented in SITES was used to evaluate genetic differentiation among treatments and lineages.

The degree of genetic admixture and the optimal number of *k* clusters was determined using ParallelStructure (Besnier & Glover 2013), an R-based implementation of STRUCTURE version 2.3.4 (Falush *et al*. 2003; Pritchard *et al*. 2000) accessible via the CIPRES REST API (Miller *et al*. 2015). Structure-formatted files were generated from genome-wide SNPs using SNAP Map (Aylor *et al*. 2006). The admixture model implemented in STRUCTURE was used to assign individuals to *k* clusters. Estimates of allele frequencies and membership probabilities of individuals in subpopulations were based on a Markov Chain Monte Carlo (MCMC) strategy of 100,000 sampling iterations after a burn-in period of 50,000 iterations; three independent simulations for possible *k* values ranging from 1 to 10 were performed for each subpopulation. To determine the optimal *k*, probability distributions were examined using *LnP(D)* and delta *K* methods (Evanno *et al*. 2005) implemented in Structure Harvester v0.6.93 (Earl & vonHoldt 2011). The estimated cluster membership coefficient matrices were examined in CLUMPP v1.1.2 (Jakobsson & Rosenberg 2007) to determine the optimal number of *k* clusters across multiple runs. The individual cluster membership results from Structure were visualized using histograms in outer rings surrounding the total combined chromosomal ML phylogeny, as implemented in the DeCIFR toolkit (https://tools.decifr.hpc.ncsu.edu/structure).

### Population Parameter Estimation

Estimates of effective population sizes (*N_e_*), population migration rates (*N_e_m*; haploid migration rate) and population splitting times in years were based on the Isolation-with-Migration (IM) model assuming constant population sizes and migration rates, as implemented in the IMa3 and IMfig programs (Hey *et al*. 2018). Prior to running IM simulations, variation across each of eight *A. flavus* chromosomes was subjected to four-gamete filtering to retain the largest nonrecombining partitions. While focusing on an IM model and recombination free datasets lowers the statistical power for detecting migration rates that are greater than zero, the alternative approach of ignoring recombination and assuming a finite sites model results in disjunct posterior probability distributions and should be avoided (Hey & Wang 2019). The IMgc program (Woerner *et al*. 2007) was used to obtain recombination free chromosomal partitions using a weight ratio of 10 to 1 for individual strains and sites, respectively. This ensured that the largest number of strains and variable sites across each chromosome were retained for population parameter estimation.

Estimates of demographic parameters were based on an IM model of two populations (lineages IB and IC) and a potential third unsampled ghost population that served as an outgroup population. A mutation rate of 4.2 × 10^−11^ per base per generation in *A. flavus* (Alvarez-Escribano et al. 2019) and a generation time of 0.17 years (Horn *et al*. 2014) were used to obtain demographically scaled estimates of population parameters. Prior values for effective population sizes, migration rates, and divergence times were based on the geometric mean of Watterson’s estimate of *θ* across all eight chromosomes, calculated separately for lineages IB and IC in each state and sampling period. The upper bound for the prior on the population size parameter was set to five times the largest value of these geometric means across lineages IB and IC; the upper bound for the prior migration rates was set to five times the inverse of the geometric mean of *θ;* and the upper bound for the prior on splitting times was set to two times the geometric mean of *θ*, according to the IMa3 documentation (Hey 2019). These are considered ballpark estimates and were used as a guide for selecting a set of priors that would work across all populations.

Several preliminary short runs of a few chains were performed to select the best heating schemes that maintained high swap rates (between 0.7 and 1) between adjacent pairs of chains. Good mixing in longer runs was based on high swapping rates between successive chains, non-zero values of autocorrelations between parameter estimates across sampling iterations, and high effective sample sizes (ESS > 10,000). To ensure sufficient mixing, IMa3 runs were done with 256 heated chains with a geometric heating model (0.99 and 0.65 for the first and second heating parameter, respectively), a burn-in of 1,000,000 steps prior to sampling, and 50,000 sampled genealogies per chromosome. The final MCMC mode runs were repeated at least twice with a different random number seed to ensure convergence of parameter distributions. Migration arrows drawn using the IMfig program denoted population migration rates that were statistically significant (Nielsen & Wakeley 2001). All runs were performed through the REST API service at CIPRES (Miller *et al*. 2015) using program calls from the IMgc (https://tools.decifr.hpc.ncsu.edu/imgc) and IMa3 (https://tools.decifr.hpc.ncsu.edu/ima3) tools implemented in the DeCIFR toolkit.

### Mating Type Distribution

Mating types were scored using diagnostic ddRADseq fragments located within the *MAT1-1* idiomorph spanning positions 1,581,022 – 1,581,132 on chromosome 6 of the *A. oryzae* reference genome. A 1:1 distribution of *MAT1-1* and *MAT1-2* is indicative of populations undergoing sexual reproduction and this was tested using a binomial test implemented in MS Excel. The test was also performed on clone-corrected haplotypes to correct for skewness in mating type distributions due to clonal amplification. Clone-correction was performed by counting the total number of unique GWHs in each *MAT1-1* and *MAT1-2* category; individuals or GWHs containing both mating types were counted twice as a *MAT1-1* and a *MAT1-2*. A representative sample of 47 isolates from each *MAT* idiomorph were selected for PCR validation using *MAT1-1* and *MAT1-2* specific primers, as reported previously (Ramirez-Prado *et al*. 2008). In the presence of strong population structure, mating type distributions were examined separately for each distinct genetic cluster.

### Phylogenetic Incongruence and Recombination

Since populations of *A. flavus* are reported to have both a clonal and recombining population structure (Moore *et al*. 2013), topological concordance across chromosome and mitochondrial phylogenies was used to further examine the contributions of clonality and recombination in the evolution of lineages, GWHs, and individuals. This phylogenetic method implemented in the DeCIFR toolkit (https://tools.decifr.hpc.ncsu.edu/trees2hypha) uses the Hypha package module of Mesquite v3.51 (Maddison & Maddison 2011; Oliver *et al*. 2013) to display the clonal and recombinant history of each ancestral node and all of its descendant strains. Specifically, Hypha was used to compare the internodal support values harvested from each chromosomal phylogeny on the total evidence tree inferred from a concatenated genome-wide SNP matrix. Nodal grid support values were based on a bootstrap threshold support value of 70% and were output as node annotations on the total evidence display tree. Support values were visualized for each chromosome phylogeny using grids on branches of the display tree with colors showing node bipartitions that were supported at a bootstrap support value ≥70% (black node) or <70% (white node); if the specific node bipartition was not found in the display tree this was reported as missing or inapplicable (grey node). Grids that were filled in with mostly black squares indicated that the descendants of that node were predominantly clonal. High conflict (red node) was used to indicate a node bipartition in the chromosome tree that conflicted with the displayed tree at a bootstrap support value ≥70% most likely due to recent recombination (i.e., independent assortment and crossovers) among strains in descendant branches at terminal nodes or ancestral recombination at internal nodes.

Low conflict (cyan node) was used for nodes that were not recovered by the bootstrap analysis because there was either insufficient variation or too much confounding variation (i.e., homoplasy) due to recombination among strains in descendant branches. We would expect node bipartitions that comprise strains that belong to the same GWH to be congruent across all chromosomes. If GWHs associate closely with VCGs they can be used as a proxy for VCGs. In the case of reference strains that are known *a priori* to belong to the same VCG and are of the same mating type, high conflict at one or more chromosomes provides evidence of recombination in the immediate common ancestor.

### Chromosomal Linkage Disequilibrium

Chromosome-wide linkage disequilibrium (LD) between SNP markers distributed across each chromosome was performed using Haploview 4.2 (Barrett *et al*. 2005). Both *r^2^* and *D’* pairwise LD measures were calculated between adjacent SNP markers in all populations for each sampling period: pre-application, 3-months, 1-year and 3-years post-treatment. Intra-chromosomal LD blocks were estimated using the Solid Spine (SS) method of Haploview using the default parameters and a missingCutoff of 0.8. The SS algorithm identifies an LD block if the first and last markers in a block are in strong LD with all intermediate markers, but those intermediate markers can be in weak LD or no LD with each other. Haploview outlines the edge of the spine of strong LD in a triangular matrix of pairwise LD statistics. The coloring scheme is based on the values of *D’* and LOD (logarithm of the likelihood-odds ratio) where bright red represents strong LD (LOD ≥ 2, *D’* = 1), shades of pink/red represent intermediate LD (LOD ≥ 2, *D’* < 1), blue represents weak LD (LOD < 2, *D’* = 1) and white represents no LD (LOD < 2, *D’* < 1). The strength of LD was determined by averaging *r^2^* for each lineage, chromosome, and sampling period. We expect the release of biocontrol strains to initially increase the clonal component of population in the short-term and result in higher mean *r^2^* values when compared to mean *r^2^* values in pre-application fields. In the more long-term, the released biocontrol strains are expected to increase the opportunities for both intra- and inter-lineage recombination and result in lower mean *r^2^* values than in pre-application fields.

### Aflatoxin Quantitation and Analysis

For each state (TX, NC, AR, and IN) and sampling period (pre-application, post 3- months, post 1-year, and post 3-years) at least one strain from each lineage (IB and IC), treatment (untreated, Afla-Guard, and AF36), substrate (soil and kernel), and aflatoxin cluster configurations (missing, partial, and full) was selected when available for aflatoxin B_1_ quantitation. Aflatoxin was quantified using a method which allows detection of AF B_1_ production from fungal mycelia in liquid culture (Gell & Carbone 2019). Reference strains were included for comparison with previous quantification methods (Horn & Dorner 1999; Horn & Greene 1995; Horn *et al*. 1996). Briefly, isolates were grown on Potato Dextrose Agar (PDA) plates at 30°C for a total of 7 days: 5 days in the dark and 2 days in light. Glass vials with 7 mL of YES media (Koehler *et al*. 1975) were inoculated with a loop-full of conidia for each isolate, with three replicates grown per isolate. Liquid cultures were incubated in the light at 30°C for 7 days. For each culture, a 1-mL aliquot of media was transferred to a new vial, 1 mL of chloroform was added, and samples were vortexed and then allowed to separate at rest. A total of 500 µL of the chloroform layer was transferred to a clean vial and evaporated under nitrogen stream. Dried aflatoxin samples were resuspended in 1 mL of methanol for analysis and purified by passing through 1-mL polypropylene SPE tubes containing a 200 µL tube full of alumina basic.

Aflatoxins were quantified by HPLC on two different instruments: (1) HPLC with fluorescence detection at the Biomanufacturing Training and Education Center’s Bioprocess and Analytical Services at North Carolina State University, and (2) HPLC with mass spectrometry detection at our laboratory at North Carolina State University. The detection by HPLC-fluorescence (excitation 365 nm, emission 455 nm) followed a previously published protocol (Huang & Elmashni 2007). Serial dilutions were used to determine the limit of aflatoxin B_1_ detection at 0.0020 µg/mL and quantification at 0.0039 µg/mL (Gell & Carbone 2019).

HPLC with mass spectrometry detection was done on a quadrupole LC-MS (Ultimate 3000 UPLC/ISQ EC, Thermo Fisher Scientific) in Single Ion Monitoring (SIM) mode. The instrument was also equipped with UV-vis Diode Array and Charged Aerosol Detectors. The chromatographic conditions were based on a recent mass spectrometry-based method for analysis of mycotoxins in cannabis matrices (Maguire 2019).

Separations were performed on an ACE Excel 1.7 µm C18-PFP column (100 x 3.0 mm) in a thermostated column compartment at 45 °C. The mobile phase consisted of a gradient of solutions A and B (Solution A: water + 1 mM ammonium formate + 0.1% formic acid; Solution B: methanol + 1 mM ammonium formate + 0.1% formic acid).

During the 9-minute gradient the proportion of A and B solutions was programmed as follows: 0-1 min constant at 95% A / 5% B; 1-2 min linear gradient to 55% A / 45% B; 2-8 min linear gradient to 10% A / 90% B; 8-9 min: constant at 10% A / 90% B. The column was re-equilibrated for 2.5 minutes at 95% A / 5% B before the next injection (injection volume = 50 µL) and the flow rate was constant at 0.8 mL/min. The mass spectrometer detector was set to positive mode Single Ion Monitoring (SIM) targeting Aflatoxin B_1_ (*m/z* = 313.1), B_2_ (*m/z* = 315.1), G_1_ (*m/z* = 329.1), and G_2_ (*m/z* = 331.1) and positive mode scans were acquired during the same run. The peaks for aflatoxin B_1_ (retention time = 5.87 min) and B_2_ (retention time = 5.61 min) were baseline-separated with a low background intensity. The detection limit determined by repeated injection of standards and samples was ∼0.3 ng/mL (∼1 nmol/L) in the injected sample or ∼1 ng/mL (∼3 nmol/L) in the original undiluted sample extracts. Quantification with aflatoxin B_1_ and B_2_ standards (Sigma-Aldrich A6636-5MG) was based on a six-point calibration curve in the range of 0.001-0.5 µg/mL. Standards were analyzed at the beginning and end of every analysis batch, and two blank injections were included between samples of different cultures. The extracted samples in methanol were diluted with A and B solutions to 70% aqueous / 30% organic to achieve symmetric chromatographic peak shapes. The dilution was adjusted so that injected samples contained < 0.5 µg/mL (< 1.6 µmol/L) of aflatoxin B_1_ to remain within the linear range and minimize carry over. The cultures produced primarily aflatoxin B_1_ and concentrations before dilution were estimated using a fluorescence plate reader (BioTek Synergy HTX, excitation: 360 nm, emission 460 nm).

To determine if there was quantitative variation in B_1_ aflatoxin concentrations between lineages IB and IC, aflatoxin frequency distribution plots and cumulative distribution functions were graphically portrayed for each lineage. Mean toxin concentrations were determined and significant differences in toxin distributions between lineages IB and IC were tested by performing Kolmogorov-Smirnov tests, as implemented in Matlab (MathWorks Inc., Natick, MA, USA).

## RESULTS

### Sampling and Variant Discovery

A total sample of 815 isolates was examined across all states, treatments, and sampling periods (Tables 1, S1). Also included was a panel of 26 reference strains for which mating type (Ramirez-Prado *et al*. 2008), toxin profile (Horn *et al*. 1996), VCGs (Horn & Greene 1995) and multilocus haplotypes (Moore *et al*. 2009) were known and used to cross-validate genotyping by sequencing results (Table S2). A total of 672,302 unfiltered variants were discovered and 6,806 biallelic variants were retained after filtering (Table 2). The filtering was performed using a minimum and maximum value of the minor allele frequency set to 0.001 and 0.5, respectively, to ensure that rare sequence variants were kept for genotyping individual biocontrol strains with high resolution. Moreover, because phylogenetic inference, calculation of patristic distances, phylogenetic incongruence, and LD analysis rely on informative SNPs that group together two or more individuals, filtering included a genotype call rate of 90% across all individuals.

**Table 1.**
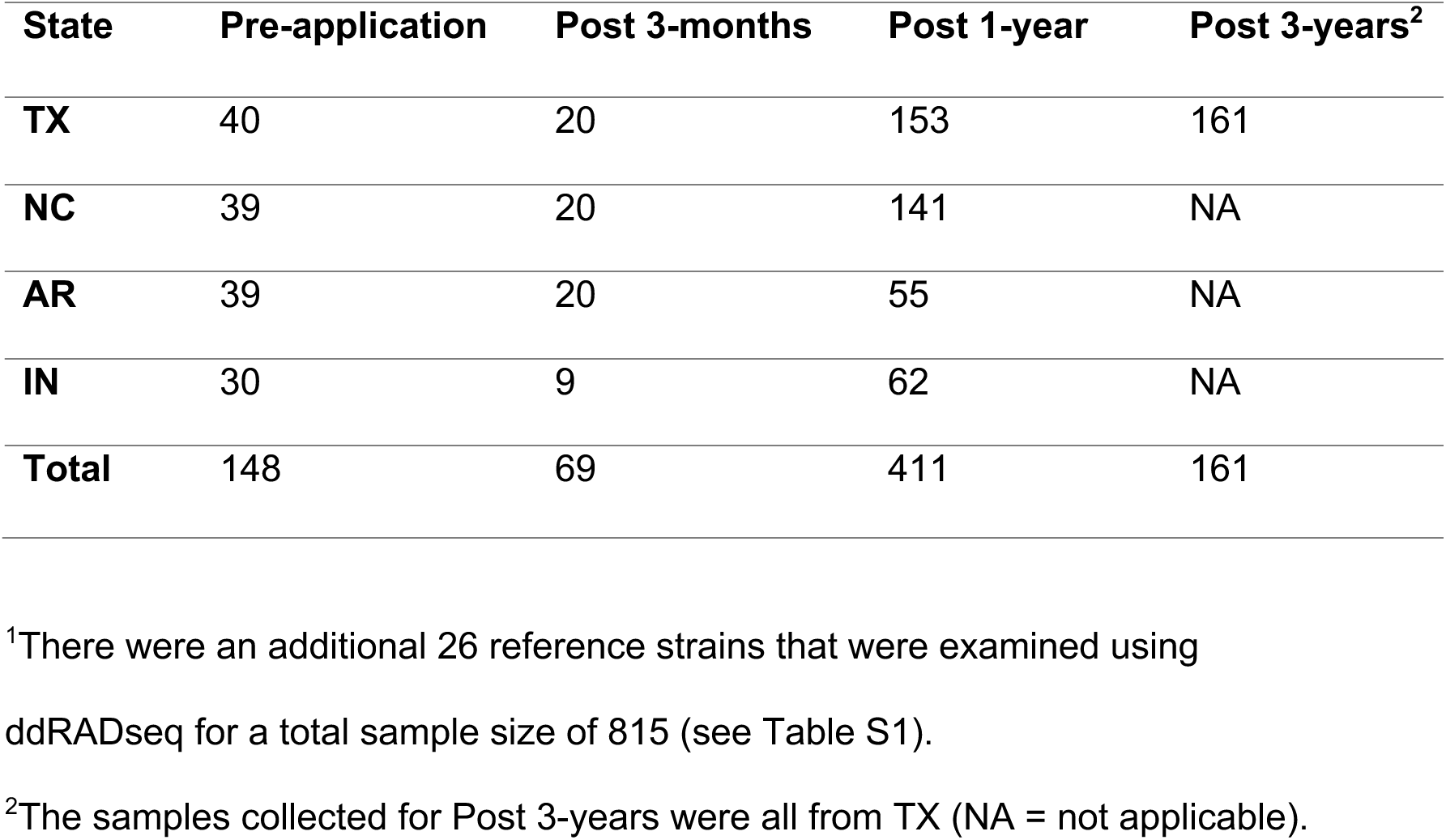
Population Samples of *A. flavus^1^* for different sampling periods.

**Table 2.** Number of variable sites for phylogenetic and population genomic analysis

| **Sampling period** | **Sample size<sup>1</sup>** | **Unfiltered** | **Filtered<sup>2</sup>** | **SNPs<sup>3</sup>** |
| --- | --- | --- | --- | --- |
| **Pre-application** | 172 | 451,283 | 27,529 | 610 |
| **Post 3-months** | 93 | 451,283 | 27,529 | 592 |
| **Post 1-year** | 436 | 447,324 | 21,260 | 713 |
| **Post 3-years** | 187 | 271,813 | 14,677 | 1,282 |
| **TOTAL** | 815 | 672,302 | 6,806 | 5,750<sup>4</sup> |

<sup>1</sup>Number including reference strains.
<sup>2</sup>Number of sites in character matrix for phylogenomic inference and patristic distances.
<sup>3</sup>Number of SNPs (excluding ambiguous sites) for PCA and STRUCTURE analysis.
<sup>4</sup>Number of SNPs (ambiguous sites included) in the total sample for clone-correction and population parameter estimation.

### Genome-wide Haplotype Inference

Overall, a total of 353 unique genome-wide haplotypes (GWH H0-H352) were inferred among the 815 individual strains (Table S1). This was based on clustering 6,806 variants using a sequence similarity threshold of 0.987 and a delta value of 5. The two biocontrol strains IC201(=Afla-Guard; VCG 24) and IC1179 (=AF36; VCG YV36) belonged to haplotypes H1 and H115, respectively. Also included in H1 was IC253 from the same VCG 24 as IC201 (Table S2). Other VCGs with multiple representatives were also clustered into unique haplotypes such as IC221 and IC222 (H40; VCG 4). The 0.987 clustering threshold separated reference strains that belong to different VCGs into distinct haplotypes: IC277 (H29; VCG 32), IC278 (H338; VCG 33), IC301 (H333; VCG 56), IC307 (H341; VCG 62), IC308 (H336; VCG 63), and IC313 (H271; VCG 76). Two VCGs with multiple strains: VCG 1 (IC217, IC218) and VCG 5 (IC225, IC226) were clustered into the same haplotype H21. During the clustering processes MeShClust generates a semi-ordered list of clusters such that adjacent clusters (i.e., GWH numbers) have more similar sequences. This was observed with H69 (IC220; VCG 2) and H70 (IC219; VCG 2), and with H329 (IC230; VCG 6) and H330 (IC229; VCG 6).

In subsequent haplotype analyses, strains from GA that were used as VCG reference standards in the ddRADseq analysis pipeline were excluded. The haplotype analyses were conducted on the remaining 628 strains from AR, IN, NC and TX for the pre-application, 3-month and 1-year post-treatments. The 3-year post-treatment analysis in TX included 161 strains and was conducted separately. The 628 strains represented 276 unique genome-wide haplotypes (Table S3). The haplotype frequency distribution is shown in Table S3, where haplotypes H1 (28%) and H115 (3%) highlighted in red bars denote the IC201 (Afla-Guard) and IC1179 (AF36) biocontrol haplotypes, respectively. The remaining 255 genome-wide haplotypes each had a single strain and a frequency of 0.16%, which accounted for 41% of the total sample.

When summarized by state (AR, IN, NC, TX), the only two haplotypes that were shared among all four states were the two biocontrol haplotypes H1 and H115 (Table S4). The majority of GWHs were unique to individual states: TX (*n* = 70); NC (*n* = 95); AR (*n* = 65); and IN (*n* = 36)). The remaining eight haplotypes were shared between two or three different states. Among the 276 GWHs, eleven were shared between soil and kernel samples (including H1 and H115), 114 haplotypes were unique to the kernel samples, and 151 were unique to the soil samples (Table S5). Further summarizing the soil and kernel isolates by state revealed additional shared haplotypes on a regional scale. There were five shared haplotypes in NC between kernels and soil (H1, H115, H140, H229, H240), six shared haplotypes in TX (H1, H21, H27, H28, H29, H55); and only the biocontrol haplotype H1 was shared between kernel and soil in AR and IN (Table S6). Overall, only five haplotypes (H1, H140, H22, H229, H240) were shared across the three different sampling periods: pre-application, post 3-months, and post 1- year, and twelve haplotypes were in common between the pre-application and post 1-year (Table S7). There was a 2.5-fold increase in the number of unique haplotypes across all states in post 1-year (*n* = 178) compared to the pre-application (*n* = 73); a similar fold increase was observed in NC and TX and to a lesser extent in AR and IN (Table S7).

### Phylogenetic Inference and Patristic Distances

Figure 1 shows the best-scoring ML phylogenies inferred for longitudinally sampled cornfields in TX, NC, AR and IN for the pre-application, post 3-months and post 1-year sampling periods. The best ML phylogeny for the commercial TX cornfields is shown in Figure 2. Several consistent patterns were observed across all phylogenies. First, two distinct evolutionary lineages IB and IC based on sequence variation across eight *A. flavus* chromosomes were found in all sampled cornfields and treatments. Lineages were identified using reference isolates that were included as standards in ddRADseq (Table S2) and from pairwise patristic distances separating isolates from either IC201 (Afla-Guard – AG; lineage IB) or IC1179 (AF36; lineage IC). Second, there was evidence of clonality (i.e., isolates with the same patristic distance from either AG or AF36 across all chromosomes) and recombination (i.e., isolates with different patristic distances across chromosomes) in both lineages IB (Figs. 1A, 2A) and IC (Figs. 1B, 2B). The strong divergence between IB and IC lineages made it difficult to visualize small differences in patristic distances within lineage IB (Figs. 1A, 2A). Better resolution was achieved by adjusting the heat map of patristic distances so that the maximum color value fell within lineage IB (Figs. 1C, 2C). This adjustment revealed heterogeneity in patristic distances across chromosomes in one sublineage within lineage IB and a second more homogeneous sublineage comprised almost entirely of isolates with a missing aflatoxin gene cluster.

**Figure 1.**
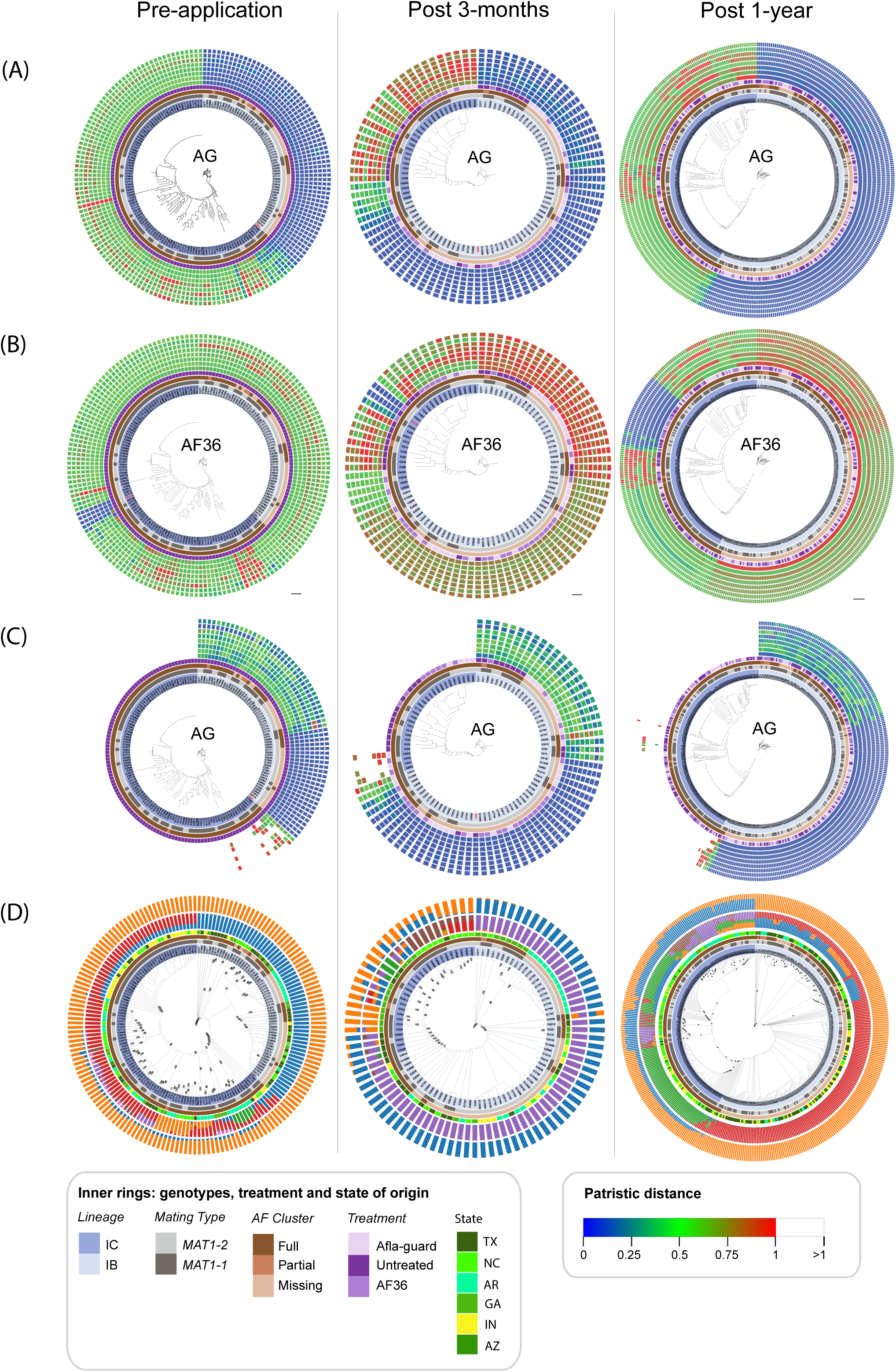
Longitudinal population dynamics of *A. flavus* across four states. *Aspergillus flavus* biocontrol strains Afla-Guard=AG and AF36 were applied to cornfields in TX, NC, AR, and IN. In parts (A), (B) and (C) the four innermost rings show the overall recovery and frequency of *A. flavus* evolutionary lineage (IB, IC), mating types (*MAT1-1, MAT1-2*), aflatoxin (AF) cluster configuration (full, partial, missing) and treatment (untreated, Afla-Guard, AF36) across field populations. In (D) the treatment ring was replaced with sampling locations for field experiments (TX, NC, AR, IN) and reference strains (GA, AZ). The phylogenies displayed in the center of the rings are based on maximum likelihood searches using variation across the entire genome, with branches scaled to the number of nucleotide substitutions per site; the scale bar (0.02 substitutions per site) is shown at bottom of tree in part (B). Patristic distance is shown in the eight outermost rings and used to examine variation among all strains as compared to either Afla-Guard=AG (A) or AF36 (B). The distance of each strain from AG or AF36 as a reference is shown using a heat map, where a value of zero (blue) indicates very close genetic similarity of the strain to the reference and a value of one (red) is high genetic dissimilarity. In part (C) the patristic scale is adjusted so that the maximum color value falls within lineage IB, revealing heterogeneity in patristic distances and potential clonality and recombination in lineage IB, not observed in (A). Part (D) shows results from STRUCTURE analysis in the two outermost rings; the inner rings are the best clusters inferred from STRUCTURE LnP(D) and the outer rings are distinct clusters inferred using Evanno’s method.

**Figure 2.**
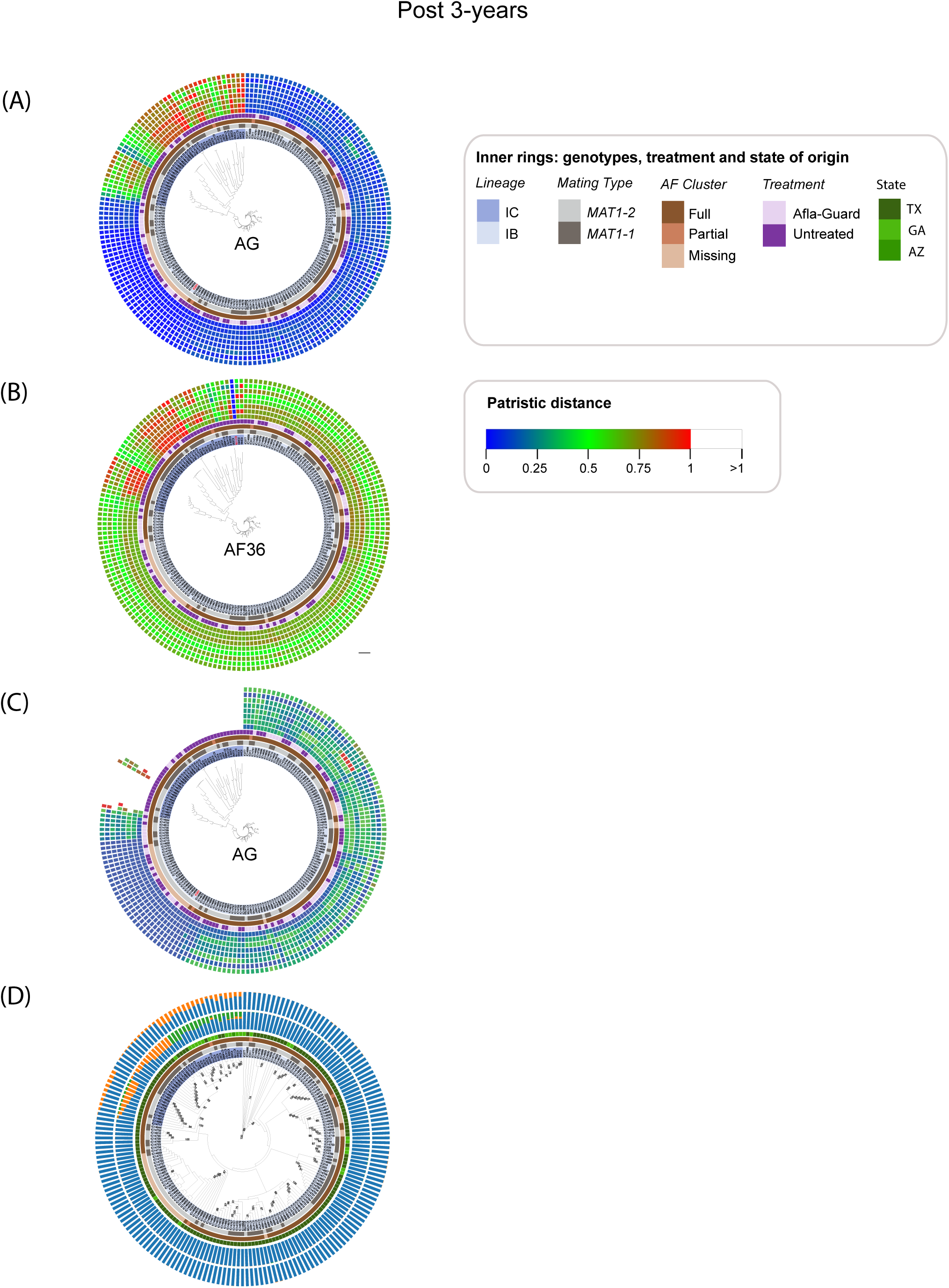
Population structure of *A. flavus* in commercial TX cornfields. The phylogenies displayed in the center of the rings are based on maximum likelihood searches using variation across the entire genome, with branches scaled to the number of nucleotide substitutions per site; the scale bar (0.04 substitutions per site) is shown at bottom of tree in part (B). In parts (A), (B) and (C) the four innermost rings show the overall recovery and frequency of *A. flavus* evolutionary lineage (IB, IC), mating types (*MAT1-1, MAT1-2*), aflatoxin (AF) cluster configuration (full, partial, missing) and treatment (untreated, Afla-Guard) across field populations. The eight outermost patristic rings show the extent of clonality and recombination with respect to Afla-Guard (A) or AF36 (B); AF36 was included in the patristic analysis of the TX cornfields, that were treated only with AG, to determine if AF36 occurred naturally in untreated plots. In part (C) patristic distances were adjusted to filter out lineage IC strains that represent distances greater than one (white). The two outermost STRUCTURE rings (D) show the number of distinct genetic clusters and degree of lineage admixture based on LnP(D) for the inner ring and Evanno for the outer.

Phylogenetic reconstructions across different sampling periods allowed assignment of strains into lineage IB or IC. The distribution of lineages in soil and kernel samples is summarized in Table S8. Although both lineages IB and IC were represented in kernel and soil samples, the fraction of each lineage varied by substrate (kernel, soil), sampling period (pre-treatment, post 3-months, post 1-year) and state (TX, NC, AR, IN). Across all states, lineage IB predominated in kernel populations 3-months after biocontrol application; soil populations were more skewed to lineage IC before biocontrols were applied (Table S8). Texas and North Carolina consistently showed opposing lineage frequency distributions across the same substrates and sampling periods. Lineage IC was the dominant lineage in NC kernels and soil populations before biocontrols were applied and 1-year later, whereas lineage IB predominated in TX. Although TX and NC showed different dominant lineages, within each state the lineage ratio was conserved across the untreated, AF36 and Afla-Guard treated plots (Table S8). In the case of TX, the skew to lineage IB was also observed in the post 3-years untreated and treated plots. Aflatoxin cluster structure (missing, partial, or full) corresponded closely with lineage membership (Fig. 1; Table S9). Across all three sampling periods (pre-treatment, post 3-months, post 1-year) strains in lineage IC had full clusters with the only exception observed for two partial cluster strains (IC13995, IC11274) that were sampled 1-year after biocontrol application. By comparison, lineage IB strains harbored full, partial, or missing AF clusters across the three sampling periods, but their frequency varied across states with TX having a greater proportion (65%) of full cluster strains in lineage IB compared to other states (Table S9). Similarly, lineage IB predominated in TX commercial fields and 80% of the isolates in the untreated and treated TX commercial fields had full AF gene clusters (Fig. 2; Table S10).

### Population Structure

The Gap statistic in PCA analysis revealed two distinct clusters across all sampling periods that were significantly associated with lineage (*P* < 0.001) and state (*P* < 0.001) (Fig. S1). There were consistent and significant differences in the proportions of lineages IB and IC across sampling periods and between untreated and treated fields. For example, the post 1-year untreated and treated populations of *A. flavus* in TX were significantly (*P* < 0.05) skewed to lineage IB, whereas populations in NC and AR were significantly skewed (*P* < 0.05) to lineage IC (Table 4). Similar trends were observed in NC and AR cornfields prior to biocontrol application where *A. flavus* field populations were significantly (*P* < 0.01) skewed to lineage IC. The observed lineage skew to IB in the post 1-year TX cornfields was also observed in the post 3-year commercial TX fields. There was a significant association (*P* < 0.05) of lineage with treatment (Afla-Guard, AF36) in the post 3-months fields in TX, NC, AR, and IN, but no significant association (*P =* 0.13) of lineage with treatment (Afla-Guard, AF36, untreated) in the post 1-year across the four states. The widely dispersed data points in principal component space indicates greater sequence diversity within lineage IC compared to IB in pre-treatment fields (Fig. S1). This was also supported by the per base pair estimates of the population mean mutation rate and nucleotide diversity showing a 5-fold higher value in IC compared to IB in the pre-treatment fields in AR, NC and TX (Table S11). A similar trend of higher diversity in the IC lineage compared to IB was observed for treated fields post 3-months and for treated and untreated fields, sampled one year and three years after biocontrol application (Table S11).

STRUCTURE admixture analysis of the pre-treatment populations determined a best value of *k* = 2 clusters using the Evanno method and *k* = 7 clusters based on STRUCTURE LnP(D) (Table S12). The two outermost rings in Figure 1D show the ancestral composition of each isolate for the three different sampling periods using the Evanno (inner ring) and LnP(D) (outer ring) cluster inference method. In the pre-treatment populations, lineage IB has a single dominant ancestry (i.e., blue cluster) whereas isolates in IC are a mix of several populations with one predominant genetic ancestry (i.e., red cluster) (Fig. 1D). Most of the isolates in lineage IC have a mixed ancestry, with the greatest heterogeneity observed for isolates with long branches and bootstrap support values >70% in terminal clades (Figs. 1C, 1D). Clonal lineages show identical genetic backgrounds, for example, the largest clonal lineage in IC includes isolates that have the same proportion of two different ancestries (i.e., blue and red clusters). The shared genetic ancestry of the blue cluster is evidence of admixture between lineages IB and IC in the pre-treatment fields. In contrast to lineage IB which shows a single genetic background, the highly heterogenous genetic background for many strains in lineage IC indicates extensive genetic admixture and a large, structured population.

Evanno and Structure LnP(D) yielded best cluster estimates of *k* = 2 and *k* = 6, respectively, for populations sampled post 3-months (Table S13) and post 1-year (Table S14). Lineage admixture was also observed in *A. flavus* isolates sampled post 3- months. For example, based on the Evanno clustering method, IC6338 from TX sampled from the AF36 treated plot contained a 49% of the dominant genetic background in IB (i.e., blue cluster) and 51% of the genetic background that was detected in the IC ancestry (i.e., orange cluster) (Fig. 1D; Table S13). The lineage ancestry was more difficult to discern in the post 1-year fields for *k* = 6 (inner ring; Fig. 1D; Table S14). For example, IC11367 and IC15888 are from TX and NC, respectively, and contained approximately 34% of the dominant genetic background in IB (i.e., red cluster) and 65% of another ancestry (i.e., blue cluster). The outer ring representing *k* = 2 in post 1-year indicates that isolates in lineage IC have a mixed ancestry with varying contributions from lineage IB (Fig. 1D). The TX commercial cornfields that were sampled 3 years after Afla-Guard was applied were significantly skewed (*P* < 0.01) to lineage IB (Table 4) and were predominantly of a single ancestry (Fig. 2D; Table S15). There was also evidence of mixing of IB with IC and many isolates in lineage IC contained orange and blue clusters in their ancestry; the third population (green cluster) is population subdivision with the reference isolates from GA. Although genetic admixture between lineages IB and IC was detected across all treatments, there was limited structure within lineage IB and *F*_ST_ values were greater than 0.5 (Table S11) indicating significant differentiation between lineages.

### Population Parameter Estimation

We estimated population migration rate parameters, population splitting times and effective population sizes for *A. flavus* lineages IB and IC in the untreated and treated samples for the longitudinally sampled populations in TX, NC, AR, and IN, and the TX commercial fields. Given the possibility that these lineages may have undergone genetic exchange with other *A. flavus* lineages (e.g., lineage IA; Geiser *et al*. 2000) and closely related species in *Aspergillus* section *Flavi,* we also performed IMa3 analyses with a third unsampled ghost population. Patterns of migration differed between untreated and treated populations. In the TX field populations, population migration rates were found to not differ significantly from zero in the pre-application and post 1-year untreated samples, and to be significant and unidirectional in the post 3-months and post 1-year treated samples without (Fig. 3) or with (Fig. 4S) an unsampled ghost population. When bidirectional gene flow was detected, for example, in the NC pre-application and the AR post 1-year samples (Fig. S3), it was asymmetrical with more significant migration from IB into IC. Even the smaller sample sizes in IN yielded a significant signal of asymmetrical migration into IC. Across all states and sampled lineages, the ghost populations were much older than the common ancestor of IB and IC populations and there was evidence of significant asymmetrical gene flow from the ghost into the IC lineage (Figs. S2, S3).

**Figure 3.**
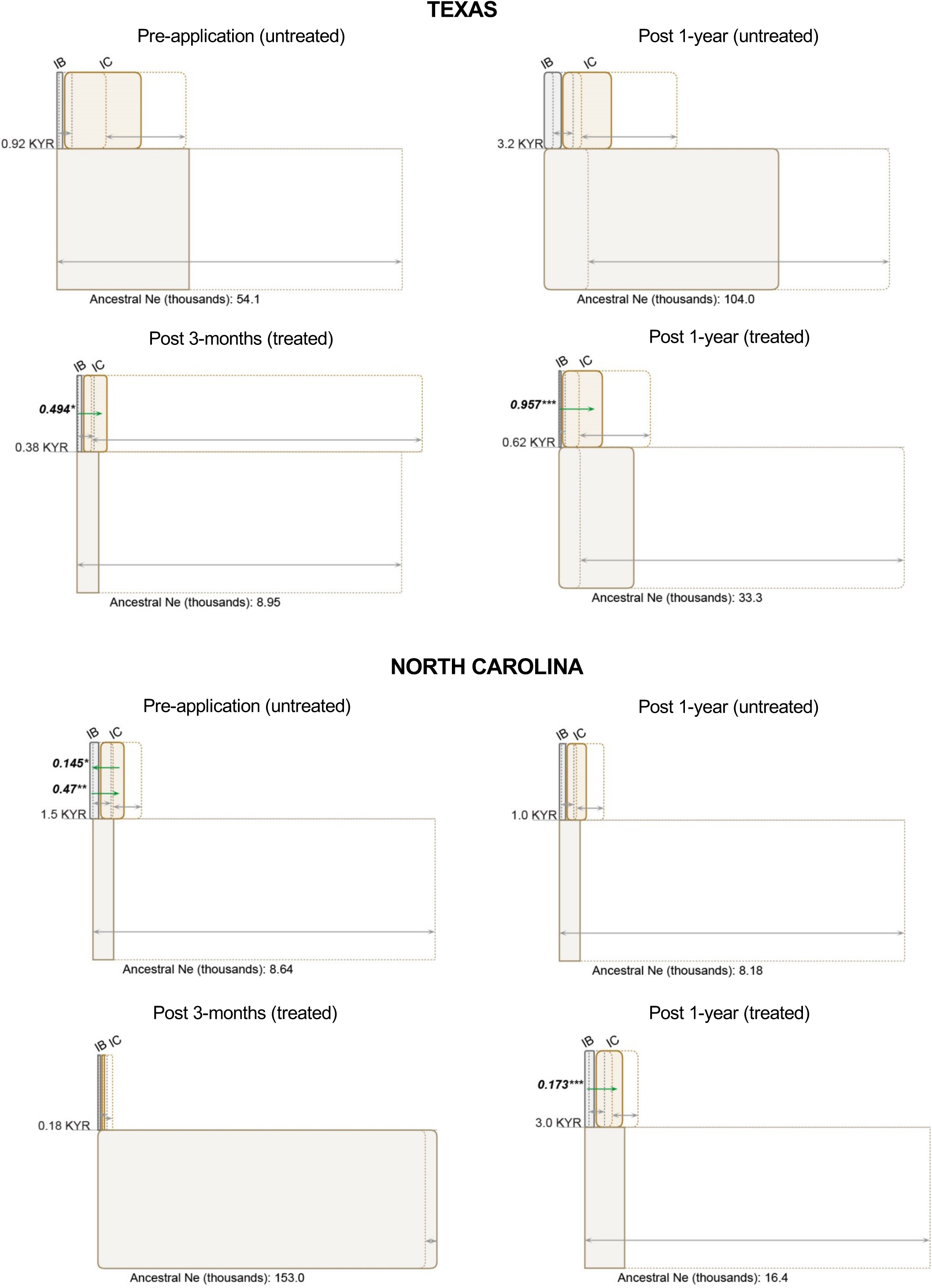
A schematic representation of isolation with migration for *A. flavus* populations in TX and NC generated by IMa3 and the IMfig program. The phylogeny for untreated and treated populations is depicted as a hierarchical series of boxes, with ancestor boxes connecting descendant populations of lineages IB and IC, and the width of boxes proportional to the estimated *N_e_*. The 95% confidence intervals for each *N_e_* value are shown as dashed lines to the right of the left side of the corresponding population box. Gray arrows to the 95% *N_e_* intervals extend on either side of the right side of each population box. Splitting times, positioned at even intervals, are depicted as solid horizontal lines, with text values on the left in units of thousand years ago (KYA). Migration arrows (in green) indicate the estimated population migration rate (*N_e_m*) values from one population into another from when the populations diverged from a common ancestor. Arrows are shown only for migration rates that are statistically significant (* *p* < 0.05, ** *p* < 0.01, *** *p* < 0.001). Estimates assumed a generation time of 0.17 years and a mutation rate of 4.2 × 10^−11^ per base per generation.

Population splitting times for lineages IB and IC in the longitudinal field experiments ranged from 0.38 to 3.2 thousand years ago (KYA) in TX and from 0.18 to 3.0 KYA in NC (Fig. 3). Splitting time and effective population size of the common ancestor of the IB and IC lineages in AR were very similar to those in NC (Fig. S3). Effective population sizes of the IC lineage in TX decreased from 31,000 in the pre-application field plots to 11,500 in the post 1-year treated plots (Fig. 3). By contrast effective population sizes in NC for the IC lineage increased from 8,640 before biocontrols were applied to 11,500 in the 1-year post application field plots. Across all states and sampling periods the ghost populations were older and had a larger effective population size relative to lineages IB and IC. The common ancestor of the ghost and the sampled lineages in TX and NC ranged from 4.0 to 56.0 KYA, which was much older than the range of splitting times (0.23 – 2.2 KYA) of the common ancestor of the sampled IB and IC lineages (Fig. S2).

Estimates of the time to the most recent common ancestor of lineages IB and IC were much larger in the post 3-years treated TX commercial plots (52.0 KYA) compared to the untreated plots (0.52 KYA) (Fig. S4). However, the estimated splitting time of the common ancestor of the ghost population in the post 3-years treated plots (57.0 KYR) was very similar to the splitting time of lineages IB and IC without the ghost (52.0 KYA). In the model without a ghost, the effective population size of lineage IC in the post 3- year untreated plots was 22,300 which was higher than the 5,300 estimated for the post 3-year treated plots (Fig. S4).

### Mating Type and Lineage Distribution

The overall distribution of mating types for the 628 strains across all four states is summarized in Table S16. Out of a total of 276 unique haplotypes, 168 and 95 were exclusively *MAT1-1* and *MAT1-2,* respectively. There were 13 haplotypes with a mix of *MAT1-1* and *MAT1-2* strains; most notable in this set were haplotypes H1 and H115 which included the strains used in the Afla-Guard and AF36 biocontrol formulations, respectively. There were 72 *MAT1-1* and 105 *MAT1-2* strains in haplotype H1; 8 *MAT1- 1* and 11 *MAT1-2* strains in haplotype H115. The other eleven haplotypes were significantly skewed to one mating type which suggests that they comprise at most one or two clonal lineages. Mating type distributions were further examined separately for the untreated and treated plots across each state and sampling period (Table 3). Except for two marginally significant values, overall mating type ratios were approximately 1:1 and not significantly different for both the uncorrected and clone-corrected samples.

**Table 3.** Distribution of mating types in *A. flavus* across states, treatments and years^1^

|  | Pre-application |  | Post 3-months |  | Post 1-year |  |  |  | Post 3-years |  |  |  |
| --- | --- | --- | --- | --- | --- | --- | --- | --- | --- | --- | --- | --- |
|  | Untreated |  | Treated<sup>2</sup> |  | Untreated |  | Treated<sup>2</sup> |  | Untreated |  | Treated<sup>3</sup> |  |
| State | *MAT1-1* | *MAT1-2* | *MAT1-1* | *MAT1-2* | *MAT1-1* | *MAT1-2* | *MAT1-1* | *MAT1-2* | *MAT1-1* | *MAT1-2* | *MAT1-1* | *MAT1-2* |
| TX | 21(12) | 19(9) | 13(7) | 7(4) | 19(11) | 30(10) | 59(29) | 45(20) | 31(26) | 53(26) | 44(20) | 33(16) |
| NC | 19(15) | 20(15) | 8(5) | 12(3) | 29(15) | 17(14) | 64(37*) | 31(20) |  |  |  |  |
| AR | 20(18) | 19(12) | 0(0) | 20(2) | 6(6) | 6(4) | 26(24) | 17(14) |  |  |  |  |
| IN | 28(12*) | 2(2) | 1(1) | 8(1) | 8(7) | 2(2) | 39(13) | 13(8) |  |  |  |  |

<sup>1</sup>Clone corrected number for each mating type is shown in parentheses; 0.01 < \*P < 0.05. Significance for the uncorrected samples is
not shown.
<sup>2</sup>Plots were treated with either Afla-Guard or AF36 biocontrol products.
<sup>3</sup>Plots were only treated with Afla-Guard biocontrol.

By contrast, the frequency distribution of haplotypes in lineages IB and IC was not proportional across different states and treatments (Table 4). For example, the lineage clone-corrected haplotype counts in both NC and AR untreated samples were significantly different (*P* < 0.01) and larger for lineage IC than for IB. The same skew to lineage IC with was observed in the post 1-year treated samples in NC and AR. A different trend was observed in the TX post 1-year samples where the skew was significant (*P* < 0.05) and in favor of lineage IB for both the untreated and treated clone-corrected haplotype counts. This was also the patterns for the TX commercial fields 3- years after being treated with Afla-Guard.

**Table 4.** Distribution of lineages in *A. flavus* across states, treatments and years^1^

|  | Pre-application |  | Post 3-months |  | Post 1-year |  |  |  | Post 3-years |  |  |  |
| --- | --- | --- | --- | --- | --- | --- | --- | --- | --- | --- | --- | --- |
|  | Untreated |  | Treated<sup>2</sup> |  | Untreated |  | Treated<sup>2</sup> |  | Untreated |  | Treated<sup>3</sup> |  |
| State | IB | IC | IB | IC | IB | IC | IB | IC | IB | IC | IB | IC |
| TX | 26(8) | 14(10) | 13(5) | 7(6) | 45(*14) | 4(4) | 93(**35) | 11(10) | 61(28) | 23(22) | 72(**24) | 5(4) |
| NC | 11(7) | 28(**23) | 11(4) | 9(3) | 16(7) | 30(*21) | 17(5) | 78(**46) |  |  |  |  |
| AR | 11(4) | 28(**26) | 20(2) | 0(0) | 6(3) | 6(6) | 17(9) | 26(**26) |  |  |  |  |
| IN | 3(3) | 27(11) | 9(2) | 0(0) | 2(2) | 8(7) | 40(11) | 12(7) |  |  |  |  |

<sup>1</sup>Clone corrected number and significance for each lineage is shown in parentheses; $0.01 < *P < 0.05$ and $**P < 0.01$ . Significance for the uncorrected samples is not shown.
<sup>2</sup>Plots were treated with either Afla-Guard or AF36 biocontrol products.
<sup>3</sup>Plots were only treated with Afla-Guard biocontrol.

### Phylogenetic Congruence and Recombination

Examination of phylogenetic congruence across chromosomal phylogenies including the mitochondrial genome showed evidence of extensive conflict in deep branches and phylogenetic concordance in terminal branches and clades (Fig. S5). As expected, node bipartitions where the descendants were isolates that were very similar genome-wide most likely belonged to the same VCG and showed ≥70 bootstrap support or low conflict across most chromosomes. For example, the Afla-Guard strain (=IC201; *MAT1-2*) which belongs to VCG 24 was widely distributed across TX, IN, NC, and AR in the pre-application field plots (Fig. S5A). A total 22 strains were very similar and monophyletic with IC201 and can be putatively assigned to VCG 24 in the pre-application fields; two additional strains (TX, IC6357; AR, IC7641) in this clade that were *MAT1-1* descended from a recombinant ancestor with low conflict across all chromosomes (Fig. S6). By contrast, the AF36 strain (=IC1179) which belongs to VCG YV36 shared a recent common ancestor with only one strain in AR (IC7716) and one in TX (IC6169) in the pre-application fields. The largest clade with very similar strains in lineage IC was distributed across TX, IN, and NC and comprised 34 strains that were all *MAT1-1,* most likely members of a large and widely dispersed VCG.

Although strains belonging to the same VCG were grouped together in phylogenies reflecting their common ancestry and close similarity, they also showed evidence of recombination in their evolutionary histories. For example, recombination was detected in the immediate common ancestor of VCG 17 (IC243, IC244; high conflict in chromosomes 7 and 8) and VCG 6 (IC229, IC230; high conflict in chromosomes 2 and 3) (Figs. S5A, S6; Table S1). Other VCGs were more clonal in their immediate common ancestor such as VCG 5 (IC225, IC226) but there was evidence of recombination one node back which included strain IC7963 (Figs. S5A, S6). There was high conflict (red nodes) across all chromosomes in the deepest branches of the pre-application (Fig. S5A, S6) phylogenies indicating a history of extensive recombination giving rise to the sampled strains; however, there was also the hallmark of recent recombination. For example, IN strain IC7086 in lineage IC clearly inherited chromosome 2 via recombination with a strain in lineage IB (Fig. S6) and inter-lineage recombination was observed with TX strain IC6338, a full-cluster strain in lineage IB (Fig. S7). Nodal support values showed that IC6338 grouped with strains in lineage IC with strong bootstrap support (>70%) in chromosomes 3 and 5 but grouped with strains in lineage IB on other chromosomes (Fig. S7).

Major lineage expansion within lineage IB was observed within a large clade including IC201 strain (=Afla-Guard) in post 1-year treated fields that showed low conflict across all chromosomes (Figs. S5C, S8). Although strains within this clade were very similar in sequence genome-wide, there was a random distribution of both mating types (Table 3), and sampled strains were missing the entire aflatoxin gene cluster. These results are consistent with sexual recombination of closely related indigenous strains with the introduced Afla-Guard biocontrol strain in the post 1-year plots which was not observed in the pre-application fields (Fig. S6). A similar clade that was highly clonal and predominantly *MAT1-1* was observed in lineage IC which did not include IC1179 (=AF36); the latter was in a clade with isolates showing evidence of recombination (red nodes) and a mix of both mating types (Fig. S8). Similar patterns of clonality and recombination were observed in the post 3-years Afla-Guard treated fields in TX (Fig. S5D). While strains in terminal nodes showed evidence of shuffling across two or more chromosomes, lineage structure was largely maintained across chromosomal phylogenies except for chromosome 5 where lineage IC was nested within IB. The loss of lineage resolution in chromosome 5 suggests that gene flow was sufficiently high to obscure lineage boundaries (Fig. S9). This was also observed in the post 1-year treated fields (Fig. S8) but not in the post 3-months (Fig. S7) or the pre-application fields (Fig. S6).

### Chromosomal Linkage Disequilibrium

The release of biocontrol agents into field populations increased opportunities for sexual recombination with indigenous strains and lineages and thereby altered the magnitude of LD in populations (Fig. S10; Table S17). For example, for chromosome 1 in lineages IB and IC, mean *r^2^* values were highest post 3-months (IB, *r^2^*=0.300; IC, *r^2^*=0.212) than in the pre-application fields (IB, *r^2^*=0.103; IC, *r^2^*=0.169) and lowest in the post 1-year samples (IB, *r^2^* =0.053; IC, *r^2^*=0.068). This translates to the strongest LD in the post 3- month period where clonality was a dominant signature in populations and the weakest LD post 1-year, which is consistent with sexual reproduction breaking down chromosomal LD structure. Similar trends were observed over all chromosomes and sampling periods (Table S17). As more time elapsed from the initial application of biocontrol, populations returned to equilibrium levels of random mating, as exemplified in the post 3-years TX commercial fields. For example, the LD structure of chromosome 1 in the post 3-year TX field plots was very similar to the pre-application fields (IB, *r^2^*=0.120; IC, *r^2^*=0.109). This trend was observed across all chromosomes (Table S17).

### Aflatoxin Production

Quantification of aflatoxin concentrations for representative pre-application isolates from IB and IC lineages showed that overall mean toxin concentrations were lower for lineage IB strains (0.40 µg/mL; SD 0.79) compared to lineage IC (37 µg/mL; SD 19) (Table S18); Kolmogorov-Smirnov tests based on cumulative distribution functions showed that this difference was significant (*P* < 0.05) (Table S19). There was a significant decrease in lineage IB mean aflatoxin concentrations (*P* < 0.05) from the pre-application (0.40 µg/mL; SD 0.79) to the post 3-month period (0.06 µg/mL; SD 0.05); similarly, mean toxin concentrations in lineage IC decreased from the pre-application (37 µg/mL; SD 19) to post 3-months (18 µg/mL; SD 18) sampling periods, but this difference was not significant (*P* = 0.56). One year after biocontrol application mean toxin concentrations between lineages IB (1 µg/mL; SD 1) and IC (57 µg/mL; SD 57) were not significantly different from each other (*P* = 0.08). By contrast, mean toxin concentrations between lineages IB (0.17 µg/mL; SD 0.18) and IC (9 µg/mL; SD 13) in the post 3-year samples (Table S18) were significantly (*P* < 0.01) different, with 80% of the isolates in lineage IC having a low concentration of B_1_ aflatoxins (<20 µg/mL) (Table S19). There was very good agreement in mean aflatoxin concentrations that were analyzed using HPLC-fluorescence and LC-MS detection methods (data not shown).

## DISCUSSION

We conducted replicated field experiments across four states and performed genotyping by sequencing on corn and kernel isolates sampled from native populations before a one-time application of Afla-Guard and AF36 biocontrol agents, and after three months and one year later. We also examined commercial corn fields in TX which demonstrated consistently low aflatoxin concentrations three years after Afla-Guard was applied. This study has three major findings: 1) *A. flavus* field populations, before and after the application of biocontrols, are structured by evolutionary lineages IB and IC. After biocontrols were applied, lineage frequency distributions in TX were more significantly skewed to IB whereas in NC and AR they were skewed to IC. 2) *A. flavus* gene flow is asymmetric and predominantly from lineage IB into IC. Migration rates from lineage IB into IC were higher in fields treated with the Afla-Guard biocontrol agent, and gene flow was 6X higher in TX than in NC. Higher gene flow increases recombination and breaks down linkage disequilibrium resulting in higher genetic diversity, larger effective population sizes and lower mean aflatoxin B_1_ concentrations in lineage IC. 3) *A. flavus* biocontrol can lead to a sustained reduction of aflatoxin contamination. More than 90% of the isolates in the commercial cornfields in TX that were treated with the Afla-Guard biocontrol agent were from lineage IB and there was evidence of ongoing and significant gene flow from lineage IB into IC. Here, we provide evidence of gene flow and sexual recombination as the two most important forces driving diversification in *A. flavus* field populations. We show that the fate of introduced biocontrol strains is largely determined by the magnitude and direction of these two forces.

### Clonality and Recombination

Both clonality and recombination structure *A. flavus* populations (Drott *et al*. 2020; Geiser *et al*. 1998; Moore *et al*. 2017; Moore *et al*. 2009; Olarte *et al*. 2012). Clonality predominated in the Afla-Guard clade in the pre-application fields across all 4 states with 16 out of 23 strains having the same *MAT1-2* mating type, missing the entire aflatoxin cluster, and sharing a very recent common ancestor as indicated by the very short branches separating strains in the clade (Fig. S5A). Similarly, clonality was observed in lineage IC with the largest clade comprising 34 strains that were all *MAT1-1* with a full aflatoxin gene cluster; however, the AF36 biocontrol strain was in a different clade with evidence of recent clonality in the most terminal nodes (i.e., phylogenetic concordance across all chromosomes) and longer interior branches indicative of a history of recombination. Three months after biocontrol application Afla-Guard was the dominant biocontrol agent isolated from the untreated, Afla-Guard and AF36 treated plots across all four states; AF36 was not isolated from the Afla-Guard treated plots (Fig. S5B). The spread of biocontrol agents between treatment plots is not an uncommon occurrence (Kinyungu *et al*. 2019) and can be attributed primarily to their increased dispersibility, as well as aggressiveness and persistence in soil. Biocontrols are reported to persist in soil after one year (Ehrlich 2014; Lewis *et al*. 2019), most likely in the form of conidia which are far more prevalent than sclerotia (Wicklow *et al*. 1993).

Our examination of phylogenetic congruence provided evidence of sexual recombination in as little as 3-months after biocontrol applications. For example, there were four TX kernel isolates (IC6346, IC6340, IC6341, IC6345) sampled from the AF36 treated plots that were all *MAT1-1*, had full aflatoxin clusters, and shared a most recent common ancestor with Afla-Guard (Figs. S5B, S7). As expected, there was evidence of high phylogenetic conflict (red node) in the common ancestor of these strains in chromosome 3 that harbors the aflatoxin cluster (Fig. S7). This indicates that three months after biocontrols were applied, the Afla-Guard strain from either the applied biocontrol product or a clonal derivative recombined with native lineage IC strains in those plots. Fertilization most likely occurred in the soil where the spores of the Afla-Guard strain fertilized existing sclerotia of a native IC strain, formed fertile ascocarps, and released ascospores from disintegrated sclerotia, which in turn colonized corn kernels. Phylogenetic congruence also identified recombination in a major expansion of lineage IB which included Afla-Guard one year after biocontrol application (Figs. S5C, S8). Across each state there was evidence of random mating (Table 3) of the Afla-Guard strain with native lineage IB strains that were very similar genetically and as a result showed very little phylogenetic conflict (Fig. S5). Similarly, there was evidence of recombination of AF36 with native strains (Fig. S8). Previous work showed that AF36 is a putative recombinant with a toxin-producing strain (NRRL 29507) in lineage IC (Abbas *et al*. 2011a) and unlike Afla-Guard, AF36 is infrequently recovered from treated plots (Lewis *et al*. 2019).

Only five haplotypes (H1, H140, H22, H229, H240) were shared across the pre-application, post 3-months, and post 1-year sampling periods and there was a high proportion of private haplotypes (96%) across all states (Table S4), suggesting a high level of recent genetic exchange giving rise to new genotypes and minimal dispersal of haplotypes into plots from other regions. Moreover, there was also a significant clonal component in each lineage which was sufficient to maintain lineage structure over the long term, even in the presence of genetic exchange and recombination. Changes in LD across different sampling periods can be attributed to the increased recombination activity of the released biocontrol strains with native populations. Biocontrol applications resulted in similar changes in the magnitude of LD across chromosomes for both lineages IB and IC (Table S17; Fig. S10). LD was highest 3-months after biocontrols were applied showing that clonality predominates in the short term, and LD was at its lowest level one-year later as biocontrols recombined with native strains.

### VCGs and Genetic Diversity

The consistent grouping of *A. flavus* reference isolates with VCG suggests that genome-wide haplotypes are a good proxy for VCG. Putative recombinants between lineages IB and IC would have genetic backgrounds of both lineages and show evidence of phylogenetic incongruence across chromosomes. For example, IC6338 from the post 3- months TX sample contained an almost even split of both lineages IB and IC in its genetic background (Fig. 1D; Table S13) and demonstrated high conflict across chromosomes 4, 6, 8 (red nodes) in the immediate common ancestor (Figs. S5B, S7).

By contrast, IC6344 and IC6345 from TX shared a recent common ancestor, a single genetic background and low phylogenetic conflict (cyan nodes) across all chromosomes (Figs. S5B, S7). The low conflict is most likely the result of recombination with closely related members in the same lineage which does not change the lineage background but reduces phylogenetic resolution. In the case of clonality we would expect to see strong phylogenetic congruence (black nodes) across most chromosomes in the immediate common ancestor, as observed with our reference isolates that are members of the same VCG, for example, VCG 1 (IC217, IC218), VCG 4 (IC221, IC222), and VCG 5 (IC225, IC226) (Fig. S5). Populations have a high degree of clonality, which can be especially evident if sampling only VCGs with multiple representatives (Grubisha & Cotty 2010, 2015; Islam *et al*. 2018; Moral *et al*. 2020; Ortega-Beltran *et al*. 2016) but the evidence is clear in the present study and previous research (Drott *et al*. 2020; Geiser *et al*. 1998; Moore *et al*. 2013; Moore *et al*. 2009; Olarte *et al*. 2012; Olarte *et al*. 2015) that natural *A. flavus* populations comprise both frequently sampled VCGs and many singleton VCGs that contribute to genetic diversity. A sampling scheme that includes the full range of VCG diversity will show that recombination is driving genetic and mycotoxin diversity in *A. flavus*. This is exemplified by the high frequency of singleton multilocus genome-wide haplotypes (i.e., very good proxy of VCGs) that show quantitative variation in aflatoxin production, as previously reported (Horn *et al*. 1996; Olarte *et al*. 2012).

Because compatible *A. flavus* mating interactions are between parental strains that belong to different VCGs (Horn *et al*. 2016; Horn *et al*. 2009; Olarte *et al*. 2012), we expect the progeny in the first generation to be recombinants of both parents. For example, there is evidence of recombination in the immediate common ancestor of VCG 17 (IC243, 244). Subsequent amplification of this recombinant ancestor would result in a strong clonal signature across the entire genome in descendant strains, such that all isolates within the VCG would be of the same mating type and show limited evidence of phylogenetic incongruence. The incongruence, if present, could be indicative of parasexuality and our inability to detect it does not rule it out. Because isolates in lineage IB are genetically very similar to each other and gene flow is predominantly from IB into IC, many recombination events within lineage IB are undetectable but clearly occurring since populations have attained mating type equilibrium (Table 3; Fig. S5).

### Magnitude and Direction of Inter-lineage Introgression

One year after biocontrol application, treated plots in TX, NC and AR showed a significant change in the relative frequency of each lineage (i.e., lineage skew) after clone correction, with lineage IB being the dominant lineage in TX and lineage IC predominating in NC and AR (Table 4). A similar skew was observed in untreated plots prior to biocontrol applications in NC and AR and one year later in the untreated plots in TX and NC. Because field plots did not have a previous history of biocontrol applications, this suggests that an *A. flavus* lineage skew can occur naturally in field populations. Moreover, in the post 1-year untreated plots in TX, we cannot rule out that the weakly significant (0.01 < *P* < 0.05) difference in lineage frequencies is the result of cross-contamination from dispersal of biocontrol strains among plots in a field. For example, the same lineage skew was observed in NC and TX in the untreated plots and plots treated with either Afla-Guard or AF36, and in the soil and kernel populations (Table S8). This does not compromise the results because plots treated with biocontrols showed more significant (*P* < 0.01) asymmetry in lineage frequencies and gene flow estimates were significantly higher and predominantly asymmetric in the treated plots compared to the untreated.

Results from IMa3 runs showed that gene flow, when present, is predominantly from lineage IB to IC. This is supported by the limited structure observed in lineage IB and the highly structured populations in lineage IC that harbor varying proportions of IB genetic clusters (Figs 2,3; Tables S12-S15). This was also reflected in nucleotide diversity estimates which were consistently higher in lineage IC than in IB (Table S11). The signature of admixture between lineages IB and IC was also evident in PCA plots with many isolates in the post 3-months and post 3-years sampling periods occupying intermediate positions in principal component space (Fig. S1). These observations are consistent with results from Drott and coworkers (Drott *et al*. 2020). They reported the existence of three distinct *A. flavus* evolutionary linages referred to as populations A, B, and C that differed in aflatoxin production, frequency of recombination, and genetic diversity. Relatively frequent sex was found in population A with characteristics that are very similar to lineage IC; population B produced significantly less aflatoxin and comprised many isolates with deletions in the aflatoxin cluster which is the hallmark of lineage IB; population C was mostly nonaflatoxigenic and sister to population A with affinities to both lineage IB and IC.

The IMa3 program was used to quantify the magnitude and direction of introgression between lineages IB and IC. Since introgression was asymmetrical from IB into IC, larger migration estimates translated to greater genetic admixture of isolates in the receiving population. Migration rate parameters did not differ significantly from zero in the pre-application and post 1-year (untreated) fields in TX. Significant and high unidirectional gene flow in the direction of IC was detected in the post 3-months period (0.494; *P* < 0.05) with even higher rates of introgression (0.957; *P* < 0.001) one year after biocontrol application (Fig. 3). Migration rate values greater than 0.5 are considered high in simulation studies (Hey 2005). The effective population size of the IC lineage in post 1-year (17,700) was double, compared to post 3-months (9,000) field plots. While lineage IC effective population sizes were very similar in the pre-application and post 1- year (untreated) plots, lineage IB effective population size more than tripled from the pre-application (2,300) to the post 1-year (untreated) samples (7,400). The increase in effective population size can be the result of higher sexual fertility of isolates in lineage IB in TX or dispersal of Afla-Guard spores from adjacent subplots that were treated, or both. Divergence time estimates between lineages IB and IC in TX ranged from 920 YR in the pre-treatment plots to 63,000 YR in the post 1-year samples (Fig. 3), consistent with divergence times reported for three common VCGs in TX (Grubisha & Cotty 2010).

Similar patterns of gene flow were observed in the NC field plots. For example, there was an increase in lineage IC effective population size in the post 1-year treated plots (11,500) compared to the post 3-months plots (1,300), and there was also evidence of significant and moderate unidirectional gene flow (0.173; *P* < 0.001) from lineage IB into IC (Table 4; Fig. 3). Although samples sizes were small, significant asymmetrical migration from IB into IC was also observed in the pre-application fields in AR (0.172; *P* < 0.01) and even higher migration estimates were observed in IN (0.552; *P* < 0.001) (Fig. S3). Gene flow from IB into IC was further supported by the phylogenetic incongruence observed for IC7086, a strain sampled prior to biocontrol application in IN that grouped with lineage IB strains in chromosome 2 and with lineage IC strains at all other chromosomes (Fig. S6). The most striking shift was observed in the TX commercial fields 3-years after treatment with Afla-Guard, where the effective population size of lineage IC in the untreated fields (22,330) decreased to only 5,300 individuals in the treated, almost eliminating lineage IC from the Afla-Guard treated plots (Table 4; Figs. S4, S5D, S9). By comparison, lineage IB effective population size increased more than 3-fold from the untreated (827) to the Afla-Guard treated (2,800) plots, and there was evidence of significant and asymmetrical gene flow from lineage IB into IC in the untreated (1.26; *P* < 0.01) and treated (0.24; *P* < 0.001) plots (Fig. S4). The post 3-years lineage skew observed for untreated and treated plots in TX was very similar to the lineage frequencies observed in the post 1-year field plots (Table 4), which suggests that an Afla-Guard-driven population shift in TX is sustainable for at least three years.

### Lineage-Specific Differences in Fertility

Mechanistically, any difference in fertility between lineages IB and IC can result in disproportionate mating and a signature of asymmetrical introgression. For example, laboratory crosses demonstrated that when the IC278 strain from lineage IC served as the sclerotial (maternal) parent and the Afla-Guard strain from the IB lineage as conidia for fertilization, about 97% of the sclerotia were fertile compared to only 1% when the Afla-Guard strain was used as the sclerotial parent. A plausible scenario in the field is that biocontrol inoculum (conidia) fertilized sclerotia that had accumulated in soil before treatments, and in that case the biocontrol strain would be paternal. It is not known if progeny from inter-specific crosses preferentially mate with the maternal or paternal parent, which would further shift the population composition to lineage IC. This process could potentially maintain the lineage structure observed in NC and AR. Since fertile matings are possible within each lineage (Olarte *et al*. 2012) the population shift can go in the direction of either lineage depending on their relative fertilities and population sizes. The degree of fertility could be dependent on strain-specific variability in gene expression of fungal mating type pheromones and receptors which are up-regulated in high fertility crosses (Luis *et al*. 2022).

The large effective population size of the ghost in both the post 1-year untreated and treated populations (Fig. S2), suggests the presence of an unsampled sister population that shares a common ancestor with the IB lineage. A possible candidate for the sister population representing the ghost is the IA lineage, which is known to be closely related to the IB lineage and has been sampled in TX (Geiser *et al*. 2000; Horn & Dorner 1998). Lineage IA contains a mix of small (S) and large (L) sclerotial-producing strains whereas IB and IC lineages are predominantly of the L type. Alternatively, the much older common ancestor of the ghost population compared to the sampled IB and IC lineages suggests that the sister population can be a closely related species, most likely *A. parasiticus*, which has been shown to be sexually fertile with *A. flavus* strains IC308 and IC278 belonging to lineage IC but infertile with lineage IB strains (Olarte *et al*. 2015).

The observed significant differences in lineage frequencies across states may be due to either latitudinal differences in clonal population densities of lineages or fertility differences among lineages such that lineage IB is more abundant and fertile in southern subtropical regions and IC in the northern more temperate regions. This suggests that *A. flavus* biocontrol strains that belong to lineage IB would be more effective as biocontrol agents in Texas since there are more opportunities for mating with compatible strains in lineage IB. Moreover, the larger population size of IB would result in more introgression of IB into IC and greatly reduce aflatoxin contamination of crops, as observed in the TX fields three years after biocontrol application. By contrast, in NC and AR, lineage IC predominates and is more genetically diverse than IB after biocontrol application; in these regions, asymmetrical gene flow from IB into IC will increase population sizes of the IC lineage.

### Biocontrol Implications

From a biocontrol perspective, the enhanced introgression of sexually compatible lineage IB strains into native populations offers the potential for sustained reductions in aflatoxin levels over subsequent generations. This suggests that the evolutionary lineage trait may be stronger than any strain-specific differences in reducing aflatoxin levels across different latitudes and environmental conditions. Even single strain formulations of lineage IB strains can persist if environmental conditions are favorable to their growth, and native populations are fertile enough with the introduced lineage IB strain to drive and maintain sexual reproduction. For example, the Afla-Guard® strain has been shown to be effective in mitigating aflatoxin contamination in Texas (Dorner 2010; Isakeit 2015) and North Carolina (Meyers *et al*. 2015; Molo *et al*. 2018).

Aflatoxin production in lineage IB was significantly lower than in lineage IC in the pre-application fields. This lineage difference in aflatoxin producing potential was also reported for native populations of *A. flavus* in the US (Drott *et al*. 2020) and Argentina (Moore *et al*. 2013). Lab experiments have shown that aflatoxin production is highly heritable and any inter-lineage mating would increase the potential for aflatoxin production in both lineages (Olarte *et al*. 2012). The higher rates of gene flow and recombination in the post 1-year samples may explain the similar aflatoxin B_1_ distributions for lineages IB and IC. Results from Kolmogorov-Smirnov tests across different sampling periods (Table S19) showed that aflatoxin distributions in the post 3-year TX fields were significantly different between lineages, and cumulative probability distributions showed that 80% of the isolates in lineage IC had a low aflatoxin concentration (<20 µg/mL). This was less than the corresponding aflatoxin concentration for 80% of the lineage IC isolates in the pre-treatment (<40 µg/mL), post 3-month (<40 µg/mL), and post 1-year (<140 µg/mL).

Although both Afla-Guard® and AF36 biocontrol products are effective in reducing aflatoxin levels in the short term (post 3-months), their efficacy in the long-term (post 1-year) may depend on which lineage they belong to. Because lineage IB isolates are predominantly atoxigenic, populations with a greater proportion of lineage IB strains relative to lineage IC are predicted to have more reduced aflatoxin levels. In TX fields where aflatoxin levels were consistently low (10-33 ppb) over several years, there was a larger proportion of lineage IB isolates compared to IC. Moreover, lineage IC in the TX commercial corn fields was predominantly sampled in the untreated plots (Fig. S5D).

While the TX commercial cornfields are predominantly lineage IB genetically, they are functionally a mix of low aflatoxin-producing and non-aflatoxin producing strains. Any balancing selection acting to maintain aflatoxin producers and non-producers in the population (Drott *et al*. 2017; Moore *et al*. 2009) can continue but the targets of selection are now predominantly low aflatoxin producers within lineage IB. This suggests that it might be possible to shift *A. flavus* populations to a state that is functionally and qualitatively similar to the native population but quantitatively have a much-reduced aflatoxin footprint than the native population. This has significant implications for reducing aflatoxin contamination in regions where lineage IC predominates such as in NC and AR.

## Conclusion and Perspectives

The high VCG diversity in *A. flavus* soil populations translates to a range of fertility differences among isolates of different mating types. Thus, selecting a mix of biocontrol strains from lineage IB that are of different mating types and VCGs (i.e., clonal lineages) may be a useful strategy for increasing the number of successful matings in field populations (Carbone 2021). Preliminary field results of biocontrol formulations containing a mix of sexually compatible strains in NC (Molo *et al*. 2019) and TX (Isakeit, unpublished data) show that they outperform single strain formations in reducing aflatoxin concentrations and increasing corn yields. This strategy of including strains from lineage IB that are sexually compatible may explain the increased efficacy observed in biocontrol products that include a mix of nonaflatoxigenic stains. For example, the Aflasafe® biocontrol product comprises four nonaflatoxigenic *A. flavus* strains (La3279, Og0222, Ka16127, La3304) that have either a partial deletion in the aflatoxin cluster (Ka16127 and La3304) or are completely missing the cluster (La3279 and Og0222) (Adhikari *et al*. 2016; Chang 2022; Johnson *et al*. 2018). These four strains belong to different VCGs and are phylogenetically distinct from AF36 and NRRL 3357 yet genetically very similar to *A. oryzae* (Chang 2022; Chang et al. 2021), which suggests that they are likely members of lineage IB. Although mating type was not a criterion in selecting these strains they are fortuitously representative of both mating types (*MAT1-1*, La3279 and Og0222; *MAT1-2*, Ka16127 and La3304) (Chang 2022).

When compared to single strain formulations, we expect strains that are of different mating types to greatly increase effective population sizes and result in an even larger disproportionate mating access to sexually fertile strains. Future studies will examine this possibility.

## ACKNOWLEGMENTS

This work was supported by the Agriculture and Food Research Initiative Competitive Grants Program grant no. 2013-68004-20359 from the USDA National Institute of Food and Agriculture (NIFA) and from the Novo Nordisk Foundation (INTERACT NNF19SA0059360 and CCRP NNF19SA0035476). We thank Jeffrey C. Oliver for tweaking the Hypha module of Mesquite to add grid support values to nodes in Newick trees, and Mark Miller for his assistance with use of CIPRES resources.

## DATA AVAILABILITY STATEMENT

Raw DNA sequences were deposited in the NCBI SRA database (BioProject: XXXXXX-XXXXXX). Accession numbers and associated metadata are given in Table S1.

## AUTHOR CONTRIBUTIONS

M.S.M., M.A.C. and I.C. conceived and designed the research; M.S.M., T.I., K.A.W., C.P.W., R.W.H. and B.H.B. collected soil and kernel samples; M.S.M., J.B.W., R.M.G., M.S.M., V.C., M.A.C. and I.C contributed to data analysis; J.B.W. and I.C. designed the software tools; M.S.M., R.S., R.M.G. and V.C. performed the ddRADseq; V.C., R.M.G. and O.B. performed the aflatoxin assays; M.S.M, M.A.C. and I.C. wrote the manuscript with contributions from T.I., O.B. and B.W.H. All authors gave final approval for publication.

