## Supplemental Figures for "Asymmetrical lineage introgression and recombination in populations of *Aspergillus flavus*: implications for biological control"

**Figure S1. Population structure using principal component analysis (PCA).** For each treatment time point two PCA scatter plots are shown for genome-wide variation in *A. flavus* across TX, NC, AR and IN (reference strains are from GA and AZ). The PCA cubes show the distribution of individuals based on their membership in one of two clusters inferred from the Gap statistic and overlaid with lineage (PCA cube on left) or state (PCA cube on right). The color scheme and shapes are unique for each PCA cube.

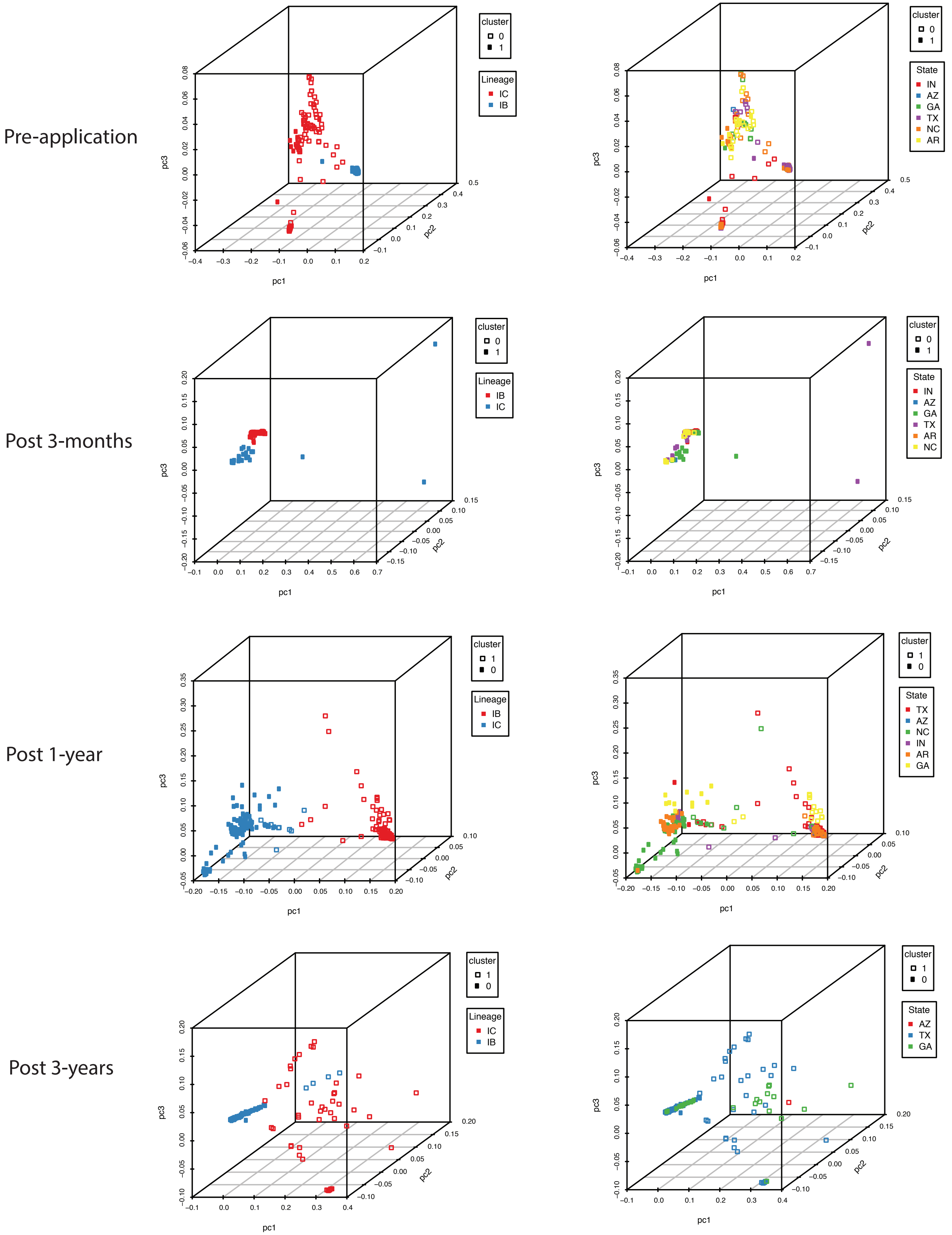
Pre-application
Post 3-months
Post 1-year
Post 3-years
cluster
0
1
Lineage
IC
IB
State
IN
AZ
GA
TX
NC
AR
pc1
pc2
pc3

**Figure S2. A schematic representation of isolation with migration for *A. flavus* populations in TX and NC including an unsampled ghost population.** The phylogeny is depicted as a hierarchical series of boxes, with ancestor boxes connecting descendant populations of lineages IB and IC, and the width of boxes proportional to the estimated $N_e$. The 95% confidence intervals for each $N_e$ value are shown as dashed lines to the right of the left side of the corresponding population box. Gray arrows to the 95% $N_e$ intervals extend on either side of the right side of each population box. Splitting times, positioned at even intervals, are depicted as solid horizontal lines, with text values on the left in units of thousand years ago (KYA). Migration arrows (in green) indicate the estimated population migration rate ($N_e m$) values from one population into another from when the populations diverged from a common ancestor. Arrows are shown only for migration rates that are statistically significant (* $p < 0.05$, ** $p < 0.01$, *** $p < 0.001$). Estimates assumed a generation time of 0.17 years and a mutation rate of $4.2 \times 10^{-11}$ per base per generation.

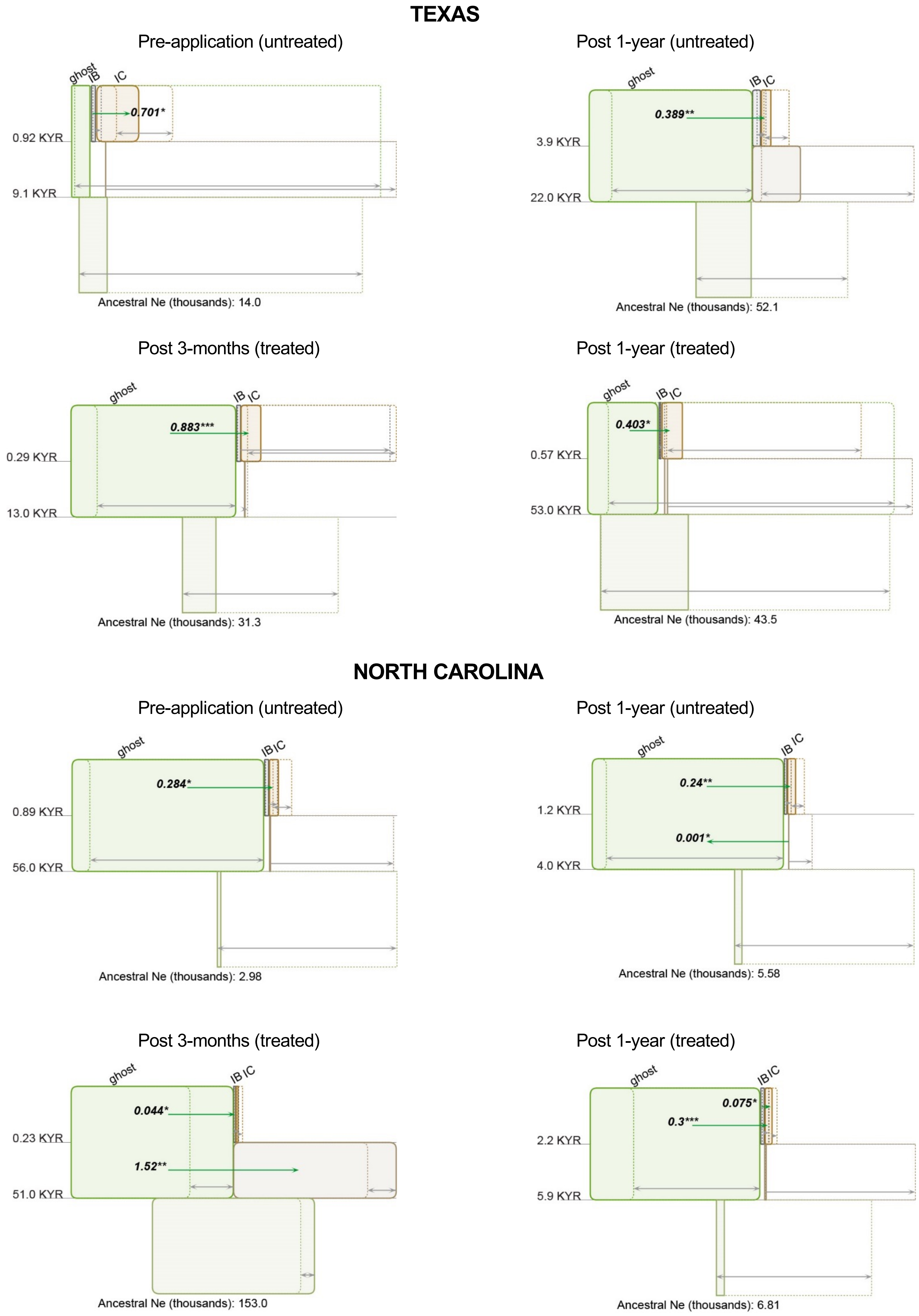
TEXAS
Pre-application (untreated)
ghost
IB
IC
0.701*
0.92 KYR
9.1 KYR
Ancestral Ne (thousands): 14.0
Post 1-year (untreated)
ghost
IB
IC
0.389**
3.9 KYR
22.0 KYR
Ancestral Ne (thousands): 52.1
Post 3-months (treated)
ghost
IB
IC
0.883***
0.29 KYR
13.0 KYR
Ancestral Ne (thousands): 31.3
Post 1-year (treated)
ghost
IB
IC
0.403*
0.57 KYR
53.0 KYR
Ancestral Ne (thousands): 43.5
NORTH CAROLINA
Pre-application (untreated)
ghost
IB
IC
0.284*
0.89 KYR
56.0 KYR
Ancestral Ne (thousands): 2.98
Post 1-year (untreated)
ghost
IB
IC
0.24**
0.001*
1.2 KYR
4.0 KYR
Ancestral Ne (thousands): 5.58
Post 3-months (treated)
ghost
IB
IC
0.044*
1.52**
0.23 KYR
51.0 KYR
Ancestral Ne (thousands): 153.0
Post 1-year (treated)
ghost
IB
IC
0.075*
0.3***
2.2 KYR
5.9 KYR
Ancestral Ne (thousands): 6.81

**Figure S3. A schematic representation of isolation with migration for *A. flavus* populations in untreated and treated field populations in AR and IN.** (A) Without a ghost population. (B) With a ghost population. The phylogeny is depicted as a hierarchical series of boxes, with ancestor boxes connecting descendant populations of lineages IB and IC, and the width of boxes proportional to the estimated $N_e$. The 95% confidence intervals for each $N_e$ value are shown as dashed lines to the right of the left side of the corresponding population box. Gray arrows to the 95% $N_e$ intervals extend on either side of the right side of each population box. Splitting times, positioned at even intervals, are depicted as solid horizontal lines, with text values on the left in units of thousand years ago (KYA). Migration arrows (in green) indicate the estimated population migration rate ($N_em$) values from one population into another from when the populations diverged from a common ancestor. Arrows are shown only for migration rates that are statistically significant (* $p < 0.05$, ** $p < 0.01$, *** $p < 0.001$). Estimates assumed a generation time of 0.17 years and a mutation rate of $4.2 \times 10^{-11}$ per base per generation.

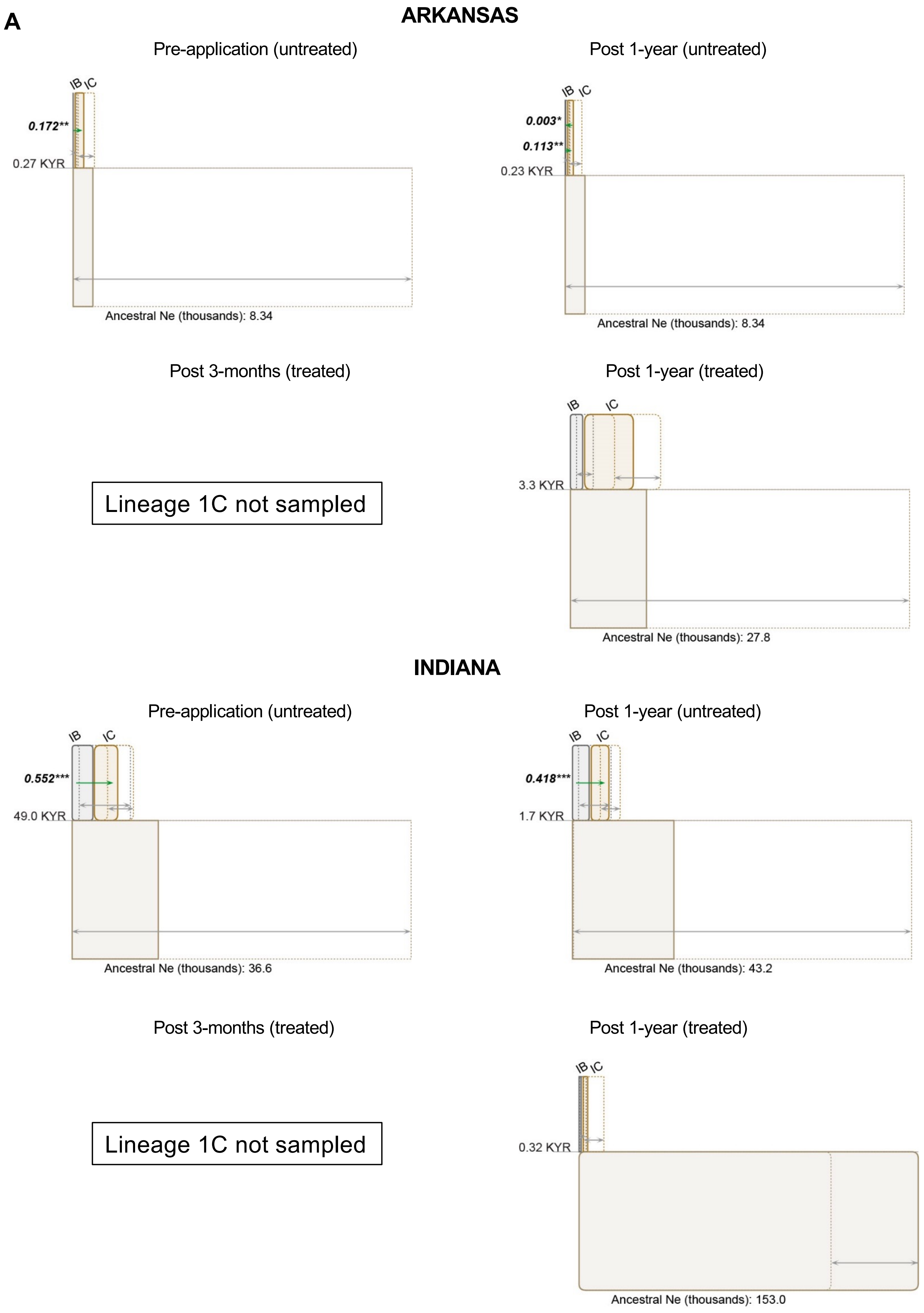
A
ARKANSAS
Pre-application (untreated)
IB
IC
0.172**
0.27 KYR
Ancestral Ne (thousands): 8.34
Post 1-year (untreated)
IB
IC
0.003*
0.113**
0.23 KYR
Ancestral Ne (thousands): 8.34
Post 3-months (treated)
Lineage 1C not sampled
Post 1-year (treated)
IB
IC
3.3 KYR
Ancestral Ne (thousands): 27.8
INDIANA
Pre-application (untreated)
IB
IC
0.552***
49.0 KYR
Ancestral Ne (thousands): 36.6
Post 1-year (untreated)
IB
IC
0.418***
1.7 KYR
Ancestral Ne (thousands): 43.2
Post 3-months (treated)
Lineage 1C not sampled
Post 1-year (treated)
IB
IC
0.32 KYR
Ancestral Ne (thousands): 153.0

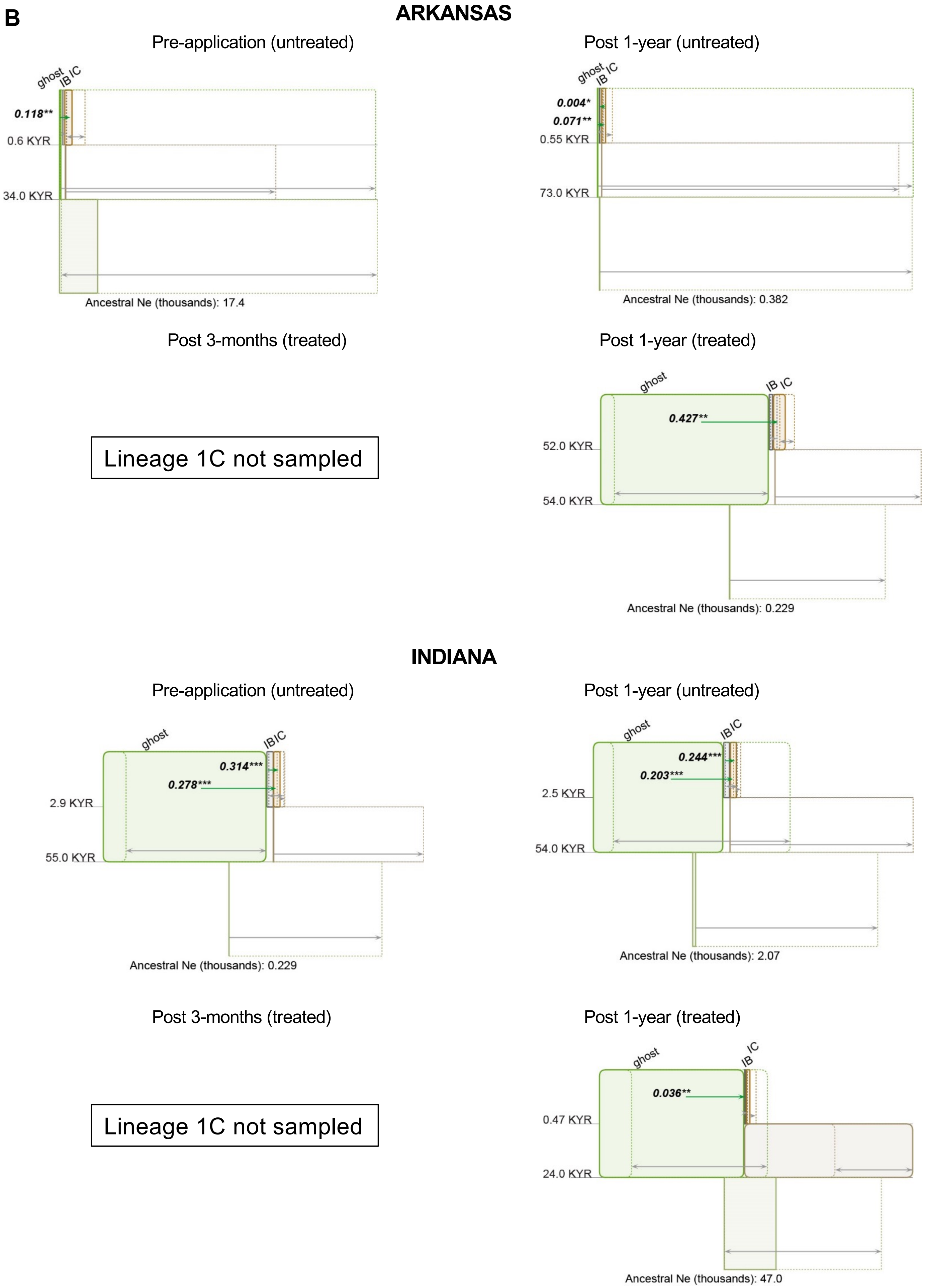
B
ARKANSAS
Pre-application (untreated)
ghost
IB
IC
0.118**
0.6 KYR
34.0 KYR
Ancestral Ne (thousands): 17.4
Post 1-year (untreated)
ghost
IB
IC
0.004*
0.071**
0.55 KYR
73.0 KYR
Ancestral Ne (thousands): 0.382
Post 3-months (treated)
Lineage 1C not sampled
Post 1-year (treated)
ghost
IB
IC
0.427**
52.0 KYR
54.0 KYR
Ancestral Ne (thousands): 0.229
INDIANA
Pre-application (untreated)
ghost
IB
IC
0.314***
0.278***
2.9 KYR
55.0 KYR
Ancestral Ne (thousands): 0.229
Post 1-year (untreated)
ghost
IB
IC
0.244***
0.203***
2.5 KYR
54.0 KYR
Ancestral Ne (thousands): 2.07
Post 3-months (treated)
Lineage 1C not sampled
Post 1-year (treated)
ghost
IB
IC
0.036**
0.47 KYR
24.0 KYR
Ancestral Ne (thousands): 47.0

**Figure S4. A schematic representation of isolation with migration for *A. flavus* populations in untreated and treated TX commercial fields**. (A) Without a ghost population. (B) With a ghost population. The phylogeny is depicted as a hierarchical series of boxes, with ancestor boxes connecting descendant populations of lineages IB and IC, and the width of boxes proportional to the estimated $N_e$. The 95% confidence intervals for each $N_e$ value are shown as dashed lines to the right of the left side of the corresponding population box. Gray arrows to the 95% $N_e$ intervals extend on either side of the right side of each population box. Splitting times, positioned at even intervals, are depicted as solid horizontal lines, with text values on the left in units of thousand years ago (KYA). Migration arrows (in green) indicate the estimated population migration rate ($N_e m$) values from one population into another from when the populations diverged from a common ancestor. Arrows are shown only for migration rates that are statistically significant (* $p < 0.05$, ** $p < 0.01$, *** $p < 0.001$). Estimates assumed a generation time of 0.17 years and a mutation rate of $4.2 \times 10^{-11}$ per base per generation.

**A**

**TEXAS Commercial**

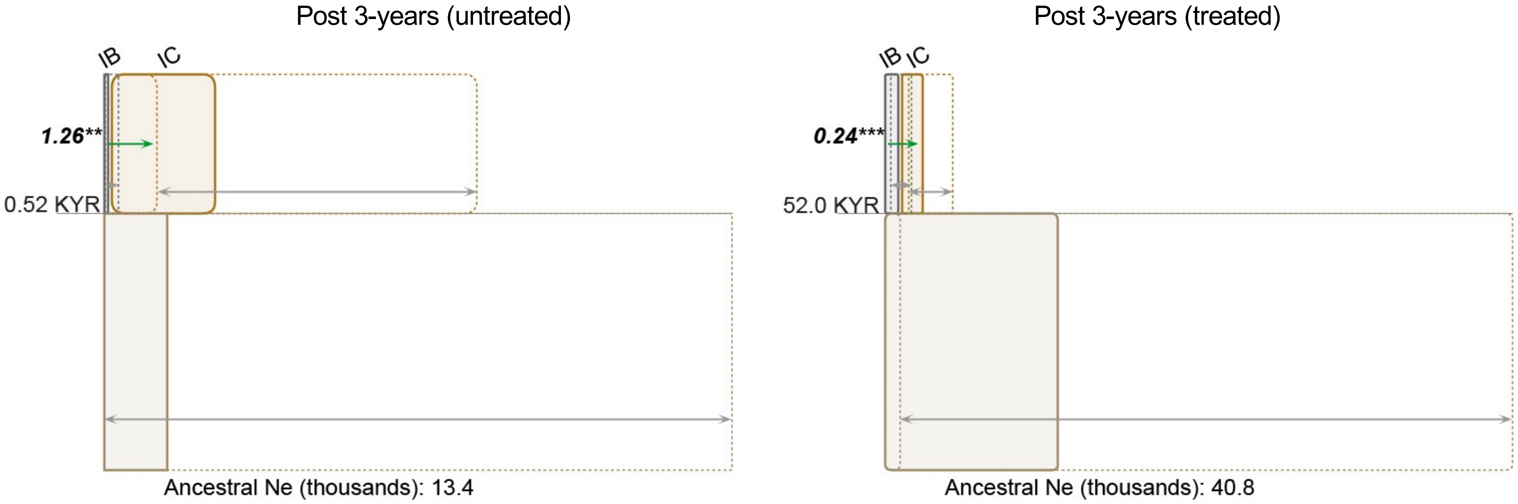


**B**

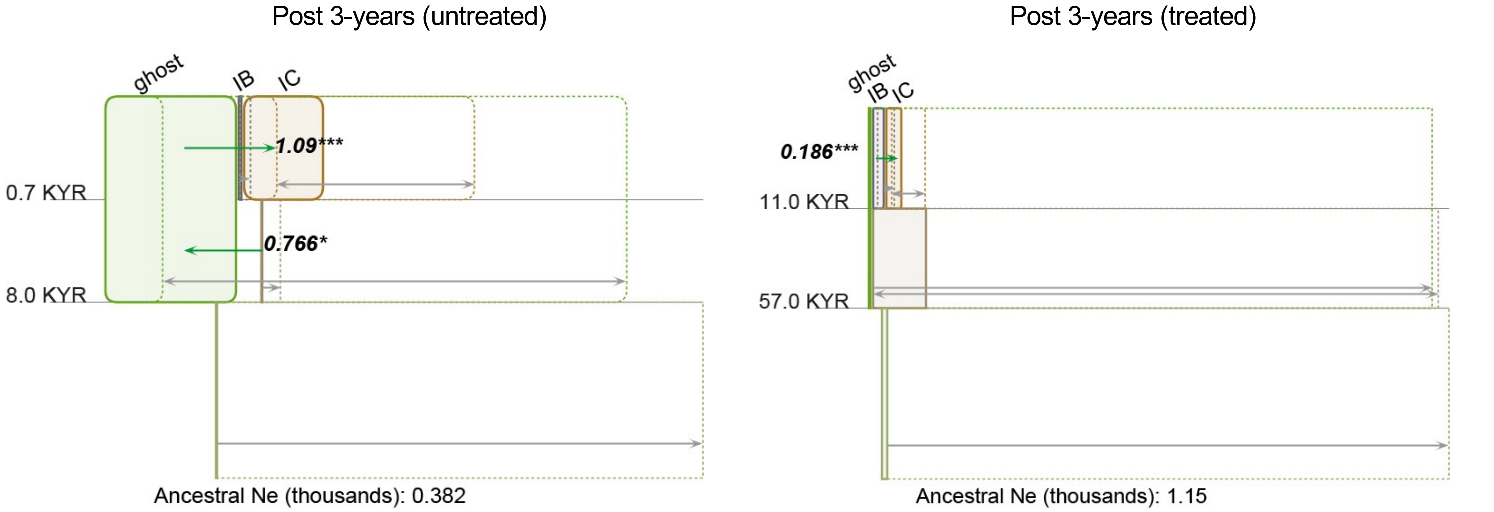

**Figure S5. Phylogenetic congruence using the Hypha module in Mesquite and displayed using T-BAS**. Phylogenetic congruence of each chromosome phylogeny compared to the total evidence display tree for the four different sampling time points (A-D) is shown using grids on node partitions. Branch lengths on the total evidence tree are drawn to scale and the scale bar is shown at the top. In each grid, bootstrap support values are displayed with each box from left to right representing one of eight chromosomes; the box on the bottom right is for the mitochondrial genome. Colors in grids represent node bipartitions that were supported at a bootstrap support value ≥70% (black color), <70% (white color), and missing or inapplicable (grey color). Phylogenetic incongruency was represented as high conflict (red color) and low conflict (cyan color). Additional attributes (lineage, state, mating type, AF cluster configuration, and substrate/treatment) are shown in columns adjacent to the strain names.

(A) Pre-application

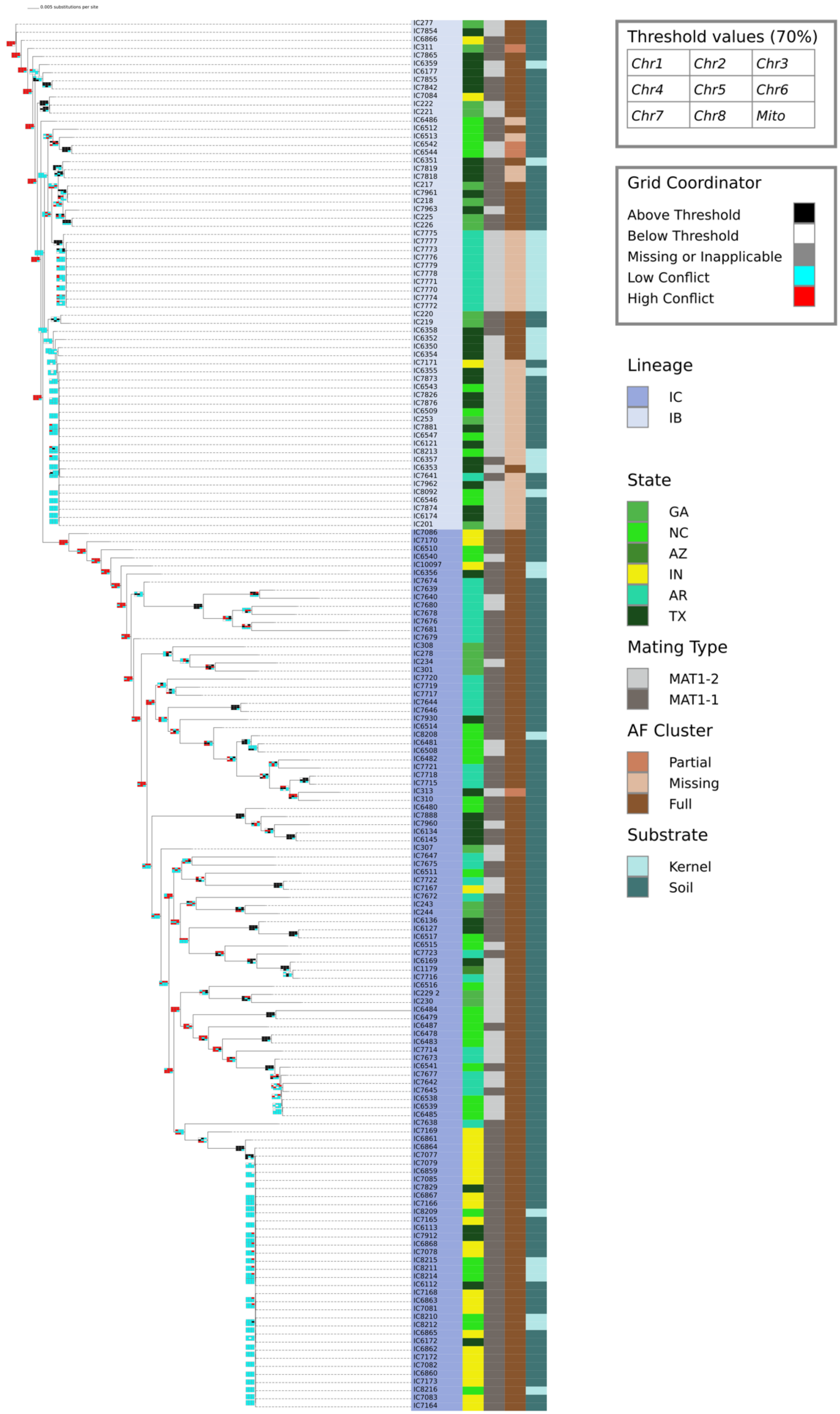

#### (B) Post 3-months

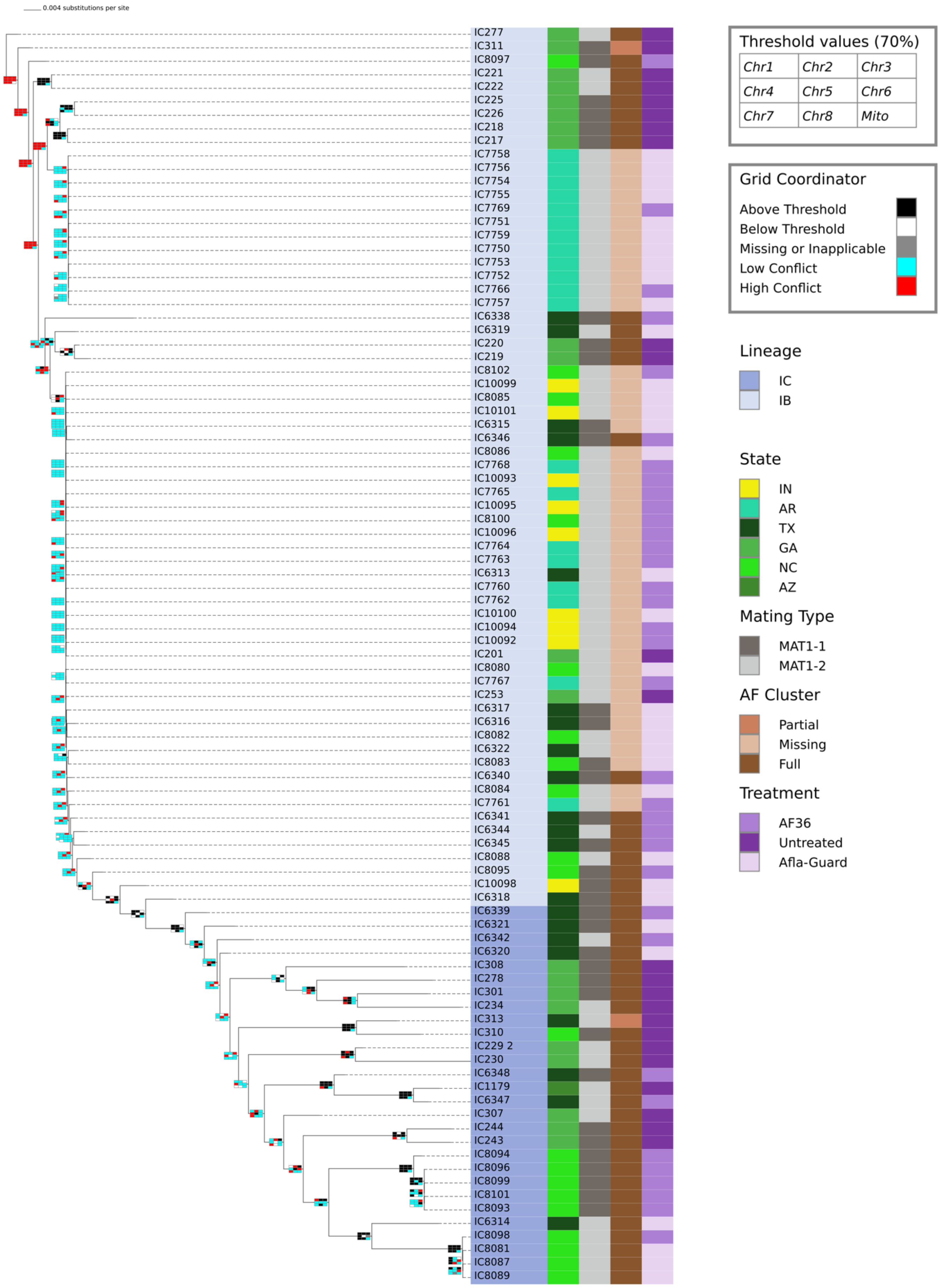

(C) Post 1-year

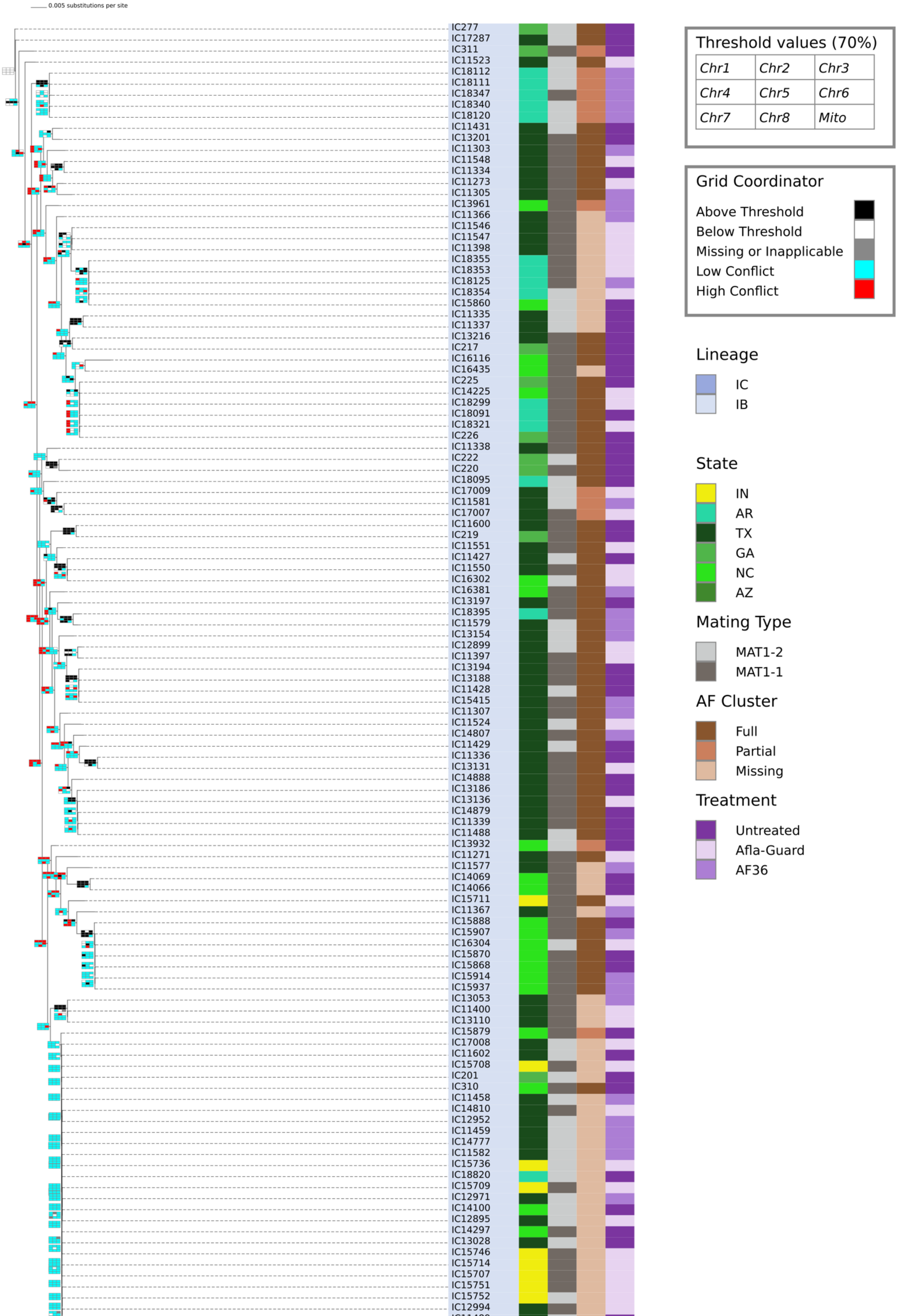

0.005 substitutions per site
IC277
IC17287
IC311
IC11523
IC18112
IC18111
IC18347
IC18340
IC18120
IC11431
IC13201
IC11303
IC11548
IC11334
IC11273
IC11305
IC13961
IC11366
IC11546
IC11547
IC11398
IC18355
IC18353
IC18125
IC18354
IC15860
IC11335
IC11337
IC13216
IC217
IC16116
IC16435
IC225
IC14225
IC18299
IC18091
IC18321
IC226
IC11338
IC222
IC220
IC18095
IC17009
IC11581
IC17007
IC11600
IC219
IC11551
IC11427
IC11550
IC16302
IC16381
IC13197
IC18395
IC11579
IC13154
IC12899
IC11397
IC13194
IC13188
IC11428
IC15415
IC11307
IC11524
IC14807
IC11429
IC11336
IC13131
IC14888
IC13186
IC13136
IC14879
IC11339
IC11488
IC13932
IC11271
IC11577
IC14069
IC14066
IC15711
IC11367
IC15888
IC15907
IC16304
IC15870
IC15868
IC15914
IC15937
IC13053
IC11400
IC13110
IC15879
IC17008
IC11602
IC15708
IC201
IC310
IC11458
IC14810
IC12952
IC11459
IC14777
IC11582
IC15736
IC18820
IC15709
IC12971
IC14100
IC12895
IC14297
IC13028
IC15746
IC15714
IC15707
IC15751
IC15752
IC12994
Threshold values (70%)
Chr1 Chr2 Chr3
Chr4 Chr5 Chr6
Chr7 Chr8 Mito
Grid Coordinator
Above Threshold
Below Threshold
Missing or Inapplicable
Low Conflict
High Conflict
Lineage
IC
IB
State
IN
AR
TX
GA
NC
AZ
Mating Type
MAT1-2
MAT1-1
AF Cluster
Full
Partial
Missing
Treatment
Untreated
Afla-Guard
AF36

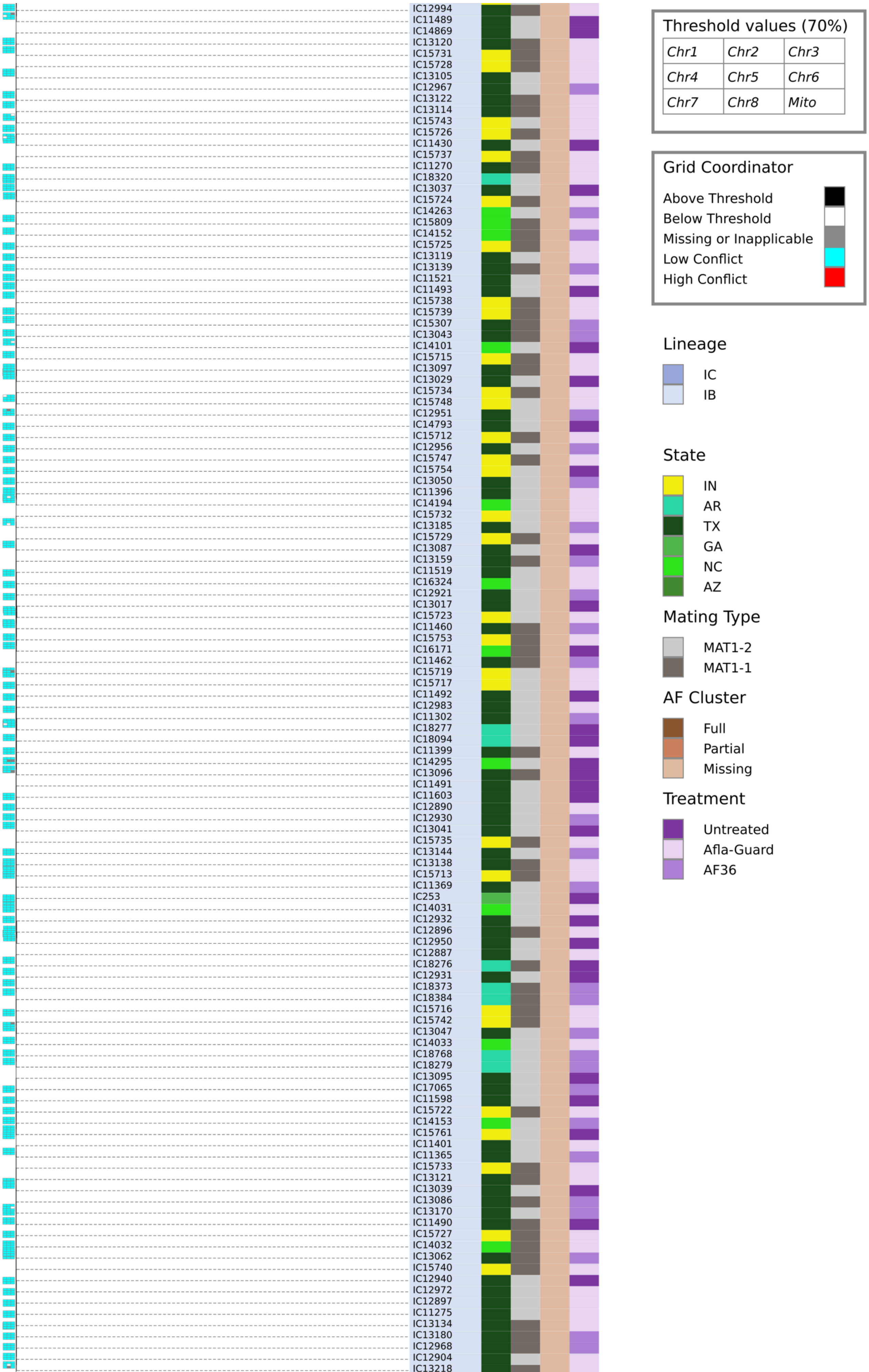
IC12994
IC11489
IC14869
IC13120
IC15731
IC15728
IC13105
IC12967
IC13122
IC13114
IC15743
IC15726
IC11430
IC15737
IC11270
IC18320
IC13037
IC15724
IC14263
IC15809
IC14152
IC15725
IC13119
IC13139
IC11521
IC11493
IC15738
IC15739
IC15307
IC13043
IC14101
IC15715
IC13097
IC13029
IC15734
IC15748
IC12951
IC14793
IC15712
IC12956
IC15747
IC15754
IC13050
IC11396
IC14194
IC15732
IC13185
IC15729
IC13087
IC13159
IC11519
IC16324
IC12921
IC13017
IC15723
IC11460
IC15753
IC16171
IC11462
IC15719
IC15717
IC11492
IC12983
IC11302
IC18277
IC18094
IC11399
IC14295
IC13096
IC11491
IC11603
IC12890
IC12930
IC13041
IC15735
IC13144
IC13138
IC15713
IC11369
IC253
IC14031
IC12932
IC12896
IC12950
IC12887
IC18276
IC12931
IC18373
IC18384
IC15716
IC15742
IC13047
IC14033
IC18768
IC18279
IC13095
IC17065
IC11598
IC15722
IC14153
IC15761
IC11401
IC11365
IC15733
IC13121
IC13039
IC13086
IC13170
IC11490
IC15727
IC14032
IC13062
IC15740
IC12940
IC12972
IC12897
IC11275
IC13134
IC13180
IC12968
IC12904
IC13218
Threshold values (70%)
Chr1
Chr2
Chr3
Chr4
Chr5
Chr6
Chr7
Chr8
Mito
Grid Coordinator
Above Threshold
Below Threshold
Missing or Inapplicable
Low Conflict
High Conflict
Lineage
IC
IB
State
IN
AR
TX
GA
NC
AZ
Mating Type
MAT1-2
MAT1-1
AF Cluster
Full
Partial
Missing
Treatment
Untreated
Afla-Guard
AF36

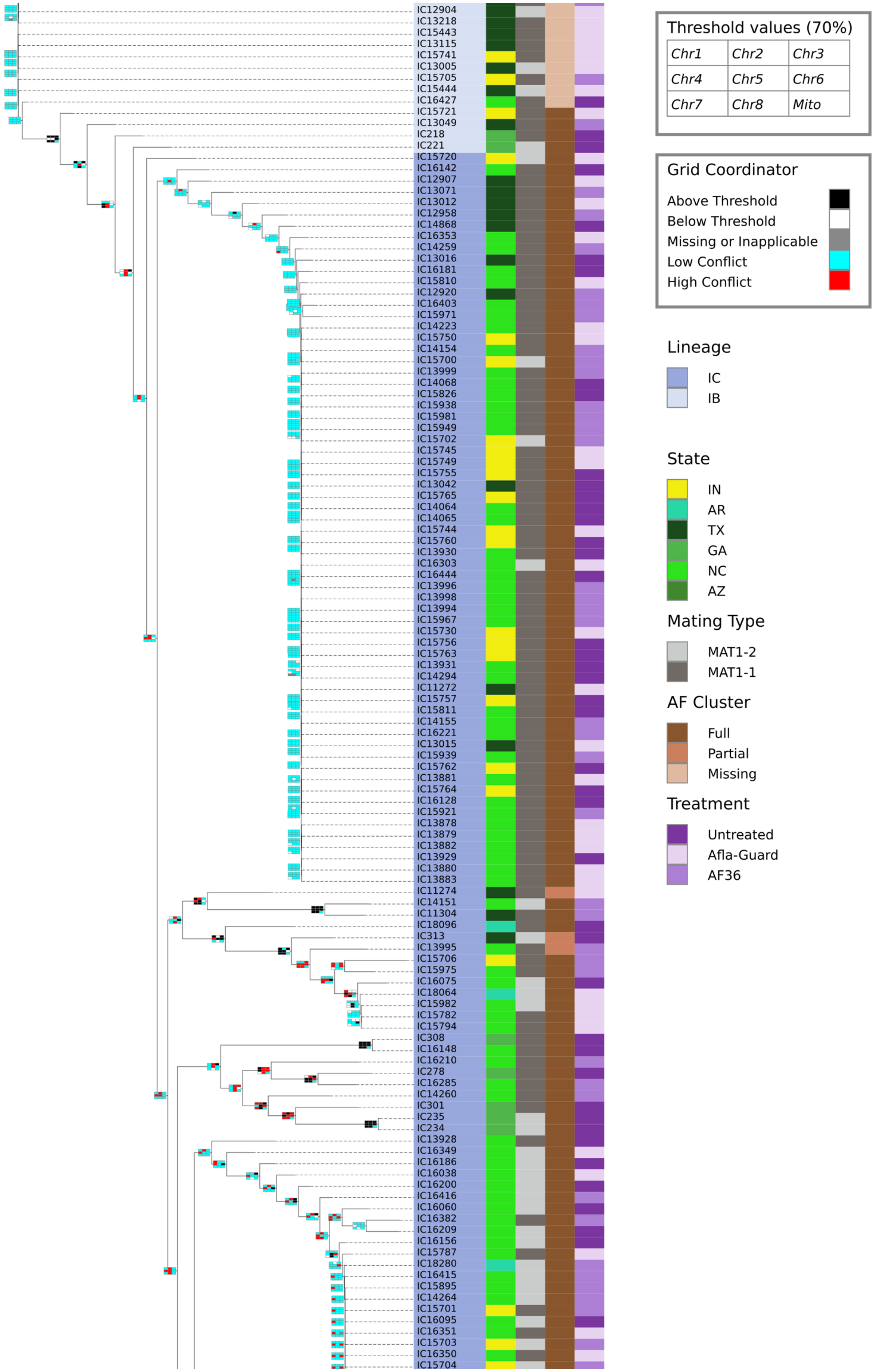
IC12904
IC13218
IC15443
IC13115
IC15741
IC13005
IC15705
IC15444
IC16427
IC15721
IC13049
IC218
IC221
IC15720
IC16142
IC12907
IC13071
IC13012
IC12958
IC14868
IC16353
IC14259
IC13016
IC16181
IC15810
IC12920
IC16403
IC15971
IC14223
IC15750
IC14154
IC15700
IC13999
IC14068
IC15826
IC15938
IC15981
IC15949
IC15702
IC15745
IC15749
IC15755
IC13042
IC15765
IC14064
IC14065
IC15744
IC15760
IC13930
IC16303
IC16444
IC13996
IC13998
IC13994
IC15967
IC15730
IC15756
IC15763
IC13931
IC14294
IC11272
IC15757
IC15811
IC14155
IC16221
IC13015
IC15939
IC15762
IC13881
IC15764
IC16128
IC15921
IC13878
IC13879
IC13882
IC13929
IC13880
IC13883
IC11274
IC14151
IC11304
IC18096
IC313
IC13995
IC15706
IC15975
IC16075
IC18064
IC15982
IC15782
IC15794
IC308
IC16148
IC16210
IC278
IC16285
IC14260
IC301
IC235
IC234
IC13928
IC16349
IC16186
IC16038
IC16200
IC16416
IC16060
IC16382
IC16209
IC16156
IC15787
IC18280
IC16415
IC15895
IC14264
IC15701
IC16095
IC16351
IC15703
IC16350
IC15704
Threshold values (70%)
Chr1
Chr2
Chr3
Chr4
Chr5
Chr6
Chr7
Chr8
Mito
Grid Coordinator
Above Threshold
Below Threshold
Missing or Inapplicable
Low Conflict
High Conflict
Lineage
IC
IB
State
IN
AR
TX
GA
NC
AZ
Mating Type
MAT1-2
MAT1-1
AF Cluster
Full
Partial
Missing
Treatment
Untreated
Afla-Guard
AF36

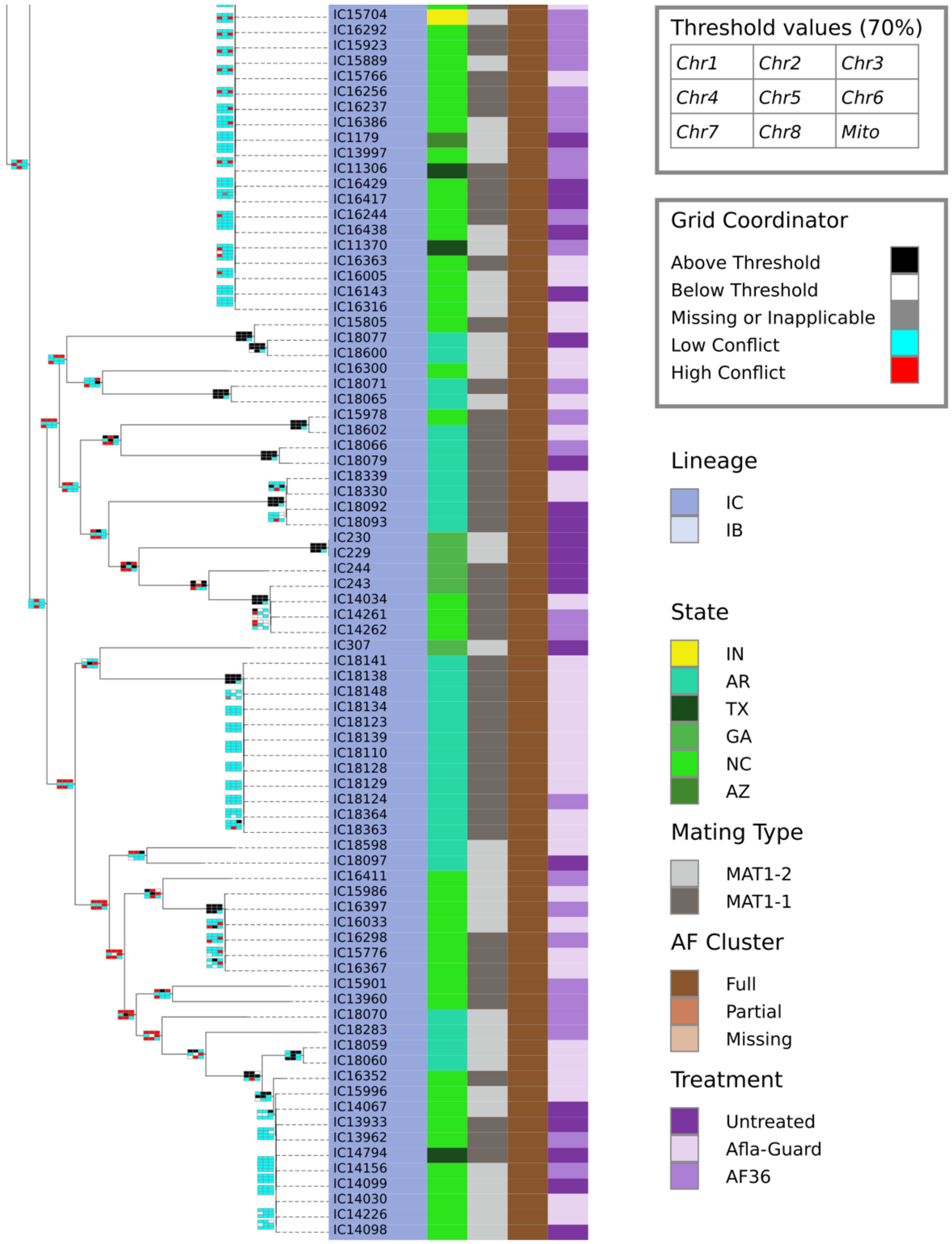
IC15704
IC16292
IC15923
IC15889
IC15766
IC16256
IC16237
IC16386
IC1179
IC13997
IC11306
IC16429
IC16417
IC16244
IC16438
IC11370
IC16363
IC16005
IC16143
IC16316
IC15805
IC18077
IC18600
IC16300
IC18071
IC18065
IC15978
IC18602
IC18066
IC18079
IC18339
IC18330
IC18092
IC18093
IC230
IC229
IC244
IC243
IC14034
IC14261
IC14262
IC307
IC18141
IC18138
IC18148
IC18134
IC18123
IC18139
IC18110
IC18128
IC18129
IC18124
IC18364
IC18363
IC18598
IC18097
IC16411
IC15986
IC16397
IC16033
IC16298
IC15776
IC16367
IC15901
IC13960
IC18070
IC18283
IC18059
IC18060
IC16352
IC15996
IC14067
IC13933
IC13962
IC14794
IC14156
IC14099
IC14030
IC14226
IC14098
Threshold values (70%)
Chr1 Chr2 Chr3
Chr4 Chr5 Chr6
Chr7 Chr8 Mito
Grid Coordinator
Above Threshold
Below Threshold
Missing or Inapplicable
Low Conflict
High Conflict
Lineage
IC
IB
State
IN
AR
TX
GA
NC
AZ
Mating Type
MAT1-2
MAT1-1
AF Cluster
Full
Partial
Missing
Treatment
Untreated
Afla-Guard
AF36

(D) Post 3-years

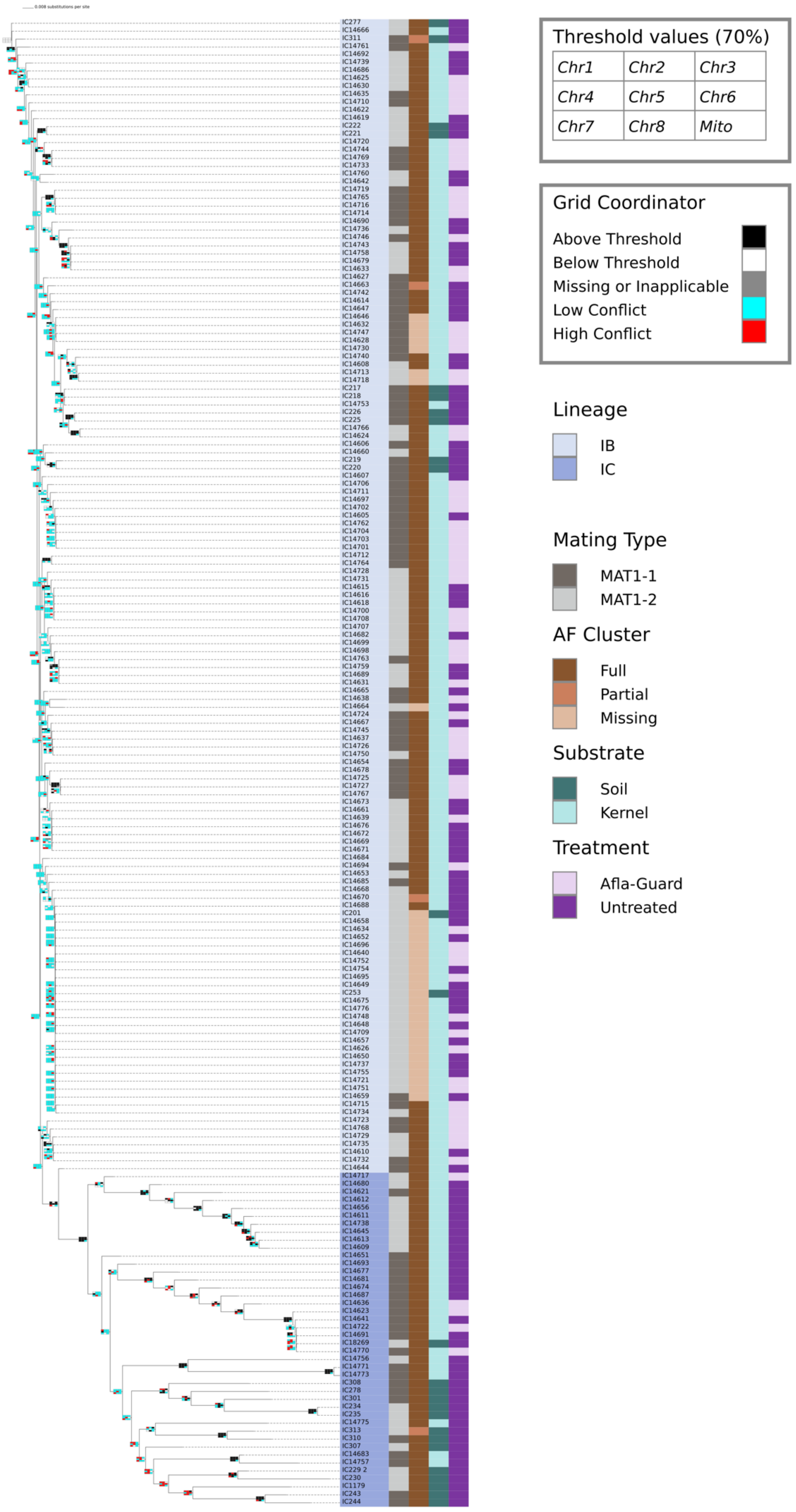

**Figure S6. Phylogenetic congruence of each chromosome and the mitochondrial genome for isolates sampled before biocontrol application**. In the total evidence display tree colors in grids represent node bipartitions that were supported at a bootstrap support value ≥70% (black color), <70% (white color), and missing or inapplicable (grey color). Phylogenetic incongruency was represented as high conflict (red color) and low conflict (cyan color). Strain names are highlighted to show their lineage membership in the total evidence tree; the mitochondrial genome has insufficient variation and poor resolution of lineage structure. The red arrows track the position of strain IC7086 sampled from IN which belongs to lineage IC; the black arrows track the position of IC6357 sampled from TX which is in lineage IB. Strain IC7086 is grouping with lineage IB strains in chromosome 2 but is placed in lineage IC on all other chromosomes with strong bootstrap support (≥90%). Strain IC6357 groups only with lineage IB strains on different chromosomes but with weak bootstrap support (<70%).

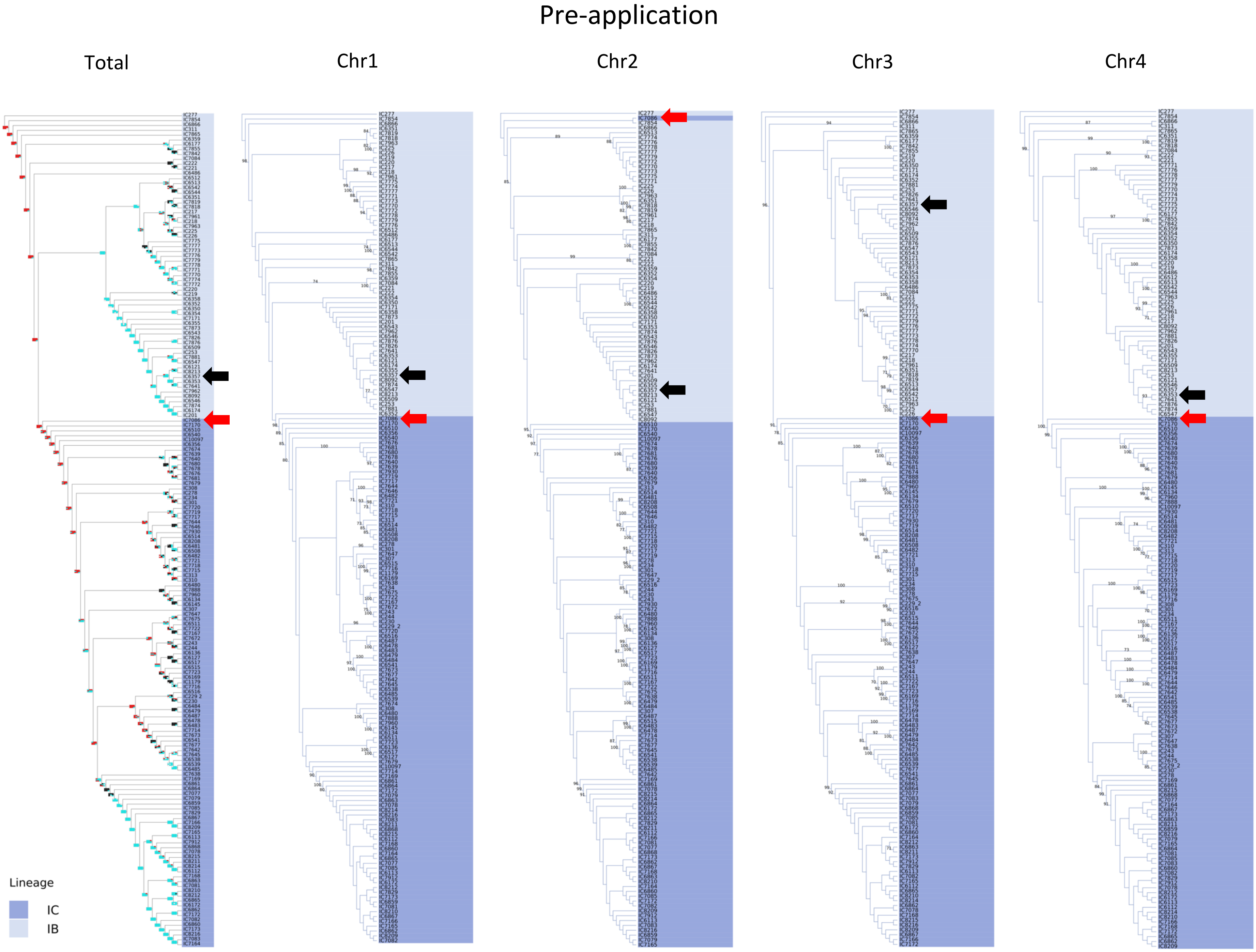
Pre-application
Total
Chr1
Chr2
Chr3
Chr4
Lineage
IC
IB

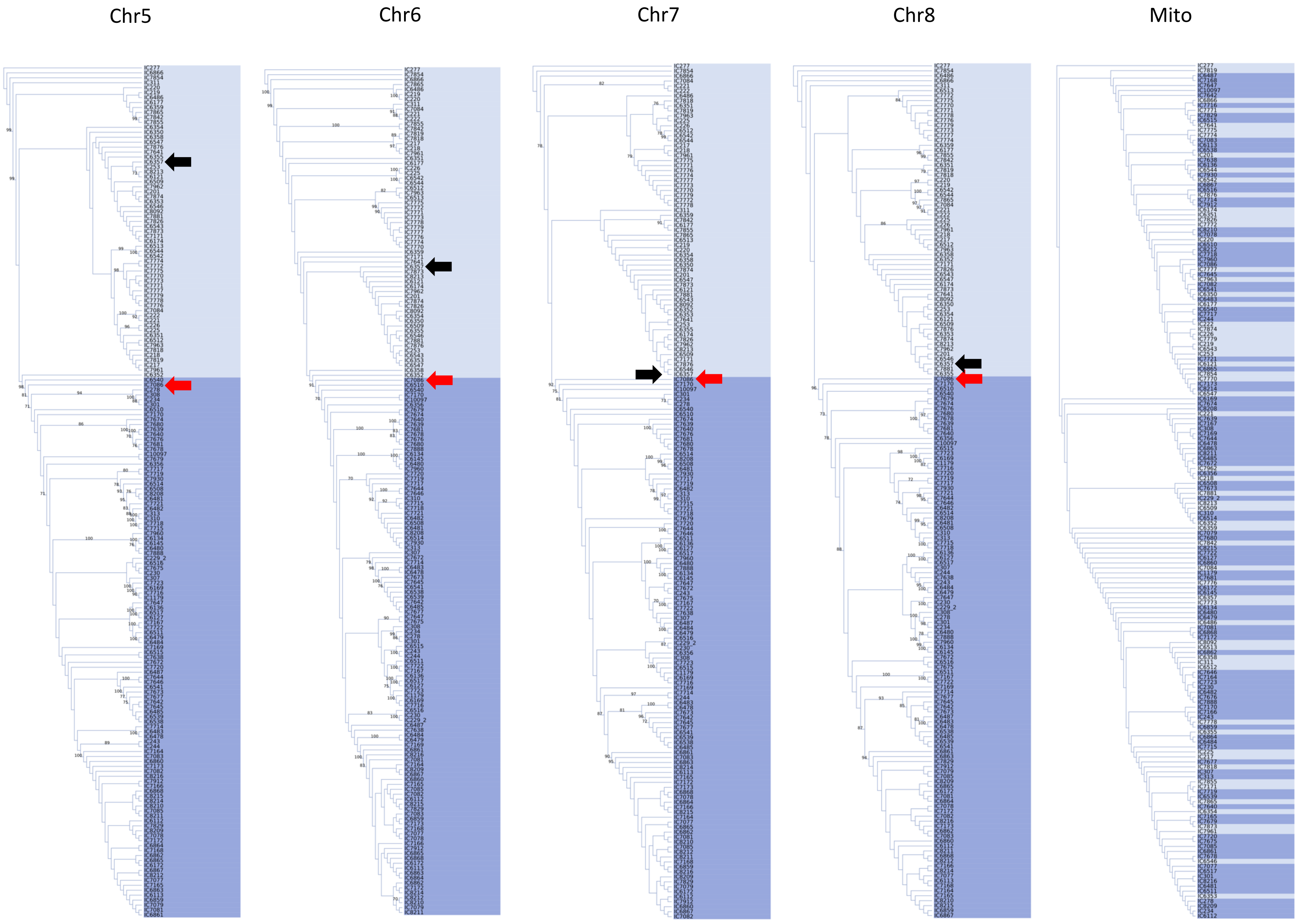
Chr5
Chr6
Chr7
Chr8
Mito

**Figure S7. Phylogenetic congruence of each chromosome and the mitochondrial genome for isolates sampled 3-months after biocontrol application.** In the total evidence display tree colors in grids represent node bipartitions that were supported at a bootstrap support value ≥70% (black color), <70% (white color), and missing or inapplicable (grey color). Phylogenetic incongruency was represented as high conflict (red color) and low conflict (cyan color). Strain names are highlighted to show their lineage membership in the total evidence tree; the mitochondrial genome has insufficient variation and poor resolution of lineage structure. The red arrows track the position of strain IC6338 sampled from TX which belongs to lineage IB in the total evidence tree. Strain IC6338 groups in lineage IB in chromosomes 1, 2, 4, 6, 7 and 8, but there is strong bootstrap support (>70%) for IC6338 grouping with strains in lineage IC in chromosomes 3 and 5.

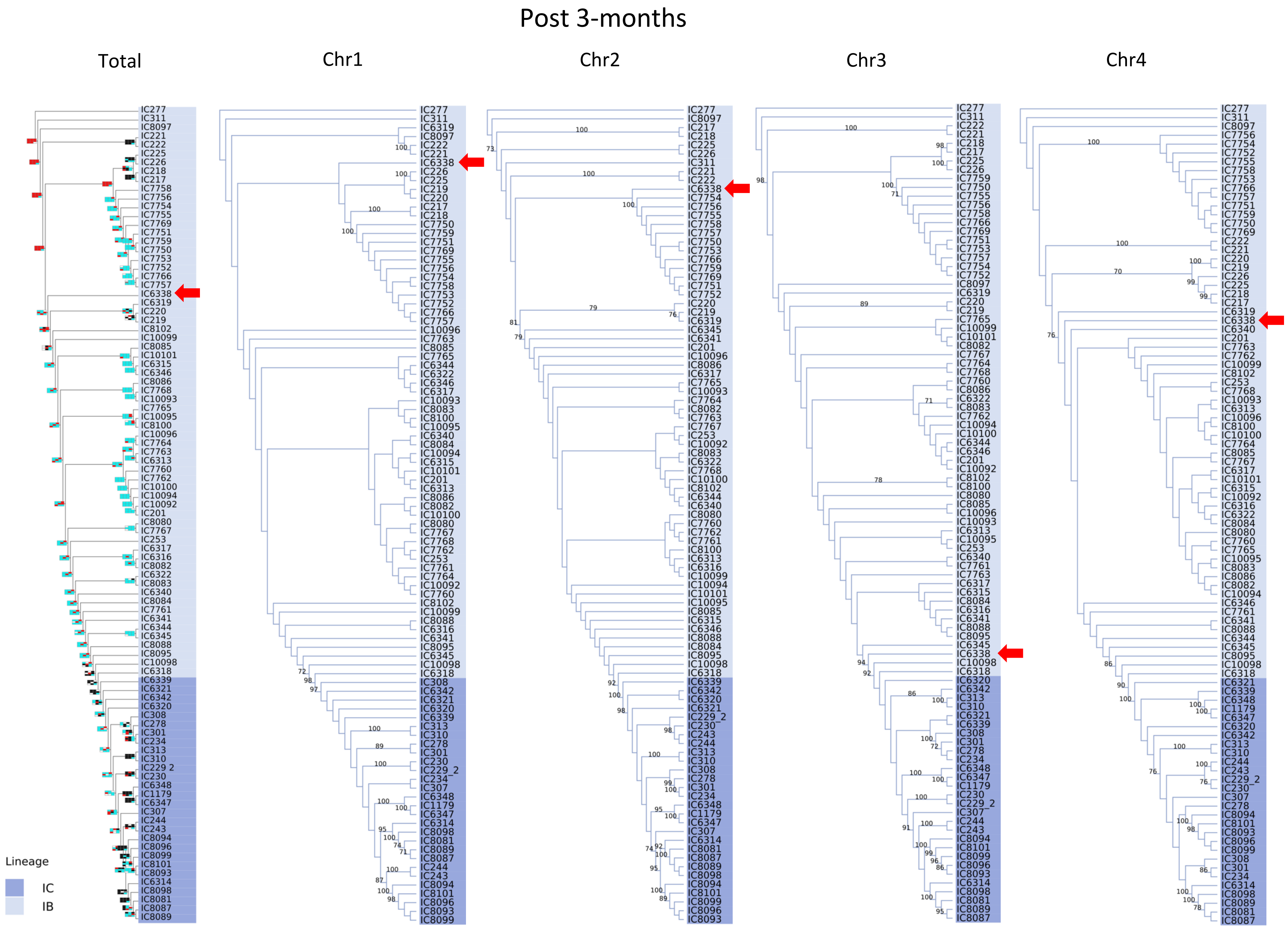
Post 3-months
Total
Chr1
Chr2
Chr3
Chr4
Lineage
IC
IB

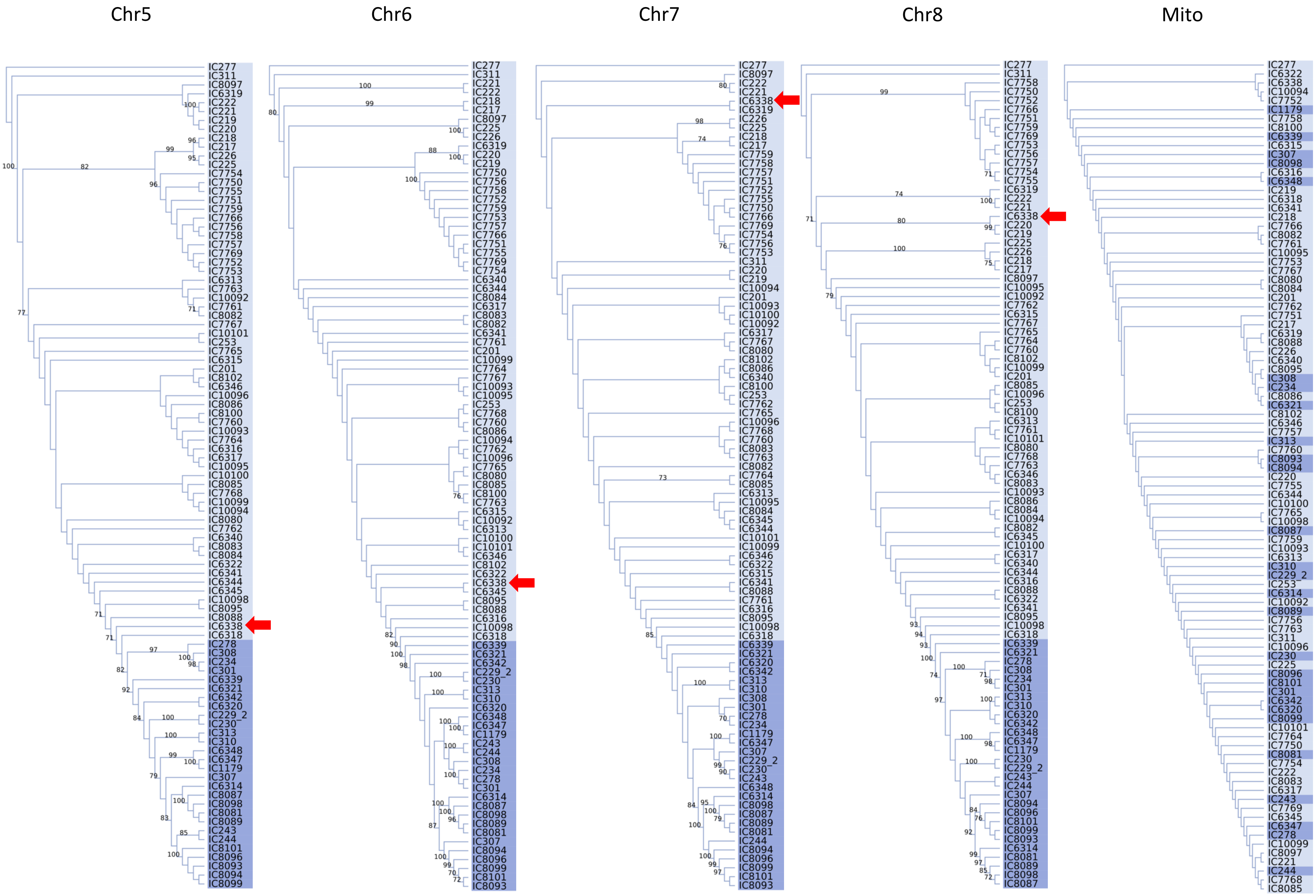
Chr5
Chr6
Chr7
Chr8
Mito
IC277 IC311 IC8097 IC6319 IC222 IC221 IC219 IC220 IC218 IC217 IC226 IC225 IC7754 IC7750 IC7755 IC7751 IC7759 IC7766 IC7756 IC7758 IC7757 IC7769 IC7752 IC7753 IC6313 IC7763 IC10092 IC7761 IC8082 IC7767 IC10101 IC253 IC7765 IC6315 IC201 IC8102 IC6346 IC10096 IC8086 IC8100 IC7760 IC10093 IC7764 IC6316 IC6317 IC10095 IC10100 IC8085 IC7768 IC10099 IC10094 IC8080 IC7762 IC6340 IC8083 IC8084 IC6322 IC6341 IC6344 IC6345 IC10098 IC8095 IC8088 IC6338 IC6318 IC278 IC308 IC234 IC301 IC6339 IC6321 IC6342 IC6320 IC229_2 IC230 IC313 IC310 IC6348 IC6347 IC1179 IC307 IC6314 IC8087 IC8098 IC8081 IC8089 IC243 IC244 IC8101 IC8096 IC8093 IC8094 IC8099
IC277 IC311 IC221 IC222 IC218 IC217 IC8097 IC225 IC226 IC6319 IC220 IC219 IC7750 IC7756 IC7758 IC7752 IC7759 IC7753 IC7757 IC7766 IC7751 IC7755 IC7769 IC7754 IC6340 IC6344 IC8084 IC6317 IC8083 IC8082 IC6341 IC7761 IC201 IC10099 IC7764 IC7767 IC10093 IC10095 IC253 IC7768 IC7760 IC8086 IC10094 IC7762 IC10096 IC7765 IC8080 IC8085 IC8100 IC7763 IC6315 IC10092 IC6313 IC10100 IC10101 IC6346 IC8102 IC6322 IC6338 IC6345 IC8095 IC8088 IC6316 IC10098 IC6318 IC6339 IC6321 IC6342 IC229_2 IC230 IC313 IC310 IC6320 IC6348 IC6347 IC1179 IC243 IC244 IC308 IC234 IC278 IC301 IC6314 IC8087 IC8098 IC8089 IC8081 IC307 IC8094 IC8096 IC8099 IC8101 IC8093
IC277 IC8097 IC222 IC221 IC6338 IC6319 IC226 IC225 IC218 IC217 IC7759 IC7758 IC7757 IC7751 IC7752 IC7755 IC7750 IC7766 IC7769 IC7754 IC7756 IC7753 IC311 IC220 IC219 IC10094 IC201 IC10093 IC10100 IC10092 IC6317 IC7767 IC8080 IC8102 IC8086 IC6340 IC8100 IC253 IC7762 IC7765 IC10096 IC7768 IC7760 IC8083 IC7763 IC8082 IC7764 IC8085 IC6313 IC10095 IC8084 IC6345 IC6344 IC10101 IC10099 IC6346 IC6322 IC6315 IC6341 IC8088 IC7761 IC6316 IC8095 IC10098 IC6318 IC6339 IC6321 IC6320 IC6342 IC313 IC310 IC308 IC301 IC278 IC234 IC1179 IC6347 IC307 IC229_2 IC230 IC243 IC6348 IC6314 IC8098 IC8087 IC8089 IC8081 IC244 IC8094 IC8096 IC8099 IC8101 IC8093
IC277 IC311 IC7758 IC7750 IC7752 IC7766 IC7751 IC7759 IC7769 IC7753 IC7756 IC7757 IC7754 IC7755 IC6319 IC222 IC221 IC6338 IC220 IC219 IC225 IC226 IC218 IC217 IC8097 IC10095 IC10092 IC7762 IC6315 IC7767 IC7765 IC7764 IC7760 IC8102 IC10099 IC201 IC8085 IC10096 IC253 IC8100 IC6313 IC7761 IC10101 IC8080 IC7768 IC7763 IC6346 IC8083 IC10093 IC8086 IC8084 IC10094 IC8082 IC6345 IC10100 IC6317 IC6340 IC6344 IC6316 IC8088 IC6322 IC6341 IC8095 IC10098 IC6318 IC6339 IC6321 IC278 IC308 IC234 IC301 IC313 IC310 IC6320 IC6342 IC6348 IC6347 IC1179 IC230 IC229_2 IC243 IC244 IC307 IC8094 IC8096 IC8101 IC8099 IC8093 IC6314 IC8081 IC8089 IC8098 IC8087
IC277 IC6322 IC6338 IC10094 IC7752 IC1179 IC7758 IC8100 IC6339 IC6315 IC307 IC8098 IC6316 IC6348 IC219 IC6318 IC6341 IC218 IC7766 IC8082 IC7761 IC10095 IC7753 IC7767 IC8080 IC8084 IC201 IC7762 IC7751 IC217 IC6319 IC8088 IC226 IC6340 IC8095 IC308 IC234 IC8086 IC6321 IC8102 IC6346 IC7757 IC313 IC7760 IC8093 IC8094 IC220 IC7755 IC6344 IC10100 IC7765 IC10098 IC8087 IC7759 IC10093 IC6313 IC310 IC229_2 IC253 IC6314 IC10092 IC8089 IC7756 IC7763 IC311 IC10096 IC230 IC225 IC8096 IC8101 IC301 IC6342 IC6320 IC8099 IC10101 IC7764 IC7750 IC8081 IC7754 IC222 IC8083 IC6317 IC243 IC7769 IC6345 IC6347 IC278 IC10099 IC8097 IC221 IC244 IC7768 IC8085

**Figure S8. Phylogenetic congruence of each chromosome and the mitochondrial genome for isolates sampled 1-year after biocontrol application**. In the total evidence display tree colors in grids represent node bipartitions that were supported at a bootstrap support value ≥70% (black color), <70% (white color), and missing or inapplicable (grey color). Phylogenetic incongruency was represented as high conflict (red color) and low conflict (cyan color). Strain names are highlighted to show their lineage membership in the total evidence tree; the mitochondrial genome has insufficient variation and poor resolution of lineage structure. The red arrows track the position of strain IC15721 sampled from IN which belongs to lineage IB in the total evidence tree but sharing recent common ancestry with strains in both lineages across most chromosomes making lineage assignment difficult. The black arrows track the position of the AF36 biocontrol strain, IC1179, which belongs to lineage IC and has low phylogenetic conflict with other stains in that lineage suggesting a history of recombination.

### Post 1-year

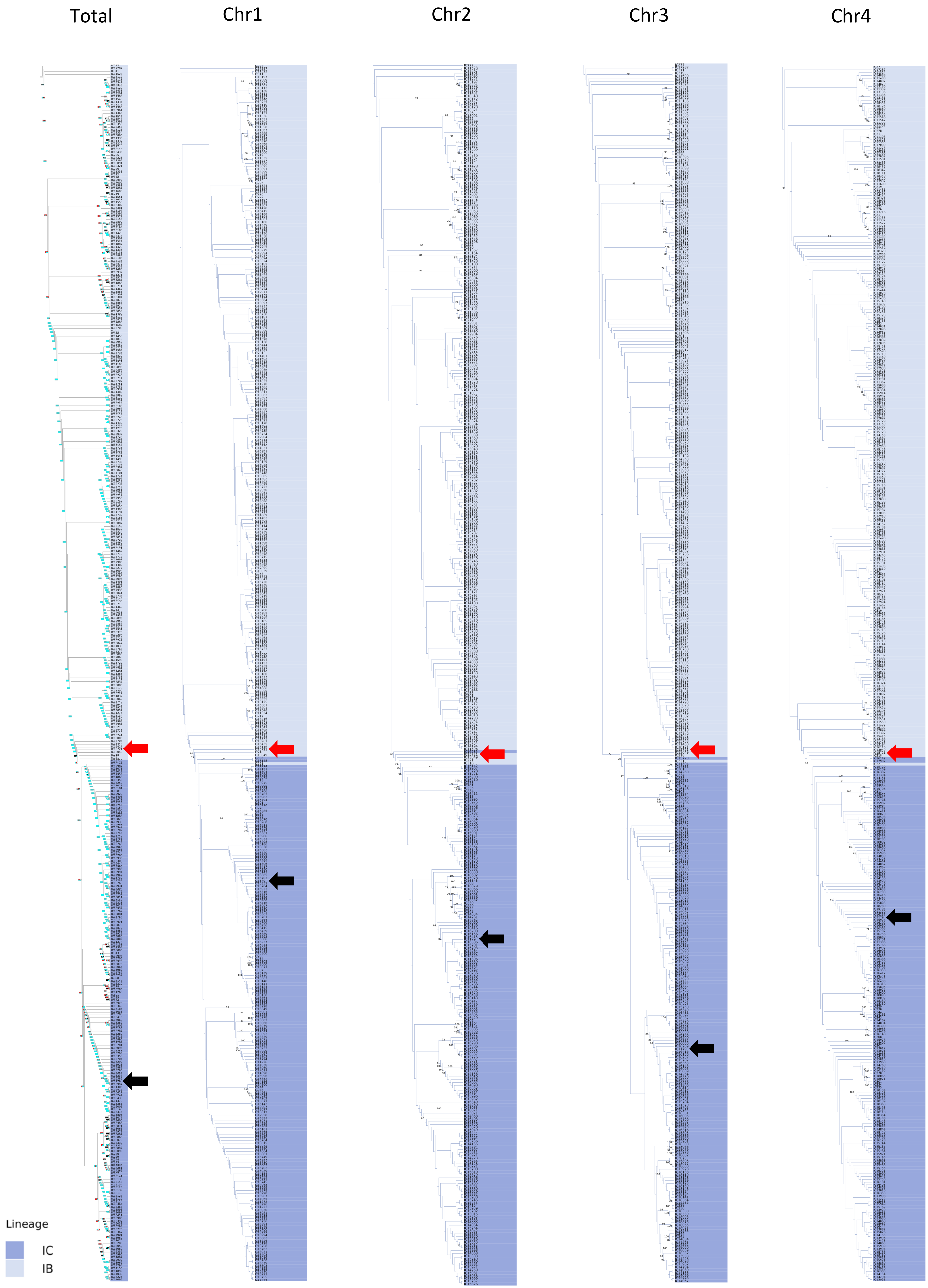

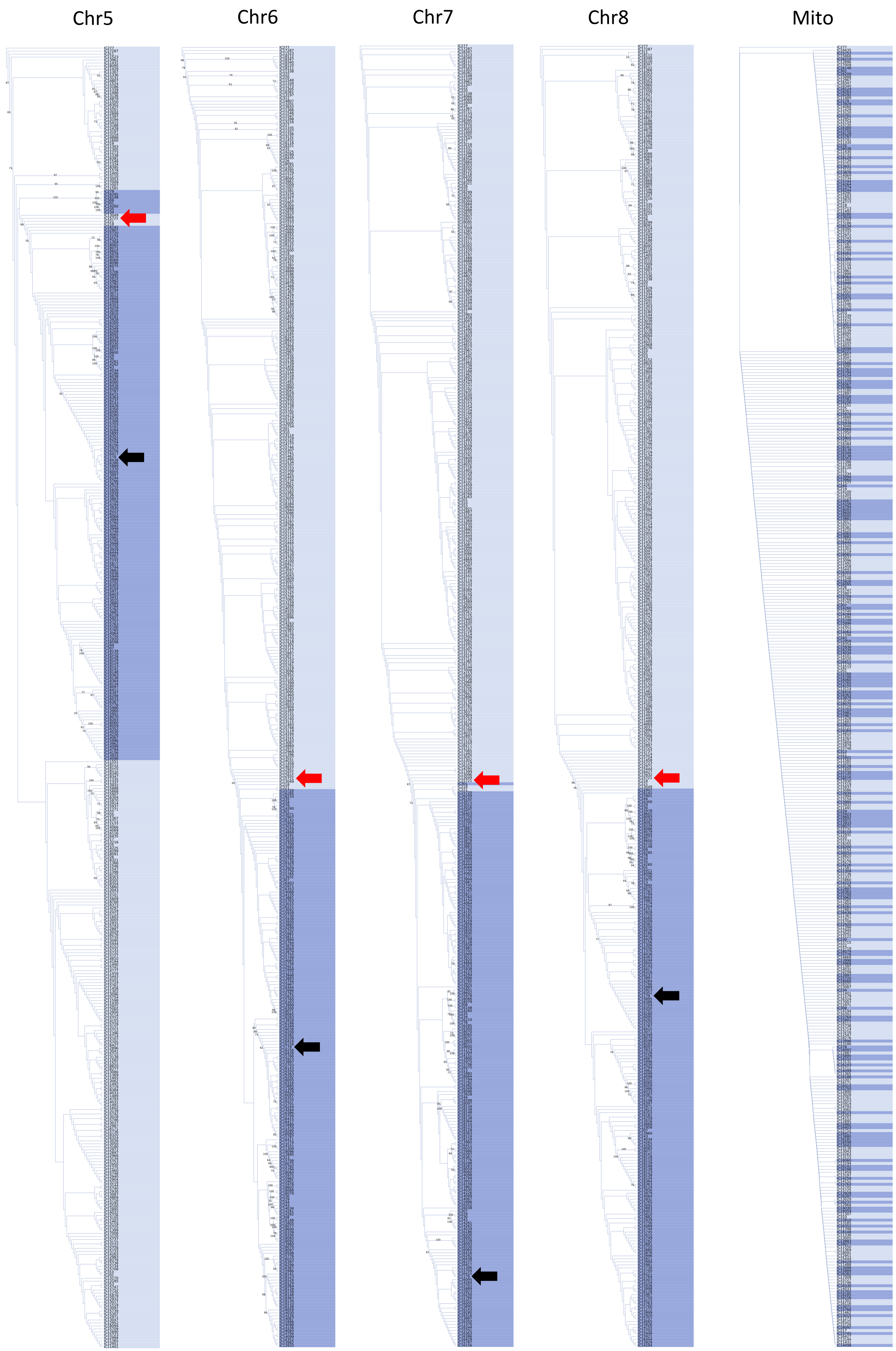
Chr5
Chr6
Chr7
Chr8
Mito

**Figure S9. Phylogenetic congruence of each chromosome and the mitochondrial genome for isolates from commercial TX cornfields sampled 3-years after application of the Afla-Guard biocontrol strain**. In the total evidence display tree colors in grids represent node bipartitions that were supported at a bootstrap support value ≥70% (black color), <70% (white color), and missing or inapplicable (grey color).  Phylogenetic incongruency was represented as high conflict (red color) and low conflict (cyan color).  Strain names are highlighted to show their lineage membership in the total evidence tree; the mitochondrial genome has insufficient variation and poor resolution of lineage structure. The red arrows track the position of strain IC14733 which belongs to lineage IB and shares very recent common ancestry with IC14744 and IC14769 with strong bootstrap support in chromosomes 1, 3, 5, 6 and 8; however, IC14733, IC14744 and IC14769 are also grouping with IC14720 with strong bootstrap support in chromosomes 2, 4, and 7.

### Post 3-years

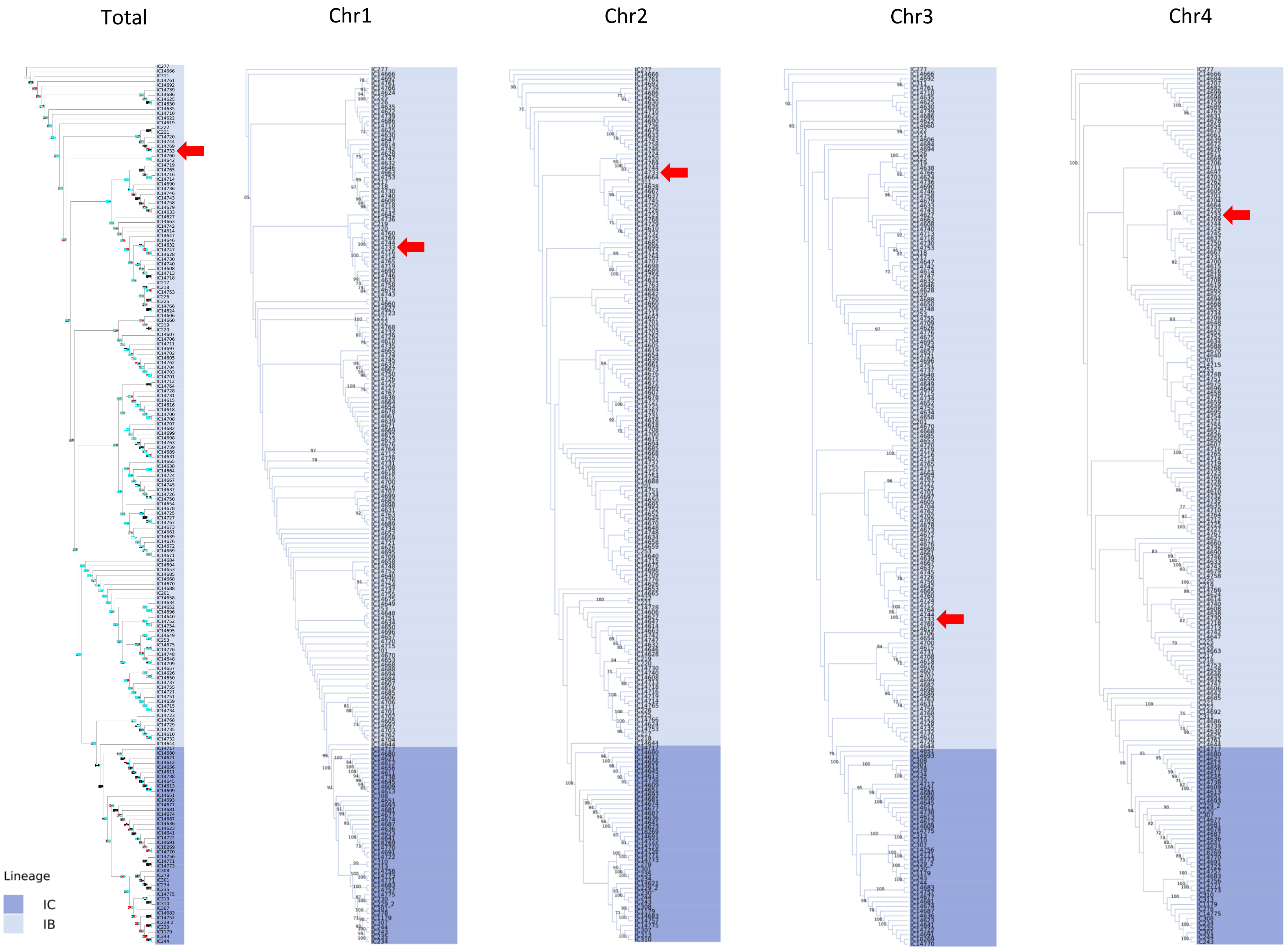

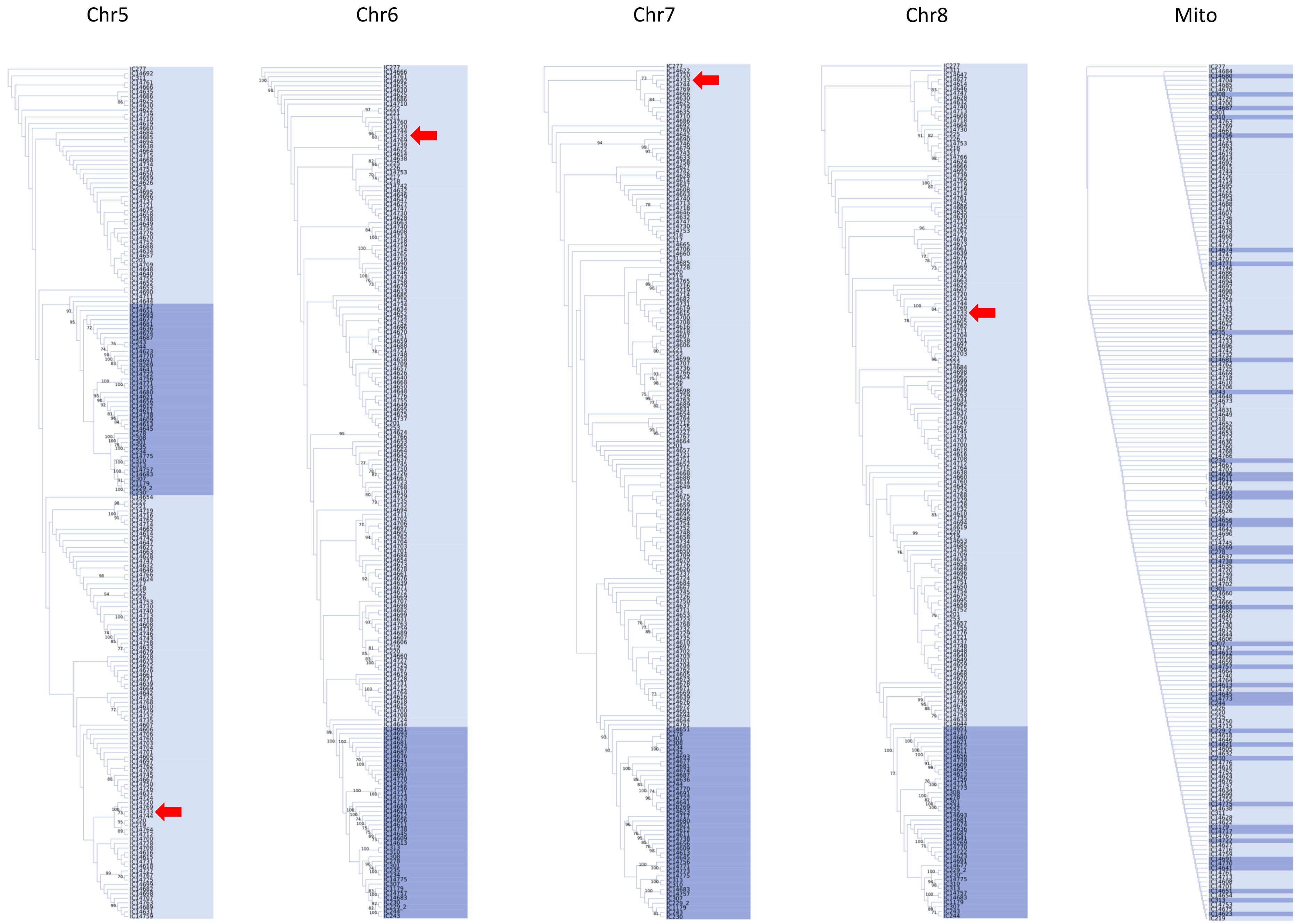
Chr5
Chr6
Chr7
Chr8
Mito

**Figure S10. Chromosomal LD plots for *A. flavus* lineages IB and IC.** LD plots are displayed using Haploview for each chromosome across four different sampling time points for lineages IB and IC. In each LD plot, black lines outline the edge of the spine of strong LD. In the coloring scheme, red represents strong LD (LOD ≥ 2, D' = 1), shades of pink/red represent intermediate LD (LOD ≥ 2, D' < 1), blue represents weak LD (LOD < 2, D' = 1) and white represents no LD (LOD < 2, D’ < 1).

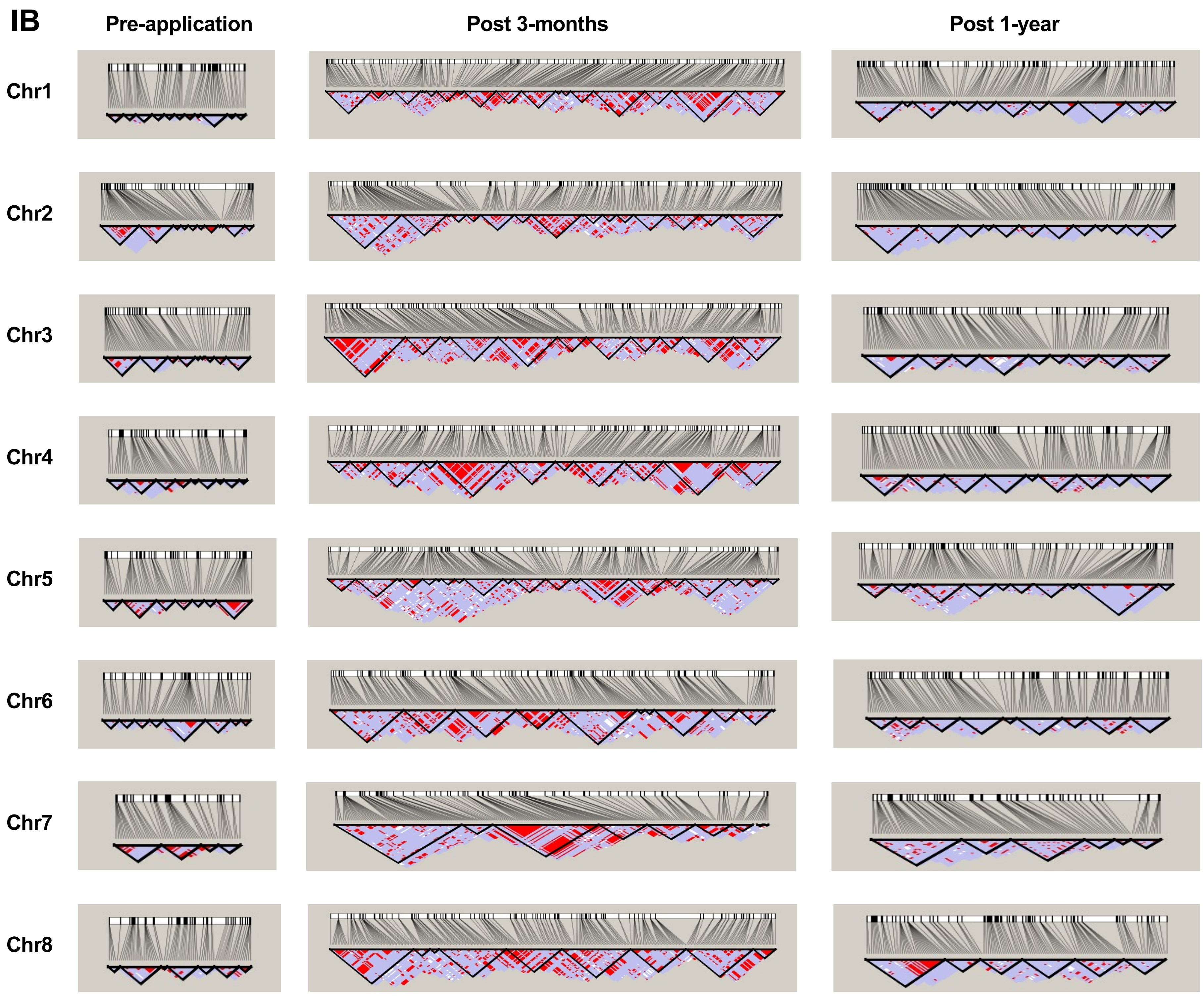
IB
Pre-application
Post 3-months
Post 1-year
Chr1
Chr2
Chr3
Chr4
Chr5
Chr6
Chr7
Chr8

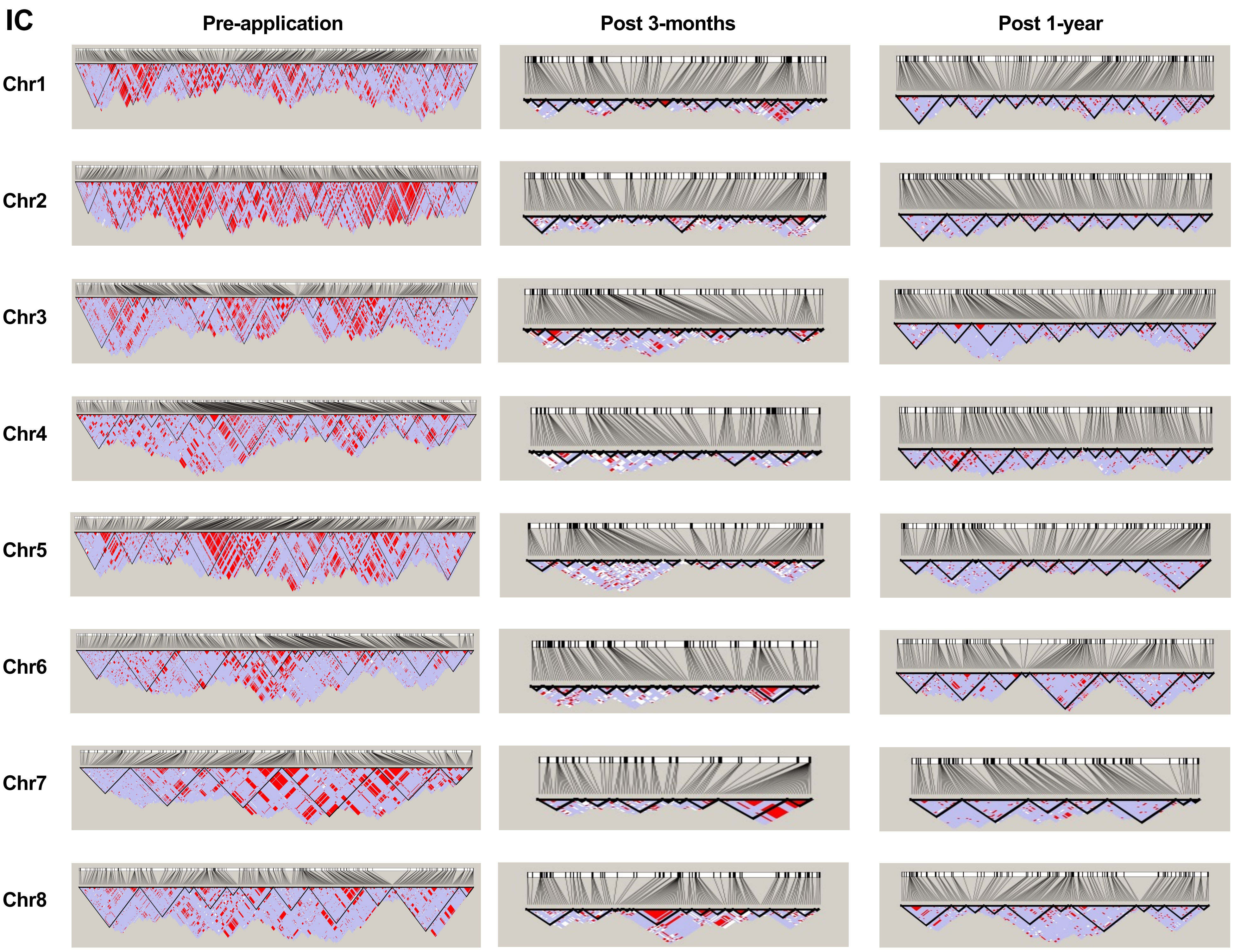
IC
Pre-application
Post 3-months
Post 1-year
Chr1
Chr2
Chr3
Chr4
Chr5
Chr6
Chr7
Chr8

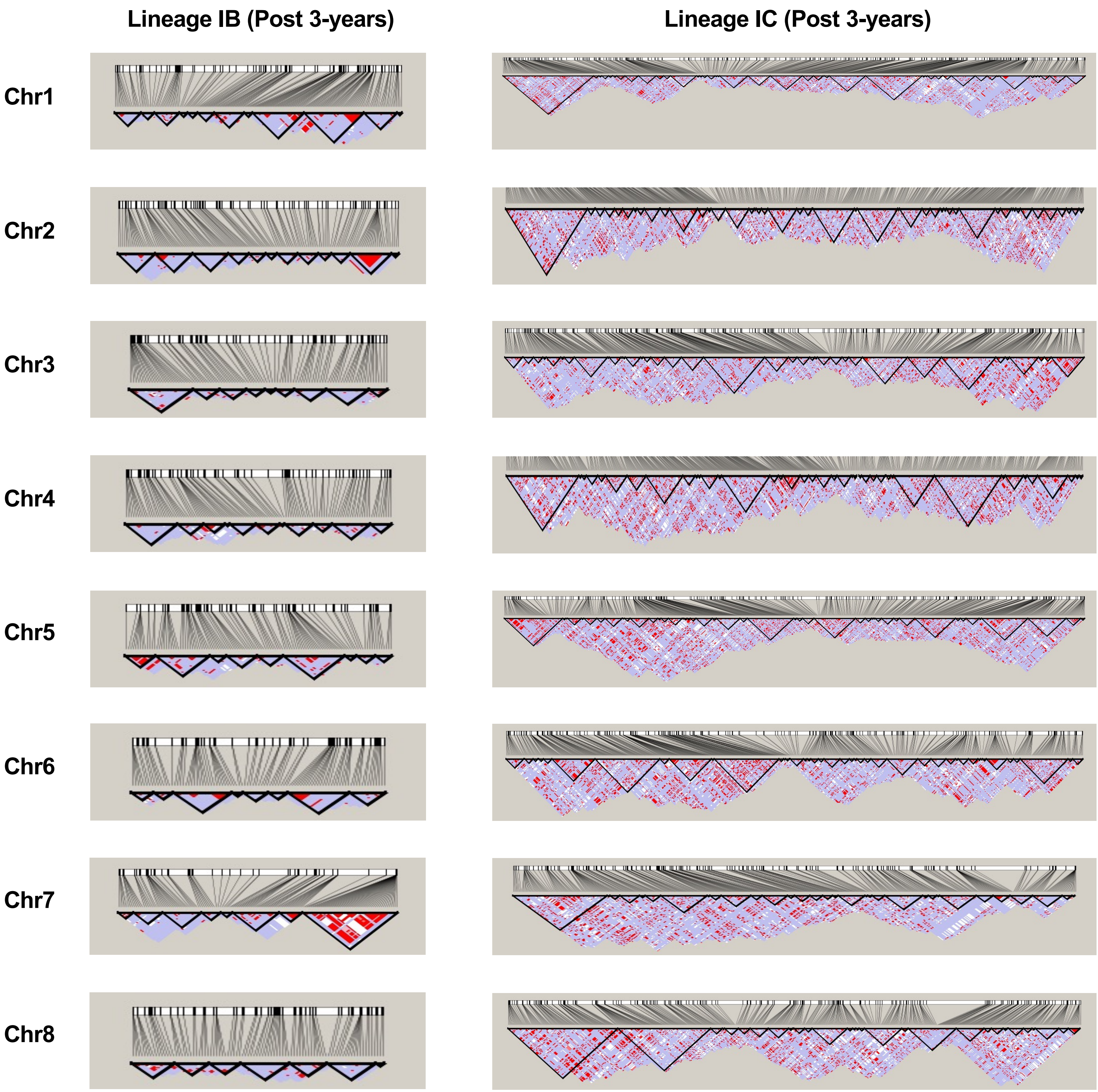
Lineage IB (Post 3-years)
Lineage IC (Post 3-years)
Chr1
Chr2
Chr3
Chr4
Chr5
Chr6
Chr7
Chr8
